## Supplementary Information for "Overt and covert paths for sound in the auditory system of mammals"

#### Si 01 About Material and Methods

#### Si 01A Aim of the study

This study was designed to test the hypothesis that the collagen fibers of the tympanum are piezoelectric when subjected to acoustic vibrations. Our protocol involved stimulating the tympanum at various frequencies and using various acoustical pressures. We measured the potentials resulting from this stimulation to determine if their electric properties were dependent upon the amplitude (dB SPL) and the acoustic frequency (Hz).

#### Si 01B Types of Collagen

Collagens II and I are piezoelectric histological components of eardrum. Their properties are very similar and we made measurements not only on collagen II of eardrum, but also on collagen I of the patellar tendon with the purpose to know at best their piezoelectric properties.

It is possible to detect an electric potential isochronous to the acoustic vibration between an indeterminate point of the tympanum and the mastoid bone^S^^[1]^. It does not follow necessarily, however, that the potential measured in this type of experiment is produced by the Outer Hair Cells (OHCs). Our methodology^S^^[2]^ allows us to demonstrate, in vivo, and under normal physiological conditions the piezoelectricity both of collagen I in tendons and of collagen II in eardrums.

#### Si 01C Piezoelectric proprieties of the fibrous collagen

The piezoelectric tensor of collagen I has a symmetry close to the hexagonal crystal structure^S113^. Its fibers are ferroelectric and piezoelectric ^S114,^ ^S115^. Stimulating these fibers by means of high frequency sounds directly affects osteogenic cells^S116^-^S117^. In a similar way, the production of the collagenous fibres of the tympanic membrane (TM) is increased and modeled by acoustic stimulations: In vitro, applied mechanical forces are able to promote TM-fibroblastic differentiation, increasing the production of collagen II, that is a peculiarity of TM structure^S118^.

A detailed analysis of the Piezoresponse Force Microscopy signal of collagen I “…revealed clear shear piezoelectric activity associated with piezoelectric deformation along the fibril axis.” (Harnagea, 2010). Piezoelectric activity of collagen fibrils can be detected in vitro in a large range of frequencies going from a few

Hz^S119^ up to more than 200 kHz^S120^ up to 1 MHz^S121^. This result corresponds to the outcomes of several studies

with respect to collagen in vivoS122 - S123 (Harnagea, 2010). The inverse piezoelectric effect is also

demonstrableS124***.*** (Harnagea, 2010). The properties of collagen I are thought to be similar to properties of

collagen IIS125 - S126 -S127 (Minary-Jolandan, 2009) . To confirm this, we have measured synchronous electrical

potentials not only on eardrum collagen radial fibres but also on the patellar ligaments of individuals at various ages. The amplitude of measured synchronous potentials increases with the frequency of the sound signal.

In order to verify the piezoelectric activity of the tympanum, an electrode is placed on the manubrium, another on the periphery according to the straight lines either A or G. The "standard" measurements we did, and studied statistically, were between an undetermined point of the posterior side of the manubrium and an undetermined point of the posterior limbus.

In order to evaluate the electrical behaviour of points belonging to the central structure (manubrium) during

acoustic stimulations, electrodes can be placed at two points on the same side of the manubrium (symbolized by an F letter in **b**). Since the collagen fibres have their positive end linked to the malleus, the system should then measure an electrical potential close to zero. On the contrary, if the electrodes would be placed facing each other on either side of the manubrium (E), the system should allow us to capture the activity of a bundle of circular fibres, ie an potential isochronous to the acoustic waves. Isochronous electrical responses between two points on the same side (inner or outer) of the annulus tympanicus (HH’) are generally impossible to measure. Regarding exceptions, it might be that electrodes had been positioned, either not on the same side of the annulus tympanicus, or on an inner circumferential collagen fibre. An analogous difficulty could be found along the manubrium of the malleus; yet it is easier to position the electrodes at one boundary of the manubrium.

#### Si 01D What about the myringoplasty ?

The most successful myringoplasty provides a happy outcome for the patient who becomes able to listen to intelligible conversations: yet regarding frequencies above 2 kHz, it is limited to a poor hearingS128 - S129. Tympanic Membrane Perforation (TMP) causes hearing loss which increases with the perforation sizeS130 - S131. TMP hearing loss generally increases as the frequency increasesS132. Audiometric results shows that the larger the TMP, the larger the air–bone gap conductive hearing lossS133. Patients, with a small central perforation, recover with just a conservative management, but 10–15% suffer from bone conduction lossS134.

Myringoplasty is most successful if the graft is made up of type II collagen (cartilage) and if the perforation is centralS135 - S136. Whatever the case may be, a local restructured collagen II must colonize the graft before the

graft can return to its full functionality. This means that fibres should recover their previous structure and their peculiarities, including their radial disposal according to an homogenous center-periphery polarity. Very commonly, the transplant regardless of its origin and its histological nature (temporal fascia, cartilage, etc) does not perfectly restore radial or circumferential structures *ad integrum* even after several months (or years).

As a result, the hearing is restored only for low and medium frequencies in such a way that the subject permanently loses the perception of the high frequencies. This finding supports our hypothesis : A bio-electronical pathway comes from collagen fibres of the eardrum. If their geometry loses its radial appropriate design, the pT becomes ineffective, and high frequencies are poorly perceived

##### Si 01E Measurements of piezoelectric response on eardrums

For measurements on eardrums we use a lock-in amplifier to drive a loudspeaker. In this manner, we broadcast a sinusoidal sound at about one meter from the external auditory conduit. We position a probe consisting of two electrodes at the centre and the periphery of the tympanum. This probe captures the piezoelectric response of the radial tympanic fibres when they vibrate in response to the sound sent to the tympanum. The lock-in amplifier makes it possible to select only those electrical responses isochronous to the acoustic stimulation. We measure electrical responses to stimulations at different acoustic frequency levels. See fig. below :position of the electrodes of the probe.

A Lock In Amplifier, via an electric wave of V_hp_ tension (output) creates an acoustic wave at various frequencies diffused by a loudspeaker.

##### Si 01G Measurement Electrodes for the eardrums

For the measurements on the eardrums, we tried various configurations:

• electrodes attached to a plastic bracket, ensuring a constant distance

• two electrodes easy to handling, more easily affixed to appropriate parts of the eardrum, but not allowing standardization of the inter-electrodes distance.

The shielding braid is put in contact with the peripheral structure of the tympanum, away from the handle of the malleus. The central copper wire is put in contact with the central structure of the tympanum: either umbo (direction “A”) or handle of the malleus (“G”).

**Position of the electrodes of the probe**

It is possible to detect an electric potential isochronous to the acoustic vibration between an indeterminate point of the eardrum and the mastoid bone**^[3]^**.. However, it does not necessarily follow that this potential be a microphonic produced by Outer Hair Cells OHCs. On the contrary, our methodology allows us to demonstrate, in vivo, and under normal physiological conditions, that it is the result of the piezo-electricity of the collagen II of the eardrums.

| (a) | (b) |
| --- | --- |

| 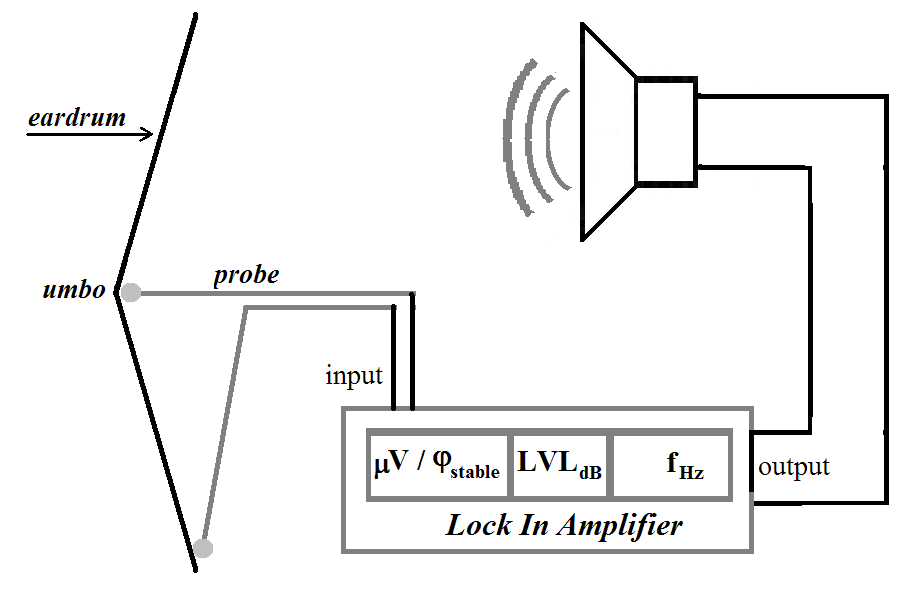 | 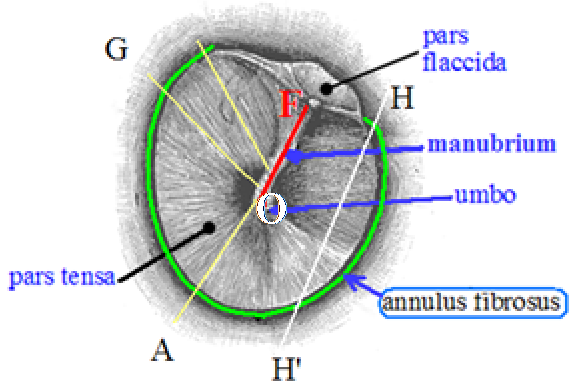 |
| --- | --- |
| **a** | **b** |

**Figure S01**

The closer together the electrodes are placed, the smaller the area recorded will be (i.e. local recordings). This is one advantage of the differential recording technique^[4]^. This "differential technique" is necessary for determining the source of a potential***^s^***^[5]^.

##### Si 01F Measurements of piezoelectric response on knees (or other tendons)

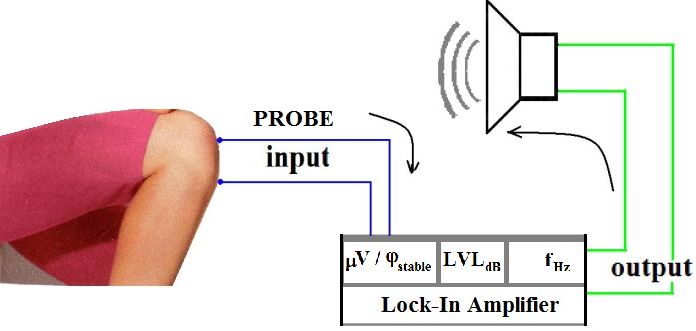

Figure S02
Diagram of the design used to measure the piezoelectric potential from the knee tendon.
V indicates the isochronous voltage obtained when the phase j is stabilized, in response to the sound wave of frequency f (Hz) with a LVL (dBSP) amplitude.

###### **Measurement Electrodes for tendons**

For measurements on the **mastoid** area, the patellar ligament or other tendons, we used « single use » *pregelled ECG-electrodes (Ref. “Comepa” 3.02.0200); they were connected to the* LockIn Amplifier by two *medical leadwires* AIP018 with *Touch proof connectors* DIN42802. (Table S09)

**Effect of the length of the collagen fibres**

To evaluate the effect of the length of the collagen fibre bundles on their piezoelectric response, we performed some experiments on the most accessible bundles (patellar ligament). This allowed us to understand in particular why the response of these bundles was much larger than those of the eardrum.

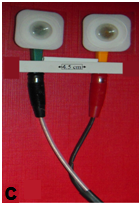

**(c)** Probe used for measurement on the **knees**

As the amplitude of the voltage generated (in µV) increases with the distance (in mm) between electrodes; we, thus, ensured a fixed distance (45 mm) between the centre of the two COMEPA, 3020200 CA electrodes.
**Figure S03**

| d (mm) | V (μV) |
| --- | --- |
| 40 | 12.1 |
| 45 | 15.1 |
| 50 | 18.0 |
| 55 | 22.8 |
| 60 | 28.2 |
| 65 | 30.0 |

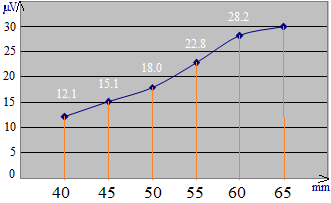

**Figure S04**

**Measurements on knee** : Acoustical frequency of the stimulus was 4000 Hz
x axis (d): distance (mm); y-axis : voltage (µV).
Result : **V (µV) = 0.76 d (mm) -19 mm ; R² = 0.98**

We have taken measures with different distances of separation between different electrodes placed in contact with the skin of a subject. The table, schema, and formula above present the effect of the distance between the electrodes. The voltage is proportional to the distance, i.e. to the length of the piezoelectric fibres.

In order to evaluate the electrical behaviour of points belonging to the central structure (manubrium) during acoustic stimulations, electrodes can be placed at two points on the same side of the manubrium (symbolized by an F letter). This system can detect whether there is an electrical isochronism between these points (synchronous potential close to zero). On the contrary, electrodes might be placed facing each other on either side of the manubrium (E). This latter system would allow us to capture the activity of a bundle of circular fibres (not completed).

The following letters were added by us : A or G: “radii” types of fibres of collagen ; HH’: arbitrary cord joining two peripheral points ; F: manubrium of the malleus ; GAH’H: annulus tympanicus. In order to verify the piezoelectric activity of the tympanum, an electrode is placed on the manubrium, another on the periphery according to the straight lines either A or G.

With humans the synchronous electrical responses between two points on the same side (inner or outer) of the annulus tympanicus (HH’) are generally impossible to measure (Table S10). Regarding the E48 RE (HH’) exception, it might be that electrodes had been positioned, either not on the same side of the annulus tympanicus, or on an inner circumferential collagen fibre

#### Si 02 The "space shift" problem

**Si 02A The "space shift" problem**

According to F. Mammano***^S^***^[6]^, and others, the tympanic potentials that we measure being low (less than one mV), they would be unable intervene in the operation of the OHCs. But, as we explained in the main part of our work, the operation of the TkS suppresses the need for strong potentials.

Yet Harnagea has pointed out that the pT potentials measured “in vitro” between the two extremities of a single collagen fibre can reach, and even exceed, 10 mV. If this is also the case “in vivo”, such potentials could reach the cochlea (OHCs) with sufficient amplitude, provided that the fibres share a homogenous direction and polarity^S^^[7]^*”.* Other authors explain that the mixed polarity of the collagen fibres in the biological structures significantly weakens their piezoelectric response^S^^[8]^. With respect to the eardrum, we took measurements by simple and soft application of the electrodes to the tympanic epidermis which is, more or less, insulating, in such a way that there was a weakening on the measured signal with respect to the genuine signal. So, we must be aware of bias caused by our measurements in vivo: We were not able to set up the two electrodes of the probe with the required precision in such a way that they would be in contact with both ends of a single fibre or bundle.

Often, there will be a mismatch between the ends of a seemingly united bundle of fibres: Let us assume that a first electrode is near the upper end of a fibres bundle and the other electrode near the lower end of the same fibres bundle; if the first electrode is located on the left of the first extremity, and if the second electrode is located on the right of the other extremity, the electrical pathway between the two electrodes does not correspond to the targeting of a single fibre: There is a space shift with implications for the conductivity and a measured voltage attenuation.

We tried to detect on the patellar ligament of knees, whether a configuration of the electrodes at the two ends of a supposedly unique collagen bundle could give higher measures than if electrodes were supposedly mismatched with the same bundle: one electrode on the external top side end of the "bundle", and the other electrode on the internal bottom side.

Our Protocol of measurement on knees was the following. We placed the subject at a fixed distance from the loudspeaker. We, then, apposed one electrode at the apex (lower point) of the patella and another electrode at the upper anterior top of the tibia. We made two trials on each knee for each of two frequencies. One trial used electrical stimulation of 70 dB SPL and the other of 80 dB SPL. As Harnagea pointed out, a maximum potential is collected if the electrodes are apposed at both ends of the same fibre, but this potential decreases or cancels if fibres overlap in all directions, as do the fibres of a "felt".

In vivo, the design of the collagen fibres is ordered by the functional necessities of the biological tissue. From a macroscopic point of view, we find very ordered structures, at the level of bones, joints, tendons, muscles or eardrum. For example, it is well known that the bone tissue is shaped according to the "law" of Wolf***^S^***^[9]^.

The placement of the electrodes on the two ends of a single fibre or a single bundle of collagen fibres is expected to get a larger potential than if both electrodes are at the extremities of two different bundles of fibres. That is much easier to be successful if measurement is done on the patellar tendon, than on the eardrum.

We tried to measure, as locally as possible, voltage variations, synchronous to sound pressures. This is achievable without too much difficulty for collagen fibres *in vitro*. In our approach of the eardrum *in vivo*, the line that goes from a central point of contact to a peripheral one is laid out so that it corresponds as closely as possible to a bundle of radial fibres. But it is not possible for us to lay out with certainty both electrodes of the probe at the two ends of the same bundle of collagen fibres; It is likely that our measures may quite often involve two more or less neighbouring bundles; For this reason, the measurement will result in lower levels than if it were taken on one and the same bundle. This practical difficulty could help us understand why potentials measured on the eardrum are weaker than those that can be measured on the knees.

##### Si 02B Effect of space-shifts on measurements of collagen piezoelectricity

The Supplementary Table1 gives the results of our measurements for two subjects (B15, E48), on their knees (right and left) for different frequencies (f in Hz). We set the potential (Vout), sent by the LockIn to the Harman Speaker to 5V; the potential measured by the probe placed on the knee is expressed in µV.

| **Sujets** | **B15** | **B15** | **B15** | **B15** | **B15** | **B15** | **E48** | **E48** | **E48** | **E48** | **E48** | **E48** |
| --- | --- | --- | --- | --- | --- | --- | --- | --- | --- | --- | --- | --- |
| **Knee (R / L)** | Right | Right | Right | Left | Left | Left | Right | Right | Right | Left | Left | Left |
| **top electrode** | med. | ext. s. | int. s. | med. | ext. s. | int. s. | med. | ext. s. | int. s. | med. | ext. s. | int. s. |
| **bottom electrode** | med. | int. s. | ext. s. | med. | int. s. | ext. s. | med. | int. s. | ext. s. | med. | int. s. | ext. s. |
| **Line** | {1} | {2} | {3} | {1} | {2} | {3} | {1} | {2} | {3} | {1} | {2} | {3} |
| **f (Hz)** | D1 | D2 | D3 | G1 | G2 | G3 | D1 | D2 | D3 | G1 | G2 | G3 |
| **500** | 30 | 25,3 | 8 | 29 | 24,1 | 16,6 | 12,5 | 33 | 20 | 28,2 | 5 | 12,3 |
| **1000** | 21,1 | 26,5 | 7,7 | 89,3 | 31,1 | 16,9 | 18,1 | 30,5 | 23,5 | 33,2 | 21,1 | 17,2 |
| **2000** | 70 | 46,5 | 9,7 | 110,2 | 41,1 | 21,3 | 29,6 | 41,5 | 28,5 | 41,1 | 28,5 | 33,1 |
| **3000** | 98,5 | 57,9 | 10 | 111,5 | 50,1 | 25 | 66 | 58,2 | 40 | 78,3 | 60,6 | 39,2 |
| **4000** | 110 | 66,6 | 18,4 | 152 | 58,2 | 26,8 | 100,5 | 73 | 41,1 | 111 | 87,7 | 44 |
| **6000** | 112,3 | 87,5 | 22,4 | 163,5 | 73 | 33 | 128,7 | 99,5 | 42,5 | 144,2 | 110 | 51 |
| **8000** | 173 | 110 | 28 | 184,3 | 88,7 | 38,5 | 180 | 105,6 | 43,7 | 201,5 | 135,8 | 53,3 |
| **9000** | 185,1 | 119 | 34,5 | 195,7 | 93,9 | 43,2 | 239,1 | 156 | 48,5 | 244,2 | 177 | 60 |
| **10000** | 215,2 | 135 | 44,3 | 185,9 | 102 | 47 | 256,7 | 200,2 | 51 | 264,1 | 189,4 | 71,1 |
| **16000** | 218,9 | 209,9 | 58,3 | 219,1 | 145 | 67,3 | 321,6 | 205 | 47,5 | 305 | 211 | 72,2 |
| **20000** | 282,6 | 270,5 | 70,8 | 381,4 | 160 | 79,9 | 450 | 233,1 | 52,7 | 412 | 258 | 79,5 |
| **25000** | 390,8 | 288,5 | 73 | 423,7 | 182 | 96,6 | 500 | 256,2 | 61 | 523 | 289,2 | 85 |
| **Mean** | 159 | 120 | 32 | 187 | 87 | 43 | 192 | 124 | 42 | 199 | 131 | 52 |
| **Median** | 143 | 99 | 25 | 174 | 81 | 36 | 154 | 103 | 43 | 173 | 123 | 52 |
| **Supplementary Table 1**  *med*. means "median"; *int.* means "internal"; *ext*. means "external"; *s*. means "side" | | | | | | | | | | | | |

The Supplementary Table 2 give the results of our measurements for two subjects (B15, E48) on their two knees (right and left) for different frequencies (f in Hz), on the central line *{1}*.

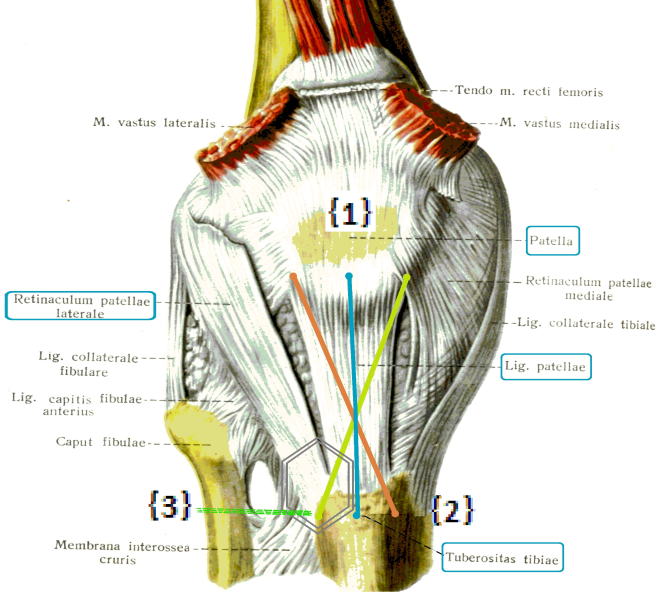

**(a)**

**(b)**

|  | {1} | {2} | {3} |
| --- | --- | --- | --- |
| 500 | 28.6 | 24.7 | 14.45 |
| 1000 | 27.15 | 28.5 | 17.05 |
| 2000 | 55.55 | 41.3 | 24.9 |
| 3000 | 88.4 | 58.05 | 32.1 |
| 4000 | 110.5 | 69.8 | 33.95 |
| 6000 | 136.45 | 93.5 | 37.75 |
| 8000 | 182.15 | 107.8 | 41.1 |
| 9000 | 217.4 | 137.5 | 45.85 |
| 10000 | 235.95 | 162.2 | 49 |
| 16000 | 262.05 | 207.45 | 62.8 |
| 20000 | 396.7 | 245.55 | 75.15 |
| 25000 | 461.85 | 272.35 | 79 |

Figure S05a and S05b

Experimental mimicry of offset on fibres of a patellar ligament: We used voluntarily three configurations for comparisons between a good precision {1} versus bad ones {2 & 3} (The knee anatomy figure is excerpted from Sinelnikov^S^^[10]^).

| **F (Hz)** | **{1 }** | **{2 }** | **{3 }** |
| --- | --- | --- | --- |
| **500** | 28,60 | 24,70 | 14,45 |
| **1000** | 27,15 | 28,50 | 17,05 |
| **2000** | 55.55 | 41,30 | 24,90 |
| **3000** | 88,40 | 58,05 | 32,10 |
| **4000** | 110,50 | 69,80 | 33,95 |
| **6000** | 136,45 | 93,50 | 37,75 |
| **8000** | 182,15 | 107,80 | 41,10 |
| **9000** | 217,40 | 137,50 | 45,85 |
| **10000** | 235,95 | 162,20 | 49,00 |
| **16000** | 262,05 | 207,45 | 62,80 |
| **20000** | 396,70 | 245,55 | 75,15 |
| **25000** | 461,85 | 272,35 | 79,00 |

Table 02

Ideally, measurements on collagen should concern a single fibre. But this is clinically very difficult: There may be an offset illustrated above (**a:** left panel), which we have minimized as much as possible. To check the effects of this type of configuration, we carried out measurements voluntarily strongly shifted on the tendon of a knee (**b**).
 * The musculo-tendinous bundle concerned by the measurement line {1} includes the central collagen fibres of the tibial ligament.
 ** The musculo-tendinous bundle concerned by the measurement line {2} goes from the upper lateral portion of the tibial ligament down to its medial lowest part.
 *** The musculo-tendinous bundle concerned by the measurement line {3} goes from the upper retinaculum patellae down to the membrana interossea cruris (hexagonal framing at the bottom of {3}).

Harnagea showed in vitro that the response is low or non-existent if the electrodes are arranged in any two points of the culture, but this response reached tens of mV if the electrodes are placed at both ends of a **single fibre (**Denning et al. 2017)^[11]^. The longitudinal piezoelectric coefficient for idividual fibrils at the nanoscale was found to be roughly an order of magnitude greater than that reported for macroscopic measurements of tendon, the low response of which stems probably from the presence of poorly oriented fibrils. Thus we can assume that the pT voltages are in the mV range and may reach the DOHC complex.

Our assumption, based on the reflections of Harnagea, had made us envisage a shift between the central curve on the one hand {1}, and the two shifted curves on the other hand. We supposed that the curves {2} and {3} would be of amplitude and/or slope lower than the curve of reference {1}, which verified by our measures.

Moreover, it appears that the curve of configuration {3}(hexagonal framing at the bottom of {3}) is definitely lower than the two other curves; this is coherent with the fact that the measurement line {3} encompasses two musculo-tendinous bundles, which fact logically dampens the signal collected on this level compared to the signals collected in {1} or {2}.

| F (Hz) | *D1* | *G1* | *D1* | *G1* |  |  |  |
| --- | --- | --- | --- | --- | --- | --- | --- |
|  | Sujet B15 | Sujet E48 | *median {1}* | *mean {1}* | *sd {1}* | | |
| 500 | 30 | 29 | 13 | 28 | 29 | 25 | 8 |
| 1000 | 21 | 89 | 18 | 33 | 27 | 40 | 33 |
| 2000 | 70 | 110 | 30 | 41 | 56 | 63 | 36 |
| 3000 | 99 | 112 | 66 | 78 | 88 | 89 | 20 |
| 4000 | 110 | 152 | 101 | 111 | 111 | 118 | 23 |
| 6000 | 112 | 164 | 129 | 144 | 137 | 137 | 22 |
| 8000 | 173 | 184 | 180 | 202 | 182 | 185 | 12 |
| 9000 | 185 | 196 | 239 | 244 | 217 | 216 | 30 |
| 10000 | 215 | 186 | 257 | 264 | 236 | 231 | 37 |
| 16000 | 219 | 219 | 322 | 305 | 262 | 266 | 55 |
| 20000 | 283 | 381 | 450 | 412 | 397 | 382 | 72 |
| 25000 | 391 | 424 | 500 | 523 | 462 | 459 | 62 |
| **Supplementary Table 3** Electrodes on the median line between the tip of the patella and the top of the tibia: *Configuration {1}* | | | | | | | |

###### **Si 02Ba The "space shift" (configuration 2 & 3):**

The Supplementary Table 3, give the results of our measurements for two subjects (B15, E48), on their two knees (right and left) for various frequencies F (Hz), on the line of configuration {2}.

| f (Hz) | *D2* | *G2* | *D2* | *G2* | *median {2}* | mean {2} | sd {2} |
| --- | --- | --- | --- | --- | --- | --- | --- |
|  | Sujet B15 | Sujet E48 |  |  |  | | |
| 500 | 25 | 24 | 33 | 5 | 25 | 22 | 12 |
| 1000 | 27 | 31 | 31 | 21 | 29 | 27 | 5 |
| 2000 | 47 | 41 | 42 | 29 | 41 | 39 | 8 |
| 3000 | 58 | 50 | 58 | 61 | 58 | 57 | 5 |
| 4000 | 67 | 58 | 73 | 88 | 70 | 71 | 13 |
| 6000 | 88 | 73 | 100 | 110 | 93 | 93 | 16 |
| 8000 | 110 | 89 | 106 | 136 | 108 | 110 | 20 |
| 9000 | 119 | 94 | 156 | 177 | 138 | 137 | 37 |
| 10000 | 135 | 102 | 200 | 189 | 162 | 157 | 46 |
| 16000 | 210 | 145 | 205 | 211 | 208 | 193 | 32 |
| 20000 | 271 | 160 | 233 | 258 | 246 | 230 | 49 |
| 25000 | 289 | 182 | 256 | 289 | 272 | 254 | 50 |
| **Supplementary Table 4**  One electrode shifted up on the outer side of the patella and the other electrode staggered downstairs on the internal side of the top of the tibia: *Configuration {2}* | | | | | | | |

The table Supplementary Table 4 below, give the results of our measurements for two subjects (B15, E48), on their two knees (right and left) for various frequencies *f* (Hz), on the line of configuration {3} (**green**).

| F (Hz) | *D3* | *G3* | *D3* | *G3* | *median {3}* | *mean {3}* | *sd {3}* |
| --- | --- | --- | --- | --- | --- | --- | --- |
|  | Sujet B15 | Sujet E48 |  |  |  | | |
| 500 | 8 | 17 | 20 | 12 | 15 | 14 | 5 |
| 1000 | 8 | 17 | 24 | 17 | 17 | 16 | 7 |
| 2000 | 10 | 21 | 29 | 33 | 25 | 23 | 10 |
| 3000 | 10 | 25 | 40 | 39 | 32 | 29 | 14 |
| 4000 | 18 | 27 | 41 | 44 | 34 | 33 | 12 |
| 6000 | 22 | 33 | 43 | 51 | 38 | 37 | 12 |
| 8000 | 28 | 39 | 44 | 53 | 41 | 41 | 11 |
| 9000 | 35 | 43 | 49 | 60 | 46 | 47 | 11 |
| 10000 | 44 | 47 | 51 | 71 | 49 | 53 | 12 |
| 16000 | 58 | 67 | 48 | 72 | 63 | 61 | 11 |
| 20000 | 71 | 80 | 53 | 80 | 75 | 71 | 13 |
| 25000 | 73 | 97 | 61 | 85 | 79 | 79 | 15 |
| **Supplementary Table 5** One electrode shifted up on the inner side of the patella and staggered  downstairs on the external side of the top of the tibia: *Configuration {3}* | | | | | | | |

###### **Si 02Bb The "space shift" : comparison of the three configurations**

We compare the three medians (Supplementary Table 5); “*a*” and “*b*” are the coefficients of the polynomials of the curves of linear regression of the three configurations with “*a*” parameter of slope, and “*b*” ordinate at the linear origin of the curves of regression:

| Parameters | Median {1} | Median {2} | Median {3} | Decreasing order |
| --- | --- | --- | --- | --- |
| a | 0,018 | 0,010 | 0,003 | {1} > {2} > {3} |
| b | 31,12 | 27,76 | 20,05 | {1} > {2} > {3} |
| Supplementary Table 6  Table of Linear regressions | | | | |

To compare the effect of the three configurations, we joined together the data belonging to each one (Supplementary Table 7).

| f (Hz) | {1} | {2} | {3} |
| --- | --- | --- | --- |
| 500 | 28,60 | 24,70 | 14,45 |
| 1000 | 27,15 | 28,50 | 17,05 |
| 2000 | 55,55 | 41,30 | 24,90 |
| 3000 | 88,40 | 58,05 | 32,10 |
| 4000 | 110,50 | 69,80 | 33,95 |
| 6000 | 136,45 | 93,50 | 37,75 |
| 8000 | 182,15 | 107,80 | 41,10 |
| 9000 | 217,40 | 137,50 | 45,85 |
| 10000 | 235,95 | 162,20 | 49,00 |
| 16000 | 262,05 | 207,45 | 62,80 |
| 20000 | 396,70 | 245,55 | 75,15 |
| 25000 | 461,85 | 272,35 | 79,00 |
| Supplementary Table 7 Median, depending on the frequency-test (f_Hz_) according to each of the three configurations. | | | |

Si 02C Application to fibers of the tympanum (Denning et al.)

 As a preliminary remark, when the electrodes are laid out at the two ends of the handle of the hammer and on the same side, it is expected that the measured voltage be null or so. We verified it on chinchilla and human.

Otherwise, we can apply the previous observation on knees about the measurement lines {1} , {2} and {3} to our measurements on eardrum fibres: To get a maximum voltage, the optimal design would be to put the electrodes at both ends of a same radial fibre of collagen. If there is a spatial shift, there will necessarily be a lessening of the amplitude of the signal. Not being able to meet that theoretical requirement, we have to accept that constraint. In order to decreases its statistical effect, we may calculate an individual parameter for each series of measures in a given configuration (see § statistics).

Denning et al. (2017)^[12]^ have shown that nanoscale fibrils can generate higher electrical potentials (mV) than long-tendon fibers (μV). As suggested by Harnagea they deduce that these long fibers have a polar orientation ratio of their constituent fibrils significantly different from 50/50%.

It is probable that, in the resting position, the polarity of the collagen radial fibers be oriented due to their centripetal development, resulting their positive end in the centre and their negative end in the periphery.

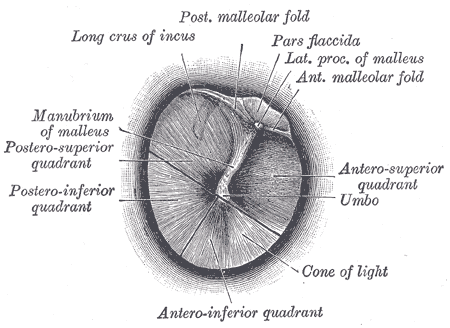

Figure S06

internal aspect of tympanic membrane

The radial fibres of collagen grow from the umbo to the periphery .

The ENT experimenter tries to put both of the probe electrodes at the ends of one single fibre. Yet such an utmost precision is unreachable because every fibres are extremely thin and not separately identifiable. So it happens an unwished offset on fibres of the tympanum.

#### Si 03 Variations

##### Si 03A Variation of eardrum tension

The subject on which we take the measurements is equipped with several types of very efficient biological control mechanisms, which are usually managed by reflex mechanisms (unconscious and involuntary activity), in a non-controllable way. For this reason, the acoustical strain of the eardrum (and by implication, its piezoelectric response) may vary in a non-controllable manner according to the expectations of the subject, his psycho-physiological state, the potential effect of local anaesthesia on the subject’s biological reactions.

Eardrum tension depends mainly on two types of muscle. The "tensor tympani" is a striated muscle, that some subjects may contract, voluntarily, on demand, for a fairly short period (^S^^[13]^-^S^^[14]^). Furthermore, for everybody, it contracts by reflex, involuntarily and unconsciously, when it is appropriate to attenuate or target the perception of low frequency sounds.

In several species of mammals, and interestingly in those which have the best sense of hearing (bats), a lot of radial myofibrocytes are included within the annulus tympanicus (which encircles the eardrum). These myofibrocytes are closely and individually connected to the radial fibres of collagen^S^^[15]^ attached to the bony circumference of the eardrum. So their activity may help to regulate the tension of the eardrum by means of that ring comprising radially oriented smooth muscles^S^^[16]^. This arrangement suggests a role in creating and maintaining tension on the tympanic membrane much like a drum tuning key which is used to adjust the “*tension rods*” and pitch of a drum.

Cf. below : Si 08

**Si 03B Variation of contact pressure of the electrodes on the skin**

A stronger pressure of an electrode on the skin might increase admittance of the pathway between the electrode and the collagen under the skin (owing to better contact and/or capacitive modification). It is therefore likely that the strength of pressure from the electrodes on the skin of knees should have some slight consequences on the results. However, we did not accurately study it.

Regarding the measurement on the eardrum, we face two antagonist phenomena: more pressure, better contact and better conductivity; But also, more pressure, less freedom of movements, so damping of vibrations and decrease of the associated piezoelectricity.
That is the conundrum that we face without being able to solve it at the moment.

#### Si 04 Possible Artefacts ?

**Si 04A Artefact resulting from links to ground**

If two pieces of electronic equipment are plugged into different power outlets, there will often be a difference in their respective ground potentials. The two ground connections may cause ground loops when the components are interconnected by signal cables (Fig. S02 below). There will be a parasitic standing wave created at the AC mains base frequency (50 or 60Hz) and the [harmonics](https://en.wikipedia.org/wiki/Harmonic) thereof (120 Hz, 240 Hz, and so on). So it is preferable to have "single-point grounding", with the system connected to the building ground wire at only one point. To avoid this phenomenon as much as possible, we have measured the validity of ground link.

Furthermore we have connected the different outlets of the system (lock-in, speaker) on the same power strip and the same ground socket.

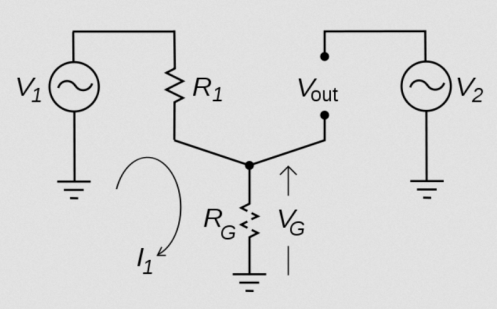

Figure S07
"ground artefact" excerpted from en-WP***^S^***^[17]^ (R resistance, G ground, V Voltage, I intensity; in this schema there are three links to the ground).

**Si 04B** **Artefact resulting from a “hand effect”**

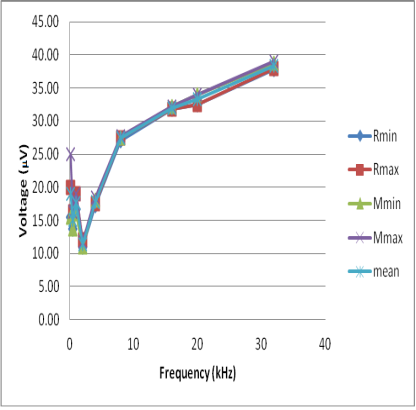

Figure S08
Mean Voltages (μV) measured for several frequencies
(R means Results without Hand Effect; M means results with Hand effect). In all configurations, when there is a change, the "hand effect" causes an increase of the voltage <1.5 μV for voltages around 25 μV , ie 6% (Fig. S08 above).

Additional connection of a conducive bracelet around the wrist of the experimenter engenders a higher "hand effect" unless the metallic case is also connected to the ground. The most practical solution, in order to avoid any hand effect, is to remove the hand of more than 20 cm from the casing.

**Si 04C Artefact depending on the contact between electrodes and the skin-target**

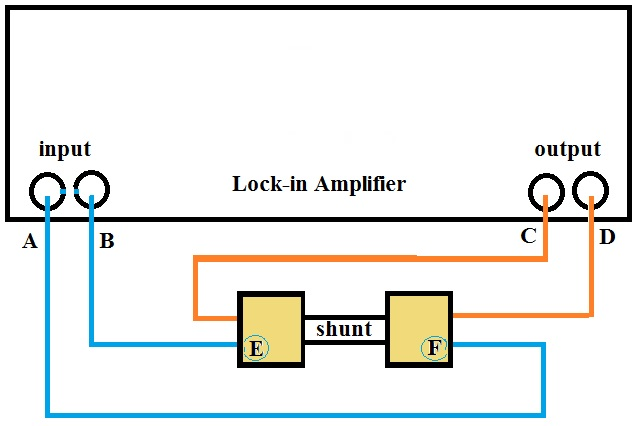

Figure S09

In fig. S04 (above) we replaced the skin of the subject by a simple copper shunt, with a resistance Rs = 0.2 Ω (AOIP 02.2351); This shunt is purely resistive up to a few tens of kHz. And the current from the output of the Lock-in (C and D) is introduced (by red circuit: coaxial cable and alligator clips) into the shunt.

The electrodes (E and F), normally applied to the skin, here are connected by the blue wires to the Lock-in Amplifier; they have different forms and surfaces (round or square COMEPA, or Alligator Clips).

Since our measures on tendons present values in the range from 10 to 200 µV, we adjust the settings so that the potential difference between E and F is close to 200 µV. As the resistance of the shunt is 0.2 Ω, we must send a current of 200 µV / 0.2 Ω, ie 1mA flowing through this shunt. To arrive at this value of 1 mA, if we take the device as a whole, including the two resistors in series, the output of the Lock-in (Ro = 50 Ω), and the resistance of the shunt (Rs = 0.2 Ω), we have to adjust the output voltage of the Lock-in to about 50 mV since {1 mA * (Ro + Rs)} = 1 * 50.2 = 50.2 mV. The resulting voltage between E and F (in µV) to the terminals of the shunt is measured at the input of the Lock-in (be A and B; fig. S04 above) with the blue cables. By varying the frequency (kHz), we obtain the results presented in Table S01 and Fig. S05.

|  | **two square electrodes** | **one round and  one square electrode** | **two round electrodes** | **two alligator clips** |
| --- | --- | --- | --- | --- |
| **f (kHz)** | (µV) | (µV) | (µV) | (µV) |
| **0.2** | 0.135 | 0.142 | 0.164 | 0.196 |
| **0.8** | 0.136 | 0.138 | 0.166 | 0.198 |
| **2** | 0.138 | 0.145 | 0.165 | 0.196 |
| **4** | 0.142 | 0.146 | 0.160 | 0.196 |
| **8** | 0.134 | 0.135 | 0.143 | 0.202 |
| **20** | 0.185 | 0.188 | 0.198 | 0.219 |
| Ro=50 Ω et Rs=0.2 Ω  **Table S08** | | | | |
| 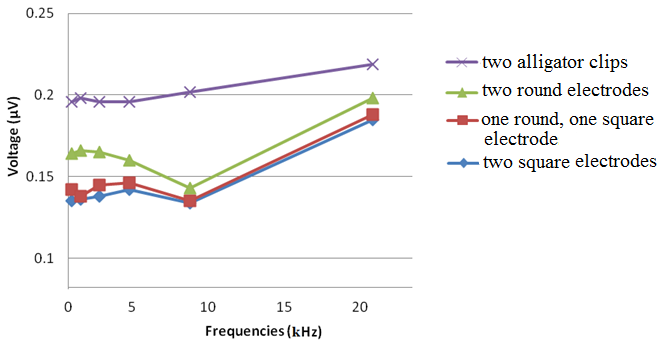  Figure S10 | | | | |

**Si 04D Is there artefact depending on microphonic of measurement cables ?**

Mechanical noise can be translated into electrical noise by microphonic effects. Physical changes in the cables (due to vibrations for example) can result in electrical noise over the entire frequency range of the lock-in. For example, consider a coaxial cable connecting a detector to a lock-in. The capacitance of a coaxial cable is a function of its geometry. Mechanical vibrations in the cable translate into a capacitance that varies in time typically at the vibration frequency. Since the cable is governed by CV= Q*^[18]^* taking the derivative yields: C(dV/dt) + V(dC/dt) = dQ/dt = i. Mechanical vibrations in the cable which cause a dC/dt will give rise to a current in the cable. This current may affect the measured signal ^S^^[19]^. We conducted an evaluation of this effect in the context of our measurements: We set the power of the loudspeaker by the lock-in with an increased voltage (5V against 2.5V for previous experiments) and with the speaker very close to the coaxial cable that we use. Variations in capacitance of the coaxial cables that we used, are very low given the broadcast sound power. We observe no voltage cable, short circuit, or open on the range 0.1 - 40 kHz.

In these measurements on metal, no sound is emitted; it is a classical voltage measurement. The increase here is due to the capacitive effect of the measurement contacts. For 20kHz, the increase is less than 30% against a factor 3 to 10 in the case of the tendons or the eardrum; it cannot, therefore, be an artefact in the case of the tendon or the tympanum. This increase is observable also in the measurement using alligator clips. This parasitic capacitance comes not only from the bad contact between the electrodes and the skin, but also from the contact between clips and electrodes.

In the case of electrophysiological recordings, it is usual to optimize skin impedance by cleaning the skin (Poch-Broto et al., 2009) and drying it, when necessary. The impedance of the path of contact also depends on the thickness of the subcutaneous fat layer, which might be roughly assessed by the Body Mass Index (BMI). There were not a significant correlation between that measure and the voltages results; So we think influence of this parameter is weak (see §statistics).

#### Si 05 About a supposedly cochlear origin of the isochronous response which we measured between the tympanic and mastoidian fibers ends

It is possible to detect an electric potential synchronous to the acoustic vibration between an indeterminate point of the tympanum and the mastoid bone***^S^***^[20]^. It does not follow necessarily, however, that the potential measured in this type of experiment is produced by the Outer Hair Cells (OHCs).

**Si 05A Value of the microphonic potentials**

Microphonic potentials (119), measured near the cochlea, are on the order of μV at the threshold. They increase from 100 to 400 μV for average sounds and up to a maximum of 800 μV for the most intense sounds***^S^***^[21]^.

Proto-microphonic potentials (about 10% of the microphonic that persists despite the functional elimination of the OHCs), would be around 1 to 80 μV. These values correspond to 1/1000 of the experimental values measured in vitro on collagen I (149). That residual potential persists in cases where the OHCs or IHCs are destroyed or no longer function for whatever reason; This persistence obviously cannot be attributed either to the OHCs or to the IHCs. In addition, the destruction of the IHCs (chinchilla) does not alter the cochlear microphonic and does not alleviate the Electrically Evoked Oto Acoustic Emissions (EEOAE); moreover, these responses tend to increase at high frequencies ^S[22]^. That being the case, we suggest that the residual potential is due to the piezoelectricity of the tympanum (the piezo-tympanic signal or "pT").

We have been able to record, from the region of the mastoid bone, synchronous potentials which are not weakened when the auditory canal is occluded (Cf. § hereafter). Similarly, if a sound is sent to the eardrum and not to the superficial mastoid, recorded synchronous potential is lower than if the sound is sent to the superficial mastoid and not to the eardrum. Therefore, we demonstrate that if a sound is sent in the direction of the mastoid area, the synchronous evoked potential is not from the cochlea, but from local generators. This means that neither the ossicular chain nor the Bekesy Traveling Wave (TW) is involved. Rather, the synchronous potential is attributable to the collagen present in the mastoid region.

**Si 05B Could tympanic or bone fibers isochronous potentials be from cochlear origin ?**

It has been sometimes accepted in the literature that synchronous potentials recorded at the level of the mastoid are purely of cochlear origin. If this were the case, obstructing the external ear canal should indeed lead to a reduction of these synchronous potentials.

The cochlear microphonic (CM) is an alternating current (AC) isochronous to the acoustical stimulating waves^S^^[23]^. Its threshold is very low. Above 60 dB SPL its amplitude decreases almost at once. On the contrary, for moderate incident sounds (< 40 dB SPL), an amplification, proportional to the sound level occurs after a certain delay. This amplification is, for the most part (80-90%), attributable to OHCs, but it resists the destruction of these OHCs. The origin of this poorly understood residual 10-20 %, has been mistakenly attributed to the IHCs alone^S[24]^ or to the piezoelectricity of the membrana tectoria (Offut, 1984). Our hypothesis is that this residual microphonic is generated, at least in very large part, by the piezoelectricity of the eardrum (and/or petrosal collagen). The so-called mastoid microphonic is not reducible to the diffusion of the cochlear microphonic. In Fig.S06 we describe this commonly accepted theory. The cochlear microphonic would be detectible near the eardrum***^S^***^[25]^ and it would also be detectible near the mastoid^S^^[26]^. This mastoid potential would be the emergence of the cochlear microphonic alone^S^^[27]^ and would allow for audiometric testing^S^^[28]^. Mammano^S^^[29]^ thinks that the synchronous potentials that we highlight could be interpreted as the result of the activity of the OHCs reaching as far as the eardrum (or mastoid) ....

Furthermore, in order to build a tympanum restricted to its mechanical effects, the evolutionary process should have led to the generation of collagen fibres in a convenient geometrical arrangement, but with random polarity.

We will show below that :

- The cochlear microphonic is not dependent either on the IHCs, or on cochlea

- The so-called mastoid microphonic is not reducible to the diffusion of the cochlear microphonic.

- Attempts to define a mastoid audiogram has appeared problematic.

- Bone conduction allows to hear ultrasounds.

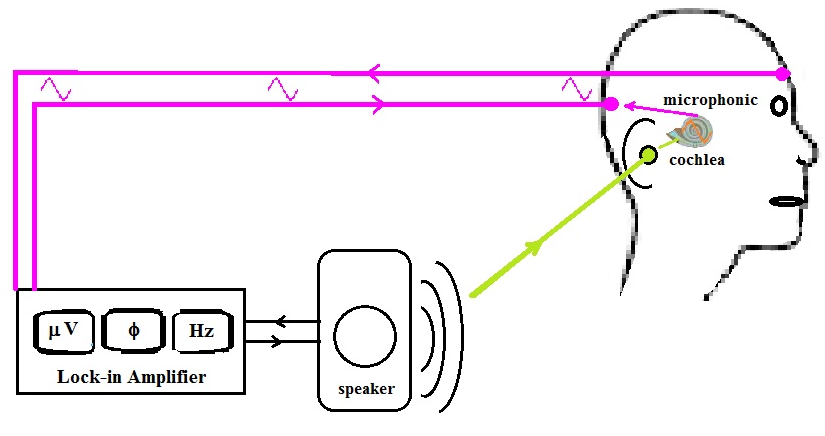

Figure S11

Sometimes accepted theory of the generation of the microphonic near the mastoid

The acoustic vibrations of the eardrum are transduced, by the cochlear OHCs, into a synchronous electric potential, known under the name of microphonics^S^^[30]^. Furthermore, such a synchronous electric potential can be recorded at the level of the skin behind the ear (with the other electrode being placed, for example, on the forehead). It is postulated that the cochlear generator alone would be responsible for that recorded potential. In other words, the recorded potential would result only from the scattering of the "cochlear microphonic" and for no other reason.

**Si 05C The synchronous potentials, tympanic or mastoidian,
 are not uniquely dependent on the cochlea**

In order to test whether pure-frequency sound stimulation produced a piezoelectric response, we measured the tympanum of a pig a few hours after its death, and after removal of the cochlea performed without damaging the anatomy of the eardrum.

**Piezoelectricity measurements on the pig tympanum at the Puylaurens abattoirs (1 07 2018).**

We carried out measurements on pig eardrums shortly after their slaughter, in the Puylaurens Archeology Club premises (room temperature 24 ° C).

**Acoustic measurements**
Sound source: Yamaha DBR12 loudspeaker, powered by the LockIn Stanford SR 830 (frequency and amplitude adjustment).
The sound intensity measurement was carried out using a Bruel & Kjaer sound level meter, Type 2231 (measurements were made by Bernard Bibé, INRA). For the tests, we opted for a relatively moderate acoustic amplitude of about 75 dB SPL
archeology of Puylaurens (room temperature 24 ° C).

**Measurements of electrical responses:**
Lock in Standford SR 830 (measuring frequencies 150 Hz-30kHz, 300ms time constant).
The probes consist of a coaxial cable that is connected to the input of the LockIn by a BNC plug.
This cable has two components: a fiber core consisting of a single fiber and a peripheral shield made of very fine braided metal fibers. The end of these two components is completed by a small metal ball.

**Measured samples:**

1. **Tympan 1** taken from a head of the day, not scalded.

2. **Tympan 2** taken from a scalded head

3. **Tympan 3** taken from a head of the day, not scalded but damaged during preparation.

4. **Bone conduction**: Bone taken from the external auditory canal, this bone is hemicylindrical (semicircular section). It consists of a wall of X mm thick, and a "channel" of Y mm width; this wall and this channel are oriented like generatrices of the bone cylinder.

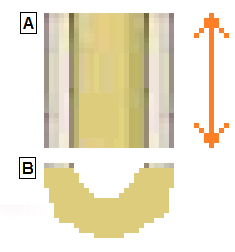

A ) Superior view of the osseous canal.
On the right: double arrow symbolizing the potential difference between the ends of the bone channel, when it is stimulated by a stable frequency acoustic wave.
B ) Sectional view of the 1/2 bony channel

6 measurements were made:

1) Tympanum 1: loudness 70dB
2) Tympanum 2: loudness 74dB
3) Tympanum 2 after removal of the cochlea: loudness 74dB
4) Tympanum 3: loudness 74dB

Hemisection of the bone canal:
5) loudness 70dB
6) loudness 90dB

We have calculated (Excel) the polynomial-type trend curve for each of the measurements made

**Measurement 1**: Tympanum 1 taken from a head of the day, not scalded: stimulus 70dB SPL

| Frequency (Hz) | Stimulus  (dB SPL) | Electrical response (μV) |
| --- | --- | --- |
| 200 | 70 | 34 |
| 500 | 70 | 9 |
| 1000 | 70 | 10 |
| 2000 | 70 | 4 |
| 4000 | 70 | 0 |
| 8000 | 70 | 3 |
| 16000 | 70 | 1,5 |

Tympanum 1; y(μV)= 2E-07 x^2^ - 0,005 x + 18,59; R² = 0,47

ordinates: voltages (μV) ; abscissa: frequencies (Hz)

**Measurements 2**: Tympan 1 (Stimulus amplitude 70dB SPL)

| Frequency (Hz) | Piezoelectric signal (μV) |
| --- | --- |
| 500 | 0,2 |
| 1000 | 0,5 |
| 2000 | 0,2 |
| 4000 | 0,06 |
| 8000 | 0,08 |
| 16000 | 0,3 |
| 20000 | 0,27 |
| Although the signal was very weak, the measurements were stable during the measurement and the repetition of the measurement gave an identical result. | |

y = 2E-09 x^2^ - 5E-05 x + 0,3214; R² = 0,30
ordinates: voltages (μV) ; abscissa: frequencies (Hz)

**Measurements 3**: Tympan 2 taken from a head of the day, not scalded, after ablation of the cochlea, sound intensity 74dB SPL.

| Frequency (Hz) | Piezoelectric signal (V) |
| --- | --- |
| 250 | 4 |
| 500 | 0,25 |
| 1000 | 0,15 |
| 2000 | 0,18 |
| 4000 | 0,07 |
| 8000 | 0,11 |
| 16000 | 0,2 |

y = 2E-08 x^2^ + 1,62 ; R² = 0,26
ordinates: voltages (μV) ; abscissa: frequencies (Hz)

**Measurements 4**: Tympanum 3 (stimulus amplitude 74dB SPL)

| Frequency (Hz) | Signal piezo (V) |
| --- | --- |
| 250 | 20 |
| 500 | 6 |
| 1000 | 3 |
| 2000 | 2 |
| 4000 | 0,4 |
| 8000 | 0,8 |
| 16000 | 1,4 |

y = 1E-07x^2^ - 0,003 x + 10,51; R² = 0,44
ordinates: voltages (μV) ; abscissa: frequencies (Hz)

**Measurements 5**: Piece of External Bone Canal (70dB)

| Frequency (Hz) | piezoelectric signal (V) |
| --- | --- |
| 150 | 0,87 |
| 500 | 0,5 |
| 1000 | 0,75 |
| 2000 | 1 |
| 4000 | 0,3 |
| 8000 | 1,1 |
| 16000 | 2,7 |

y = 1E-08x^2^ - 8E-05x + 0,7737; R² = 0,91
ordinates: voltages (μV) ; abscissa: frequencies (Hz)

**Measurements 6**: Piece of External Bone Canal (90dB)

| **Frequency (Hz)** | **piezoelectric signal (μV)** |
| --- | --- |
| 150 | 25 |
| 500 | 25.74 |
| 1000 | 4 |
| 2000 | 6,5 |
| 4000 | 1,7 |
| 8000 | 6 |
| 16000 | 15,6 |

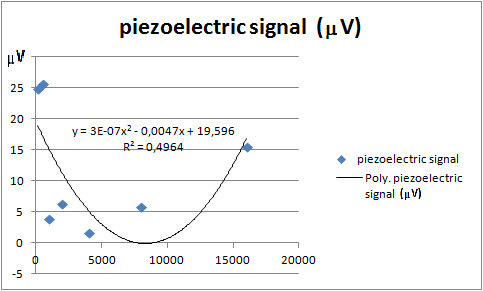

y = 1E-06x^2^ - 0,025x + 96,433; R² = 0,20
ordinates: voltages (μV) ; abscissa: frequencies (Hz)

**Si 05D Bone Conduction**

**Si 05Da Attempts to define a mastoid audiogram have been problematic**

These findings and their restrictive interpretation led Poch-Broto and al. (2009) to try to create an audiometric test based on the recording of the electric potential of the mastoid in response to auditory stimulations. In this theory, the schema of transmission without earplug complies with fig. S07.

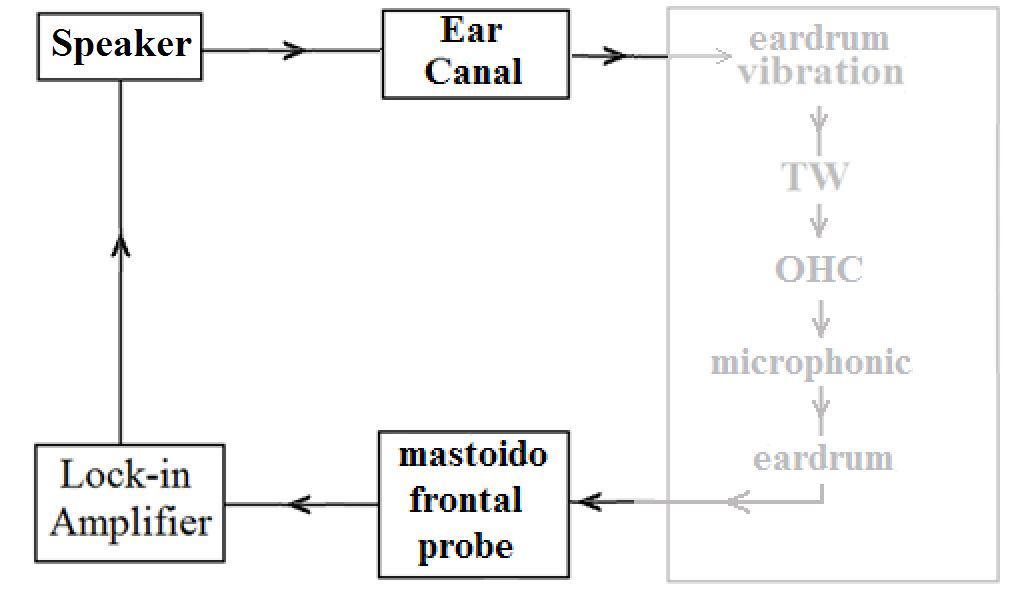

Figure S12

Capture of the mastoid “microphonic”. Dimmed: Interpretation of Poch-Broto, attributing all the microphonic amplification to the TWs alone, via the OHC***^S53^***.

But, if we compare the classic audiogram and this " mastoid " audiogram, we can see that very often (>10%) they differ by more than 10 dB HL mainly in the area of high frequencies, which is significant (fig. S08).

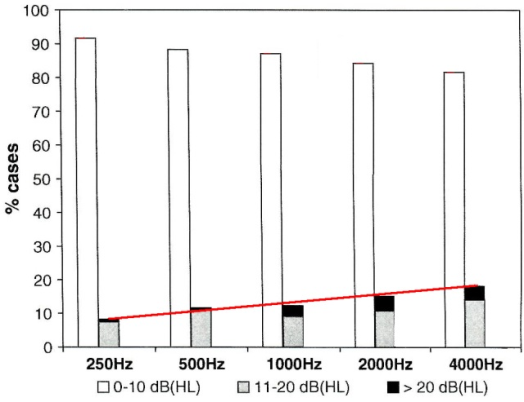

Figure S13

Modified chart based upon the data of Poch-Broto et al., 2009***^S^***^[31]^:

We grouped the black rectangles and the gray rectangles to represent all errors (> 10dB) combined. This clearly shows (red lines) that the error rate increases with frequency. The X axis displays the tested frequencies; the Y axis displays the % of cases according to whether the divergence between subjective audiometry and "microphonic" audiometry is less than (clear bars), or superior to (gray and black bars) 10 dB HL.

Since for all frequencies, in more than 10% of cases, the difference between subjective audiometry and measurement of the microphonic is greater than 10 dB, it can be concluded that the cochlear generator is not the sole origin of collected potentials (cf the line in red); this is particularly true for the high frequencies. We contacted Pablo Gil Loyzaga (corresponding author of the Group of Poch Broto and al.) and asked for their raw data. Yet, he answered that the raw data was no more available (Pablo Gil Loyzaga, com. pers., 07/18/2012).

**Si 05Db Bone conduction permits the hearing of ultrasounds**

Bone collagen is particularly adapted for the transduction of ultra-sounds***^S^***^[32]^. Ultrasounds between 20 kHz and 33 kHz are heard when they are being administered by bone conduction***^S^***^[33] -^ ***^S^***^[34]^. They act directly on the hair cells (ie their hearing is induced by the ultrasounds themselves). And furthermore, in the case of extremely high pitched sounds, toxic damage of the auditory system damages the air-conduction hearing and improves bone-conduction hearing.

It is well demonstrated that the deterioration of collagen II is causative of auto-immune deafness***^S^***^[35]^.

**Si 05E Experiments involving closing off of the external auditory canal (EAC)**

The schemas, tables and curves below show the synchronous potentials, depending on whether a plug is not or is inserted into the auditory external conduit (earplug interposed between the sound source and the eardrum, but not between the sound source and the mastoid) : fig. S09, S10, S11, S12.

**Si 05Ea** **Mastoid 'microphonic' with EAC closed off :**

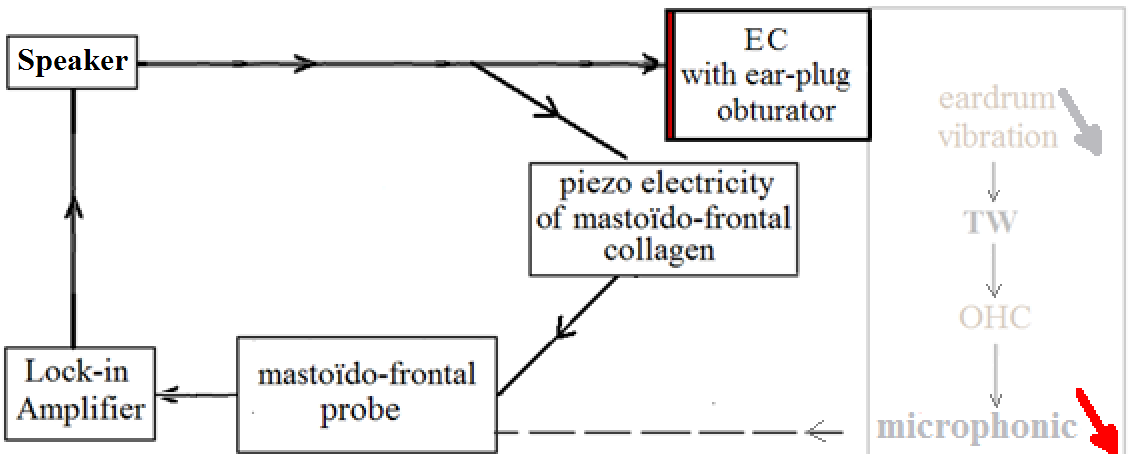

Figure S14

Capture of the mastoid 'microphonic' with the external auditory conduit closed off. (Dimmed and red arrows on the right side of the figure indicate the decrease of the mechanical vibration and of the electric potential as well).

##### Si 05Eb Mastoid potential with EAC closed off by a standard shutter of trade:

We can insert a plug to seal off the external auditory canal (EAC). The theory of Poch-Broto et al. (2009) implies that the earplug decreases the amplitude of the vibrations of the eardrum. This necessarily results in a significant decrease of the cochlear "microphonic"; And of the mastoid synchronous potential as well if the latter potential results solely from the cochlear "microphonic". If the mastoid synchronous potential is composite, and if it is generated, at least in part, by the piezoelectric activity of the mastoid collagen, the mastoid synchronous potential should be only slightly diminished by the closing off of the external auditory conduit. But, if for some subjects, the synchronous potential remains the same, whether the ears are plugged or not, we have to consider another phenomenon.

We, therefore, conducted the following experiments. Three subjects were tested (D37, D39, D44). We display the electrical response depending on whether the EAC is open or obstructed (F is the frequency of the auditory stimulus, in kHz; Vs is the electrical potential of the response, in μV).

The schemas, tables and curves below show the synchronous potentials, depending on whether an earplug is (fig. S09 above) or is not (fig. S07 above) inserted into the external auditory conduit (earplug interposed between the sound source and the eardrum, but not between the sound source and the mastoid). The earplugs used were “*quies*”, made from natural wax, with an assumed protection value reported by the manufacturer of 27 dBs^[36]^ (Noise Reduction Rate) for the subject D37 (Table S02**a**: Fig.S10), subject D39 (Table S02**b**): Fig.S11), subject D44 (Table S02**c**; Fig.S12).

| Mastoïdo-frontal potential [Vs (µV)] without earplug (“open”) or with earplug (obstructed) | | | | | | | | | |
| --- | --- | --- | --- | --- | --- | --- | --- | --- | --- |
| Table N° | **a** (D37, 51 years old) | | | **b** (D39, 17 years old) | | | **c** (D44, 25 years old) | | |
| Figure N° | S10 | | | S11 | | | S12 | | |
| **f (kHz)** | **Open** | **closed** | **Δ** | **Open** | **closed** | **Δ** | **Open** | **closed** | **Δ** |
| 0,2 | 0,2 | 1,7 | -1,5 | 1,7 | 2,2 | -0,5 | 12,5 | 6,2 | -6,3 |
| 0,5 | 0,5 | 1,7 | -1,2 | 1,7 | 2,2 | -0,5 | 10,3 | 6,6 | -3,7 |
| 1 | 2,2 | 2,8 | -0,6 | 10 | 12 | -2 | 12 | 9,5 | -2,5 |
| 2 | 2,5 | 3,7 | -1,2 | 11 | 19 | -8 | 22 | 14 | -8 |
| 4 | 3,3 | 5,2 | -1,9 | 17 | 29 | -12 | 37 | 24 | -13 |
| 8 | 5,6 | 7,2 | -1,6 | 50 | 50 | 0 | 55 | 46 | -9 |
| 10 | 6,5 | 8,5 | -2 | 55,2 | 64 | -8,8 | 73 | 56 | -17 |
| 15 | 8,3 | 10,4 | -2,1 | 95,2 | 90 | 5,2 | 67 | 70 | 3 |
| 20 | 10,4 | 11,3 | -0,9 | 139 | 120 | 19 | 80 | / | na |
| **Mean (**µV **)** | 4,7 | 5,9 | **-1,4** | 53,9 | 54,9 | **-1** | 29 | 36,1 | **-7,06** |
| Table S09 (**a**, **b**, **c**)  Synchronous mastoid voltage (µV), broadcasted by a loudspeaker Harman/Kardon, stimulated by a voltage of 2 Volts issuing from the lock-in amplifier; Δ = Vs *with (closed)* minus Vs *without (open)* | | | | | | | | | |

| Figure S15 **a** (D37, 51 years old).. | Figure S16 (Ord. Volts)  **b** (D39, 17 years old) |
| --- | --- |

|  | Open | closed |
| --- | --- | --- |
| 0.2 | 0.2 | 1.7 |
| 0.5 | 0.5 | 1.7 |
| 1 | 2.2 | 2.8 |
| 2 | 2.5 | 3.7 |
| 4 | 3.3 | 5.2 |
| 8 | 5.6 | 7.2 |
| 10 | 6.5 | 8.5 |
| 15 | 8.3 | 10.4 |
| 20 | 10.4 | 11.3 |

|  | Open | closed |
| --- | --- | --- |
| 0.2 | 1.7 | 2.2 |
| 0.5 | 1.7 | 2.2 |
| 1 | 10 | 12 |
| 2 | 11 | 19 |
| 4 | 17 | 29 |
| 8 | 50 | 50 |
| 10 | 55.2 | 64 |
| 15 | 95.2 | 90 |
| 20 | 139 | 120 |

|  | Open | closed |
| --- | --- | --- |
| 0.2 | 12.5 | 6.2 |
| 0.5 | 10.3 | 6.6 |
| 1 | 12 | 9.5 |
| 2 | 22 | 14 |
| 4 | 37 | 24 |
| 8 | 55 | 46 |
| 10 | 73 | 56 |
| 15 | 67 | 70 |

Figure S17
**c** (D44, 25 years old)

The three figures (S10, S11 and S12) show that the mastoid potential with the ear plugged

was very close to the mastoid potentials of the same ear when not plugged. Furthermore, regarding the relation between the with earplug and without earplug representative lines, there are three configurations.

In the case of D37 (fig. S10), the magnitude of the "mastoid potential" with airplug (earplug “quies”, pure wax 27 dB) is above that of the without airplug situation. In the case of D39 (fig. S11), the magnitude of the "mastoid potential" "with" airplug is intertwined with that of the "without" situation. In the case of D44 (fig. S12), the magnitude of the mastoid potential "with airplug" is below that of the "without" situation.

However, when the ear is clogged, the signal that reaches the cochlea is necessarily reduced (standard theory) and therefore the cochlear potential should be weakened as well, in all cases! Thus, as logical consequence, the mastoid potential we recorded cannot depend solely upon the cochlear microphonic. Moreover, our measures are consistent with the hypothesis that the mastoid potential depends not only on the impact of sound on the eardrum, but also on the mastoid reaction. So we think that the 'mastoid potential' results, at least in part, from the electrical signal generated by the piezoelectricity of collagenous fibres, located between the points of application of the electrodes (mastoid and forehead). According to our theory linked to the standard interpretation, this piezoelectric response is added to the electric signal from the cochlea.

##### Si 05Ec Experiment using 'active noise cancelling headphones' plus ‘custom moulded earplugs' (formed in place)’:

However, on the three cases above, the occlusion of the external auditory canal, by means of a standard over-the-counter earplug (“quies”, pure wax 27 dB) was, as such, limited to a maximum of 27 dB HL. So, the effects obtained with "earplugs" should be even more marked with “custom-made earplugs (fig. S13) + active noise cancelling headphones (fig. S14 & fig. S15)”. We, therefore, repeated the previous experiments, using active noise cancelling headphones in the context of the situation 'occluded ear'. We filled the external auditory canals of one subject with custom-made earplugs (Audial, Toulouse, Fig. S13), completed by active noise cancelling headphones, (Panasonic RP - HC 500; Fig. S14); Together : fig. S15.

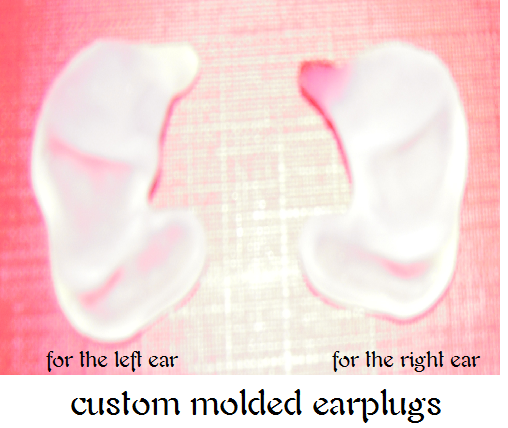

Figure S18

custom-made earplugs

We measured the audiometric thresholds using only custom-made earplugs, then using the active noise cancelling headphones alone (fig. S14), then using both (fig. S15). We used the same placement of the electrodes as in the previous experiments. The previously positioned electrodes, held in place by surgical tape, are pressed by the subject himself, the potential measured depending on the pressure on the skin. The Yamaha HS5 speaker is installed at 1 m from the ear of the subject, its centre facing the subject at the height of the ear to be tested. We performed a calibration of acoustic amplitude with a *Bruel and Kjaer 2231* sound level meter, the rod bearing the microphone being perpendicular to the face of the speaker. The voltage of the output of the Lock-in, plugged into the speaker is set so that the sound level meter displays about 80 dB.

| 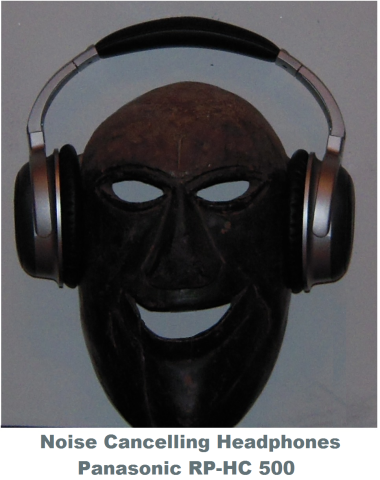 | 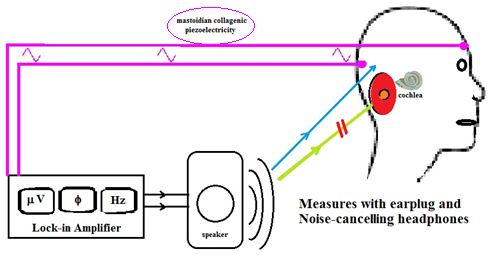 |
| --- | --- |
| Figure S19 | Figure S20 |

We present below (subject D38, Table S03, Fig. S16), the voltage output of the Lock-in amplifier and measurements of the synchronous electrical potentials, with (Plugged: P) and without (Unplugged: U) occlusive equipment. The data are displayed according to the sound frequency (f kHz) emitted by the speaker, either toward the right auditory canal (REAC) or toward the left one (LEAC).

| f (kHz) | Output voltage (to the speaker) | Vs (μV) U-REAC | Vs (μV) P-REAC | Vs (μV) U-LEAC | Vs (μV) P-LEAC |
| --- | --- | --- | --- | --- | --- |
| 0.5 | 0.4 V | 5 | 5 | 6 | 6 |
| 1 | 0.3 V | 6 | 7 | 5 | 7 |
| 2 | 0.5 V | 6 | 5 | 5 | 6 |
| 4 | 0.2 V | 9 | 7 | 8 | 7 |
| 8 | 0.2 V | 18 | 15 | 13 | 14 |
| 16 | 0.4 V | 44 | 42 | 45 | 43 |
| 20 | 1 V | 123 | 96 | 110 | 105 |
| 32 | 1 V | 144 | 143 | 170 | 145 |
| Table S10  External Auditory Canal (EAC); subject D38,  Right (R), Left (L), Unplugged (U), Plugged (P) | | | | | |

7
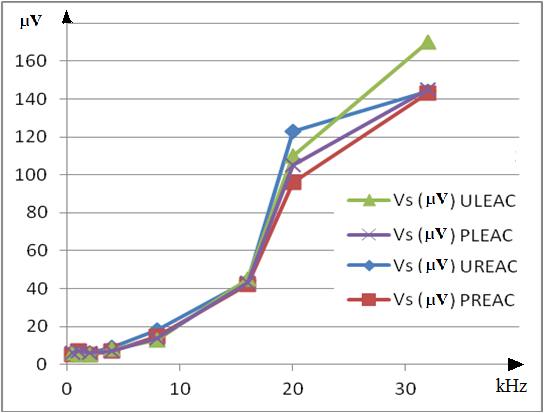

Figure S21

We found that the mastoid potentials with the ear plugged was very close to the mastoid potentials of the same ears when not plugged. As the mastoid potentials are not greatly weakened by plugging the EAC, the classical theory seems to be insufficient and we think that another phenomenon coexists. We hypothesize that this phenomenon is the piezoelectricity of the mastoid collagen. We, therefore, propose the two diagrams below (Fig. S17) in which we have added the effect of the piezoelectricity of the mastoid collagen (blue line) to the classical schema (green line) for purposes of comparison.

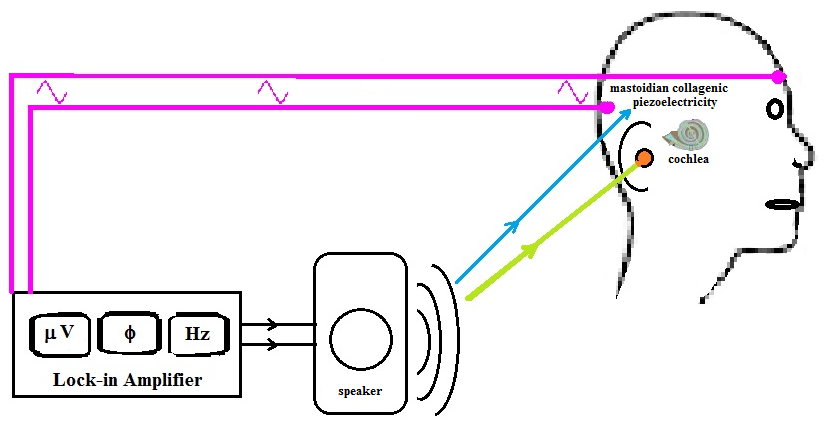

Figure S22
Stimulation restricted to the external auditory conduit (stimulation by headphone only) compared with stimulation targeting the mastoid (stimulation by loudspeaker only)

##### Si 05F Conclusion on Si 05 :

If one puts an *earplug* (in red) or/and *standard headphones* and/or "*active noise cancelling headphones"* on the ears of the subject, the acoustic vibrations (in green) encounter this obstacle which necessarily weakens the vibrations of the eardrum; Thus, in the accepted theory, the cochlear generator will produce a synchronous potential of a lower amplitude. The result would be that the synchronous electric potential measured at the level of the skin behind the ear (μV) should logically be reduced. Since it does not, we conclude that the cochlea is not the only generator of this potential.

Collagenous fibers of the mastoid region have roughly the same structure as the fibers belonging to the patellar ligament. It follows that the collagenous fibers of the mastoid are able to generate a piezoelectric potential (synchronous to the acoustical vibrations) just like the potential we measured at the level of the patellar ligament. Acoustic vibrations produced by the speaker (blue arrow) stimulate an electric non-cochlear generator, i.e. the piezoelectric collagen fibers of the mastoid.

Plugging the ear canal should not only decrease very sharply the cochlear microphonic, but it should also decrease the petrous resonances caused by the vibrations of the eardrum ! It is possible that the slight difference in favor of the non-blocked ear comes, in part or entirely, from the decrease in the phenomenon of resonance when the ear is plugged.

We may conclude that our measures are consistent with the hypothesis that the mastoid potential depends not only on the impact of sound on the eardrum, but also on the mastoid structures. We think that the 'mastoid potential' results mainly from the electrical signal generated by the piezoelectricity of collagenous fibres located between the points of application of the electrodes (mastoid and forehead).

**Si 06 Source of the mastoid synchronous potential**
A diagram of the reception of the external synchronous potential, with the mastoid piezoelectricity eliminated, is given on the Figure S18: the sound is not emitted by a loudspeaker but only through a headphone.

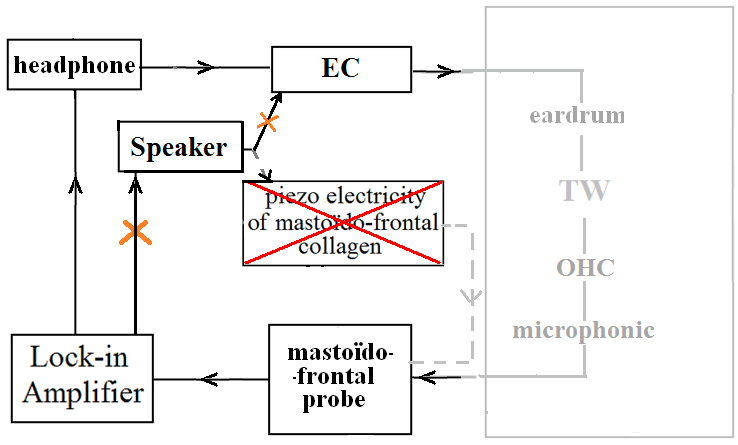

##### Figure S23 Stimulation restricted to the external auditory conduit (stimulation by headphone only) compared with stimulation targeting the mastoid (stimulation by loudspeaker only)

If the mastoid synchronous potential originated from the cochlea alone, then it would be about the same no matter whether the sound was broadcast through a loudspeaker or through the headphone.

If, on the other hand, the mastoid synchronous potential originates locally (at the mastoid), then the sound emitted by the headphone would produce a very low synchronous potential compared to the same sound broadcast through a loudspeaker.

**Si 06A Testing hypotheses: purely tympanic way vs. mixed path**

The standard theory states that, if the sound reaches the eardrum without reaching the surface of the mastoid, the synchronous potentials will not weaken (Hypothesis H0). The alternative hypothesis (H1) states that the synchronous potentials will weaken. We checked H0 versus H1.

To allow comparison of stimulation through the auditory canal alone and stimulation by the loudspeaker (free field), we replaced the loudspeaker by a headphone and we fed it in such a way that the subjective magnitude would be roughly the same as with the loudspeaker (prior calibration).

For predetermined frequencies (200 Hz, 1 kHz, 8 kHz, 15 kHz), we provided an operatic singer  (D40) with a reference acoustical amplitude by means of a loudspeaker placed at approximately 1 m from his ear.

Then, we sent the same frequency by headphone (audio InterSound HD75, without volume attenuation) fed from the output of the lock-in amplifier. We asked the singer to modify the electrical voltage generated at the output of the lock-in amplifier, connected to the headphone, until he got the same subjective impression as with the loudspeaker (see results Table S04).

| Frequency (kHz) sent by the lock-in amplifier | Voltage (mV) sent by the lock-in amplifier |
| --- | --- |
|  | to the loudspeaker |
| 0.2 | 5.10^3^ |
| 1 | 3.10^3^ |
| 8 | 104 |
| 15 | 200 |
| Table S11 [Calibration](http://en.wikipedia.org/wiki/Calibration): D40, a professional operatic singer, is stimulated by a pure frequency tone, in turn via the headphone and via the loudspeaker; owing to the voltage from the lock-in amplifier, successive settings amplify these two transducers, in such a way that a subjectively identical amplitude of sound is fixed for different frequencies | |

But for his subjective convenience, we did not impose to D40, an identical standard amplitude for every frequency. For each of the tested frequencies, we allowed him to fit the respective amplitudes of the headphone and the loudspeaker, each in turn, by trial and error, so that the heard amplitude would be as much as possible identical for both, without aiming exactly the level proposed at the beginning.

Using the results of this calibration, the subject received an equivalent stimulation for each preselected frequency in turn, via either the headphone alone (situation A) or the loudspeaker alone (situation B).

According to Hypothesis H_0_, the mastoid synchronous potential comes from the cochlear microphonic. Consequently, the measured synchronous potential should be lower in situation B (loudspeaker only) than in situation A (headphone only).

According to Hypothesis H_1_, the mastoid synchronous potential comes mainly from the mastoid collagen piezoelectricity. In this case, the measured synchronous potential would be expected to be lower in situation A (headphone only) than in situation B (loudspeaker only).

We present the results in Table S05 for D40 and for two other subjects (D41 and D42) in Table S06 for D41, and in Table S07 for D42. In the tables, a dash indicates that no synchronous potential could be measured: absence or extremely low amplitude.

| D40 | Situation A  Headphone (μV) | Situation B  Loudspeaker (μV) | H_1_ vs H_0_ |
| --- | --- | --- | --- |
| 0.02 kHz | - | 4.6 | H_1_ supported |
| 1 kHz | - | 3.6 | H_1_ supported |
| 8 kHz | - | 0.27 | H_1_ supported |
| 15 kHz | 3.7 | 1.2 | H_0_ supported |
| Table S12 (D40) | | | |

| D41 | Headphone (μV) | Speaker (μV) | H_1_ vs H_0_ |
| --- | --- | --- | --- |
| 0.02 kHz | - | 20 | H_1_ supported |
| 1 kHz | 3.7 | 16 | H_1_ supported |
| 8 kHz | 6.5 | 2.5 | H_0_ supported |
| 15 kHz | 3.3 | 6.1 | H_1_ supported |
| Table S13 (D41) | | | |

| D42 | Headphone (μV) | Loudspeaker (μV) | H_1_ vs H_0_ |
| --- | --- | --- | --- |
| 0.02 kHz | - | 180 | H_1_ supported |
| 1 kHz | 1.8 | 440 | H_1_ supported |
| 8 kHz | 1.8 | 36 | H_1_ supported |
| 15 kHz | 11.5 | 90 | H_1_ supported |
| Table S14 (D42) | | | |

###### Si 06B Conclusion about the source of the mastoid synchronous potential :

Hypothesis H1 is borne out ten times on twelve measures. In more detail:

- Hypothesis H1 is clearly supported with regard to frequencies 0.02 and 1 kHz. For these frequencies, the synchronous mastoid potential seems, for the most part, to be produced by the mastoid region collagen fibres and, therefore, does not come from the cochlear microphonic...

- Only D40 at 15 kHz, and D41 at 8 kHz, do not support our theory.

Furthermore, the specific results observed for frequencies 8 kHz and 15 kHz can be explained by the fact that vibrations imposed on the eardrum should be transmitted to the mastoid by resonance..

Finally, it should be noted that D40, involved in the calibration, had characterized his subjective assessment regarding 15 kHz as a very rough approximation.

- But note also that hypothesis H1 is fully validated for all measurements in the case of D42, whose potentials are all very high, and who has, in addition, high-performance hearing thresholds and a cut-off frequency in the high pitched sounds above 22 kHz (which is largely above the audiometric standards).

Even if error bars are higher than the current estimate of half a unit. it remains that the standard theory (H0) is largely invalidated by our experiments.

This experiment suggests that the mastoid synchronous potential disappears or weakens if vibrations mobilize the eardrum rather than the surface of the mastoid. This being the case, we must add to the effect of the piezoelectricity of the tympanic fibres, the effect, though somewhat lesser, of the piezoelectricity of the mastoid fibres.

Moreover, we have demonstrated that the synchronous potentials, measured at the level of the mastoid, are not reducible to the cochlear microphonic in its classical conception, according to which they are solely the result of the mechanical TW, amplified by the OHCs of the cochlea. An important part of the mastoid synchronous potential has a local origin: it derives from the piezoelectricity of the collagen (type I) of the mastoid.

Of course, our measurements are different from the audiometric classic ones, but an attentive examination of both for comparison would be interesting to do***^S^***^[37]^ ***^- S^***^[38]^.

#### Si 07 The pT voltages are they able to reach the DOHC complex ?

Our measures of pT may seem of a too low tension to act on the cochlea; but we must take into account the fact that the result of our measures is diminished by the interposition between the probe and the source of the electric potential of an more or less insulating layer, the epidermis of the eardrum. When we take into account the weakening of conduction by the epidermal tympanic layer, these values seem consistent with our measurements on the eardrum in vivo (hundreds of microvolts for an acoustical stimulation of approximately 70 or 80 dB SPL).

Furthermore, Harnagea showed in vitro that the response is low or non-existent if the electrodes are arranged in any two points of the biological culture, but this response reached tens of mVolts if the electrodes are placed at both ends of a single fiber... That was confirmed by Denning et al. 2017***^S^***^[39] -^ ***^S^***^[40]^. Thus we can assume that the pT voltages are in the mV range and may reach the DOHC complex.

## Si 08

#### Si 08A The mystery of the cetacean auditory mechanism

The minimum intensity detectable by mammals is a function of the frequency. It is noteworthy that they show a weaker sensitivity to very low and very high frequencies according to an U-shaped pattern. Cetaceans, most notably the small odontocetes, possess extraordinary auditory faculties, (...). They make extensive use of sound in echolocation as well as communication behaviours. They have a U-audiometric curve, typical of mammals***^S^***^[41]^***.*** In all species tested, the maximum sensitivity is at frequencies greater than 15 kHz.

The greatest sensitivity of cetaceans lies between 30 and 80 kHz. Their sensitivity decreases between 40 kHz and 1 kHz, and also between 80 kHz and 150 kHz (cf. Table S08).

| Species | Best sensitivity | Min frequency | Max frequency |
| --- | --- | --- | --- |
| Beluga whale | 30 kHz | 20 kHz | 120 kHz |
| Killer whale | 40 kHz | 0.5 kHz | 100 kHz |
| Tursiops spp | 50 kHz | 2 kHz | 135 kHz |
| Inia geoffrensis | 80 kHz | 1 kHz | 105 kHz |
| Table S15 | | | |

With respect to the frequency selectivity capabilities of cetaceans (for ex. T. truncatus) they are comparable to humans at lower frequencies, and the best reported for any mammal above 20 kHz. Moreover, the smaller toothed whales, in particular, appear to combine high temporal resolution with extremely sharp frequency tuning. (...)

"There are very short response latencies and an extremely rapid conduction of the information through the auditory pathway. The results describe an auditory system that is highly specialized for the extremely rapid conduction of auditory information from the periphery to more central structures"***^S^***^[42]^.

The external auditory canal of cetaceans is, , vestigial, although there may be an opening (very small)***^S^***^[43]^. Their eardrum is an extended, thickened membrane, the function of which is unclear***^S^***^[44]^. Ossicles are non-existent or hindered by additional or reinforced collagen ligaments, very stiff and heavier in structure (more massive, denser, ... etc.). The attachment of the stapes at the oval window of the cochlea is very tight and seems rigid in postmortem specimens***^S^***^[45]^. Although the remains of the outer ear and the middle ear are detectable in odontocetes, it is not possible to attribute to them the recognized role in terrestrial mammals.	The attachment of the stapes at the oval window is very tight and seems rigid^[46]^. Although remains of the outer ear and the middle ear are detectable in odontocetes, it is not possible to attribute to them the same role as which recognized in terrestrial mammals.

_____________________

The transmission of acoustic vibration from the surrounding sea water toward the cochlea is thought to result from a "fatty transmission" via perimandibular "acoustic fat bodies". These acoustic fat bodies are in direct contact with the malleus ossicle of the ears, especially in odontocetes but also in some mysticetes.

In addition, fibroblasts by inserting themselves into the network of collagen fibres, are also suitable for turning into adipocytes^[47]^**,** the main constituents of the lipid acoustical pathway.

The role of the eardrum and these fatty acoustic bodies are very similar ; They provide double transduction. Acoustic vibrations mechanically mobilize ossicles or their remains; This phenomenon is useful for the perception of low acoustic frequencies. Ear fat attaches to the tympanoperiotic complex, inserting itself into the "triangular opening", at the entry to the middle ear, where the ear fat contacts the hammer. Thus, the "acoustic signal" transmitted by the acoustic fat bodies, like the tympanic vibrations of terrestrial mammals reaches the hammer.

- Since the fibrous collagens are piezoelectric electrets, we may deduce that this structure is organized enough, like the eardrum of terrestrial mammals, to produce electrical potentials isomorphic to the surrounding acoustical waves; In a similar way, these potentials would be transmitted by GJs to the DOHC complex, and, thus, reach the phalanx of Deiters cells (i.e. the gate of the TkS). This hypothesis explains the mysterious functioning of the whales’ hearing as well as the extremely high frequencies heard by them.

- Yet there seems to be a drawback: how the electric potential could to be born, and then reach the DOHC complex, if every parts of the circuit would be bathed into a conductive medium (the sea salted water) ? Maybe, the solution is into the question : the collagen fibres are not in contact with the conductive salted water, but with the so called "acoustic fat bodies" which are electrical insulators. As such, their fatty structure would be needed by the sea medium, not essentially as a result of their acoustical properties, but instead due to their electrical and thermic insulator quality.

It is admissible that the electrical potentials resulting from the mobilisation of collagen content of the same fat bodies would be transmitted to the surrounding bone, and be conducted through GJs to the stereociliae of the OHCs as well as to the cup and phalanx of Deiters Cells. The large mysticetes, apart from their ultrasonic abilities and their use of echolocation, are able to hear bass sounds and even infra-sounds. The transmission of acoustic vibration from the surrounding sea water toward the cochlea is thought to result from a "fatty transmission".

This paradoxical data set may find the beginning of a solution, if we add an electrical "covert path" to that purely mechanical transmission. This fatty pathway is made of a very complex histological structure of adipocytes and a dense network of collagen fibres*^S^*^[48]^. This covert path would be similar to the one we describe between the collagen fibres of the eardrum and the DOHC complex, including the TkS. This system does not encompasses any operative tympanic structure. Yet, the tympanic ligament (homologous to the ancient tympanic membrane) attaches to the sigmoid process, a portion of the tympanic ring***^S^***^[49]^ .

It is accepted that there is a particular type of transmission, via perimandibular "acoustic fat bodies". These acoustic fat bodies are in direct contact with the malleus ossicle of ears, especially in odontocetes***^S^***^[50]^, but also in some mysticetes ***^S^***^[51]^. It is interesting to note that these acoustic fat bodies have an acoustic conductivity very similar to that of the sea water; Even so they are given the role of an 'acoustic lens' in the sound emission melon of delphinidae.

The structure of these “acoustic fat bodies” is very complex. They consist of layers of which each one has a specific acoustic conduction. These layers are, of course, generated by adipocytes and fibroblasts whose role must be coherent with measurement of specific “acoustic fat bodies” conductivity, which we have just evoked. Fibroblasts are the obvious source of the "dense network of collagen fibres" which are interlaced with adipocytes and fatty bodies, specific to each layer. These collagen fibres are piezoelectric like the  collagen fibres in general, regardless of their type (T I, II, ...). These piezoelectric variations of the collagen network might be transmitted to the OHCs by the GJs of the cochlea. It is very probable that this last stage of the “covert path” be very fast and very complex (the ganglion cells are much more numerous in the cochlea of marine mammals).

It is interesting to note as well that the eardrum membrane of terrestrial mammals is essentially the result of the action of fibroblasts, which are generators of collagen fibres. Otherwise, fibroblasts by inserting themselves into the network of collagen fibres, are also suitable for turning into adipocytes^[52]^, the main constituents of the lipid acoustical pathway.

Ear fat attaches to the tympano-periotic complex, inserting itself into the "triangular opening", at the entry to the middle ear, where the ear fat contacts the hammer. Thus, the "acoustic signal" transmitted by the acoustic fat bodies reaches the hammer, like the tympanic vibrations of terrestrial mammals.

The roles of the eardrum and these fatty acoustic bodies are very similar: they provide double transduction:

- Acoustic vibrations mechanically mobilize ossicles or their remains; This phenomenon is useful for the perception of low acoustic frequencies.

- Since the fibrous collagens are piezoelectric electrets, we may deduce that structure is organized enough, like the eardrum of terrestrial mammals, to produce electrical potentials isomorphic to the surrounding acoustical waves; In a similar way, these potentials would be transmitted by GJs to the DOHC complex, and, thus, reach the phalanx of Deiters cells (i.e. the gate of the TkS). This hypothesis might explain the mysterious functioning of the whales’ hearing as well as the extremely high frequencies heard by them.

Furthermore, cetaceans and bats use both the protein which fulfils the role of drain in the TkS (prestine).

Our description of the covert path should also be applied analogously to the hearing of mammals endowed with significant ultrasonic abilities (bats and sea mammals).

#### Si 08B The human fetus is in a similar situation to marine mammals. The mesenchymal environment of the ossicles present in the middle ear of the foetus is an obvious obstacle to their correct functioning.

"The early stages of auditory ossicles development all occur within the solid mesenchyme of the pharyngeal arches until the eighth month of development^S^^[53]^, then within a fluid-filled space for the final month, and finally only postnatal in the air-filled tympanic cavity of the neonate". This transition in auditory ossicle environment means that the middle ear does not function correctly until after birth"^S^^[54]^. This makes it difficult to understand the existence - though well demonstrated - of fetal hearing from the 22^th^ amenorrhea week^S^^[55]^ - ^S^^[56]^. So Hill concludes that "any prenatal conduction to the cochlea must be mediated through bone conduction".

However, the mechanism of bone conduction is itself poorly understood, so that the explanation of foetal hearing by bone conduction would simply move the problem on. It therefore seems that another conceivable mechanism would be the "covert path" as could be the case for sea mammals as well ....

Furthermore the frog "Sechellophryne Gardineri" is able to hear without eardrum, even vestigial, and its hearing organ is just its mouth^S^^[57]^.

#### Si 09 Electrical pathway

#### Si 09A Electrical links of the eardrum to the cochlea

The superior ligament of malleus (K), made of collagen I, attaches the head to the tegmen tympani. The SLM may electrically connect the central structure of the eardrum to bone conductive fibres and to the ground (the electric common return). Peripheral ends of the radial collagen fibres of the eardrum are connected to the osteocytic GJs network.

The eardrum consists of four layers (fig. S19): the epidermis (B), two fibrous collagenic layers (AA’), and an internal mucous membrane (C). The three ossicles are the stapes (F) with its muscle (J), the uncus (E), and the malleus which consists of three parts: the head (D), the lateral process (H) and the manubrium (I). The malleus can be mobilized by its muscle, the tensor tympani (G) which, when enabled, pulls the malleus medially, tensing the eardrum and damping its vibration.

| 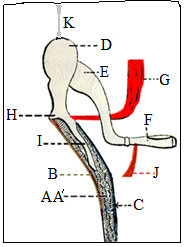 | 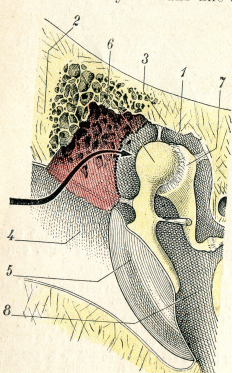 |
| --- | --- |
| Figure S24 Section of the Eardrum through the handle of the malleus***^S^***^[58]^: Schema of the Superior Ligament of the Malleus (SLM) ensuring the connection of the umbo to the electric common return (K). | Figure S25  Ligaments of the Malleus are inserted into the attic; The Superior Ligament of Malleus (SL M(3)), is inserted in the tegmen tympani***^S^***^[59]^. |

The superior ligament of malleus (K), made of collagen I, attaches the head to the tegmen tympani. The SLM may electrically connect the central structure of the eardrum to bone conductive fibres and to the ground (the electric common return). Peripheral ends of the radial collagenic fibres of the eardrum are connected to the osteocytic GJs network (fig. S19 and S20).

#### Si 09B The DOHC complex

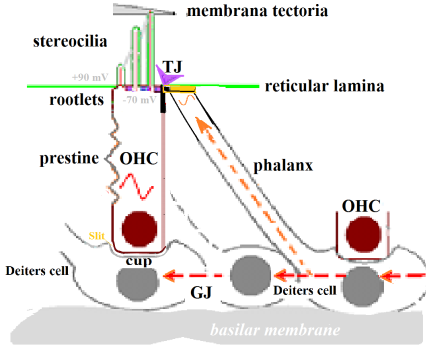

Figure S26
Schema of the DOHC complex

Stereocilia are implanted on the apex membrane of OHCs (cuticular plate / reticular lamina).
They are bathed in endolymph and mobilized by its movements (TW).

#### Si 10 Trickystor

#### **Si 10A** Source : Ionic channels and time constant

The question of the reality of the contractions at high frequencies of the walls of the OHC in physiological context is poorly understood;the cycle polarization - depolarization created by the movement of the ciliary tuft, induced by the sound, in fact involves the electric plasma membrane time constant; the value of this time constant is not compatible with frequencies of contraction up to 20 kHz in humans and 150 kHz in some bats***^S^***^[60]^. A persistent difficulty in understanding the role of electromotility in amplification is the membrane time constant; the resistance and capacitance of the membrane of an OHC cuticle would suggest that a voltage-driven process should diminish in sensitivity above a corner frequency of only 1 kHz***^S^***^[61]^ ^-^ ***^S^***^[62]^.

But, the stereociliar signal can trigger prestin electromotility at higher frequencies, provided that it is combined with a second stimulation, whose effects would not be subject to the membrane time constant***^S^***^[63] -^ ***^S^***^[64]^.

In reality, such a potential is recordable in the gap (slit) separating the DC cup from the base of the OHC***^S^***^[65]-^***^S^***^[66]^ ***^- S^***^[67]^ ***^- S^***^[68]^ ***^-S^***^[69]^ ***^- S^***^[70]^ ***^- S^***^[71]^ ***^- S^***^[72]^ (Cf. Fig. S21, DOHC5: « cup »). For high frequencies (> 8 kHz), variations of potential of the cup are rapidly transmitted to the gap by means of hemi-channels made of Cx26 and Cx30***^S^***^[73] -^ ***^S^***^[74]^ ; This being the case, the prestin could contract at the same frequencies***^S^***^[75] -^ ***^S^***^[76]^. It is also possible that variations in voltage of the Deiters cup would produce a capacitive effect, through the Deiters membrane, the cupular gap and the membrane of the base of the OHC.

This extracellular cupular signal has been attributed to electrical stereociliar phenomena, and this attribution has caused it to be confused with the "cochlear microphonic potential ". But if we refer to the prevailing opinion, according to which the microphonic potential is due, for the most part, to the mechano-electrical activity of the prestin, we are faced with a vicious circle: The contractions/extensions of the prestin would produce the microphonic and this microphonic, then, is invoked to explain the phenomena of contraction / extension of the prestin!

However, this explanation might be relevant if the initial origin of the extracellular potential were attributed to the non-OHC part of the microphonic, i.e. the pT. A fact supporting this hypothesis is the following: The DC cups themselves are the seat of intracellular***^S^***^[77]^ voltage variations (AC) with the same frequency and amplitude as the extracellular voltage***^S^***^[78]^. Variations in the intracellular voltage of the cupule of Deiters could be the result of the pT and, as such, the source of that extracellular potential, instead of its consequence***^S^***^[79]^.

The hypothesis of the trickystor, with its FET features, might indeed overcome this problem. The transconductance of an FET is not submitted to a low pass limitation; it is expressed by the ratio g = IDS/UGS (UGS is the Gate-Source voltage. IDS is the Drain-Source current). The higher g is, the greater the gain of the transistor is. This gain allows it to cancel the intensity decay which would otherwise occur because of the time constant. Its high input resistance and its low input capacitance give to a FET characteristics similar to those of a triode. The interest of such an FET lies in its excellent selectivity and its low noise factor (narrow bandwidth). which enables it to play a role of preamplifier and/or oscillator

#### Si 10B The cuticular plate bilayer

Like all cellular membranes, the cuticular membrane is a lipid bilayer (fig. 03) which consists mainly of amphiphilic lipids (generally phospholipids). They have one head group that is hydrophilic ('polar') and two hydrocarbon tails that are lipophylic ('non polar'). One tail typically is saturated, while the other tail is unsaturated with one or more cis-double bonds (conjugated or homoconjugated). Each cis-double bond creates a small kink in the tail. By forming a double layer, with the polar ends pointing outwards and the non-polar ends pointing inwards, membrane lipids form a 'lipid bilayer' which keeps the watery interior of the cell separate from the watery exterior.

The "lipid bilayer" plays a dual role in the life of the cell: both as insulator and filter.

a - Its insulating lipid molecules, arranged in a 5 to 10 nm thick bilayer, form an impermeable barrier to the passage of most water soluble molecules. They block the passage of inorganic ions (K +, Na +, Cl-, Ca2+, ...) and hinder the diffusion of polar organic solutes such as amino acids.

b - It is a filter as well: protein molecules regulate transmembrane exchanges;within itself, the membrane is permeable only to small hydrophobic molecules (O2, N2, glycerol, ...).

##### Si 10C The structure of the phospholipid bilayer is a liquid crystal (of smectic type).

The findings from the neutron diffraction work suggest that the cholesterol molecule stays at the center of the bilayer and is constrained to a maximum displacement inferior to 6 Å relative to the cellular body as a whole***^S^***^[80]^. The sterols show a strong preference for embedding themselves between bilayer leaflets, which is an unusual placement for them ***^S^***^[81]^.

Yet cholesterol has a poor affinity for phospholipids containing PUFAs and that enhances its mobility. This may affect the apical, as well as the lateral or basal, distribution of cholesterol. OHCs have an inhomogeneous distribution of free cholesterol with higher concentrations in the basal and apical membranes and lower levels in the lateral wall (***^S^***^[82]^ ***^- S^***^[83]^).

There are also quantitative differences in lipid lateral mobility of cholesterol, and rafts, among the apical, lateral, and basal regions of the OHC (Fig. S27).

| 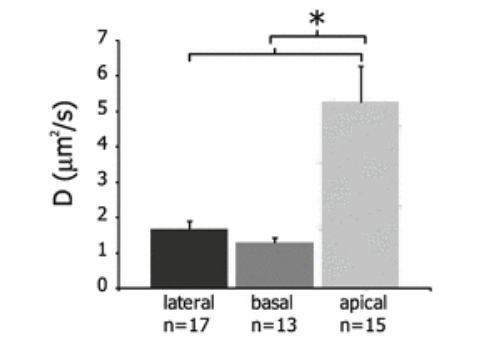 | Effective diffusion coefficients, D (µm²/s), across the three OHC regions : D values in the apical region were significantly larger (p<0.0005) than in lateral or basal regions. The "membrane fluidity" in the membrane plane is all the more important that the fatty acids of membrane lipids are unsaturated. The fluidity of the cuticular membrane of the OHC is an indication that it contains many more PUFAs than the other parts of the OHC’s cellular membrane |
| --- | --- |
| Figure S27 (Excerpted from Haroun ^S[84]^) | |

.

#### Si 10D Semi-Conductors of the TkS

The hearing is impaired by genetic disorders of the conjugated and homoconjugated (fig. S24 below) PUFAs semiconductors (Role of peroxisome)***^S^***^[85]^. Peroxisomes are [organelles](https://en.wikipedia.org/wiki/Organelle) found in all [eukaryotic](https://en.wikipedia.org/wiki/Eukaryotic) cells. There are at least 32 known peroxisomal proteins that participate in the process of peroxisome assembly. Their deficiency gives impairment hearing of high frequencies and they are involved in [catabolism](https://en.wikipedia.org/wiki/Catabolism) of [very long chain fatty acids](https://en.wikipedia.org/wiki/Very_long_chain_fatty_acid), [branched chain fatty acids](https://en.wikipedia.org/wiki/Branched_chain_fatty_acids), [D-amino acids](https://en.wikipedia.org/wiki/D-amino_acids), and [polyamines](https://en.wikipedia.org/wiki/Polyamine), [reduction](https://en.wikipedia.org/wiki/Redox) of [reactive oxygen species](https://en.wikipedia.org/wiki/Reactive_oxygen_species) – specifically [hydrogen peroxide](https://en.wikipedia.org/wiki/Hydrogen_peroxide) – and biosynthesis of [plasmalogens](https://en.wikipedia.org/wiki/Plasmalogens), i.e. [ether phospholipids](https://en.wikipedia.org/wiki/Ether_phospholipid).
Peroxin (or peroxisomal/peroxisome biogenesis factor) is a protein found in peroxisomes.

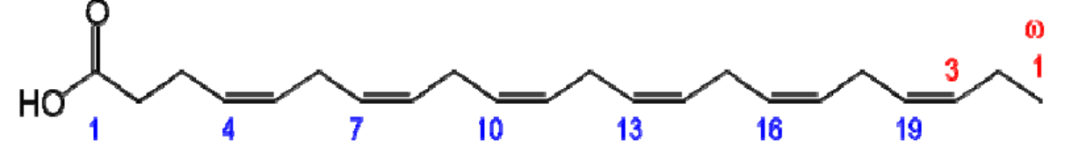

Figure S28

DHA is an example of long chain homoconjugated fatty acid.

#### **Si 10E** The electrical as well as mechanical state of DCs drives the electromotility of the OHCs

The separation of a DC phalanx from a related OHC cuticle, or of a cupule from the OHC that it supports (or the elimination of GJs between the bodies of Deiters cells) results in deterioration of the electrical behavior of the concerned OHCs (Yu & Zhao, 2009). There being no electrical synapse (GJ) between DCs and OHCs, it has been proposed that this electric effect on the OHC is due not to an electrical interruption between DCs and OHCs, but rather to an interruption of a purely mechanical nature.

Stimulation of the DCs, either electric or mechanical, can modulate the electromotility of the OHCs (Yu&Zhao, 2009) and in vivo active cochlear amplification depends on GJs between the DCs. Cx26 expression in the cochlear supporting cells plays a critical role in active cochlear amplification and its targeted deletion can eliminate active cochlear amplification.

It is coincident with a large reduction in distortion product otoacoustic emission (DPOAE) and severe hearing loss at high frequencies (changes are greater in the shortest OHCs).

#### **Si 10F** Gate ; Field effect of Deiters Cells on the cuticles of OHCs

The **tunnel field-effect** transistor (TFET) is an experimental type of transistor. Even though its structure is very similar to a metal-oxide-semiconductor field-effect ([MOSFET](https://en.wikipedia.org/wiki/MOSFET)), the fundamental switching mechanism differs, making this device a promising candidate for [low power electronics](https://en.wikipedia.org/wiki/Low_power_electronics). TFETs switch by modulating [quantum tunneling](https://en.wikipedia.org/wiki/Quantum_tunneling) through a barrier instead of modulating [thermionic emission](https://en.wikipedia.org/wiki/Thermionic_emission) over a barrier as in traditional MOSFETs. Because of this, TFETs are not limited by the thermal [Maxwell–Boltzmann tail](https://en.wikipedia.org/wiki/Maxwell–Boltzmann_statistics) of carriers, which limits MOSFET drain current [subthreshold swing](https://en.wikipedia.org/wiki/Subthreshold_slope) to about 60 mV/[decade](https://en.wikipedia.org/wiki/Decade_(log_scale)) of current at room temperature (exactly 63 mV/decade at 300 K^[1]^). The concept was proposed by Chang et al while working at IBM ^[2]^. Joerg Appenzeller and his colleagues at IBM were the first to demonstrate that current swings below the MOSFET’s 60-mV-per-decade limit were possible.

According to Yu and Zhao (2009), the electric effect of DCs on an OHC must be assigned to an intermediate phenomenon of a mechanical nature. Yet there is an electrostatic link (a field effect) between the cytoskeleton apex of the DCs phalanxes and the OHC cuticle. This link exists without a significant current, and so without electrical conduction by gap-junctions. That *ephaptic effect*^[86]^ is the field effect of a biological electric conductor, for example the field effect of an axon on its neighbourhood. It is distinguished from the phenomena of simple diffusion in the extra-cellular environment; It is also distinguished from communications between neurons by means of synapses either chemical or electrical ***^S^***^[87],^ ***^S^***^[88],^ ***^S^***^[89]^.

It is clear that this field effect must be detected mainly by the effects it produces within the cuticular bilayer, where it likely acts as a gate on the communication between the set source {stereocilia / endolymph} and the set drain {OHC cytoplasm, prestin, baso-cellular membrane, cupular slot}.
 Indeed, the field effects in the DOHC suggest that it could operate in a way similar to a triode***^S^***^[90]^. A Field Effect Transistor (FET) uses an electric field applied by a gate, to control the conductivity of a "channel" in semiconductor materials. Applied voltage on the gate electrode controls the amount of charge carriers flowing through the system.

In physiological conditions, lipids do not cross TJs: There is a strict insulation, chemical as well as electrical, between the two bound cells, and no current will flow from one to the other of their apical membranes***^S^***^[91]^***.*** For this reason, no electric current can cross the barrier between the phalangeal apexes of the DCs and the cuticle of the embedded OHC. However, this border cannot prevent a hydrophobic intercellular electrostatic coupling unrelated to any GJ^S^***^[92] -S^***^[93]^.

#### Si 10G The destruction of the cytoskeleton annihilates the electric effect of the DCs on the OHCs

The apex of the phalanxes of the DCs is connected to the cuticular plate of OHCs by Tight-Adherens Junctions (TAJs). These TAJs bring into contact the cytoskeleton of adjacent cells. It would be interesting to design “*experiments that could separate changes in intracellular ion concentration, from mechanical resistance by the cytoskeleton* **^S^**^[94]^ *as a step towards understanding the signal transduction pathways that might involve microtubules*” (Szarama, pers.com., 2012). Furthermore, the destruction of the cytoskeleton of the DCs negates the effect of electric stimulations of the DCs on OHC electromotility (Yu and Zhao, 2009).

This implies that the cytoskeleton of DCs plays a critical role in the modulation of OHC electromotility. According these authors, this is evidence that, if electrical stimulations of DCs influence OHC electromotility, it must be through the DC-OHC mechanical coupling rather than by extracellular field effect. However, the electric voltage of the phalangeal apex necessarily causes variations of the electric field in the OHC cuticle (via the intercellular border at the level of the TJ). Indeed, the destruction of the phalangeal cytoskeleton removes the electrical activity of the phalangeal cytoskeleton***^S^***^[95]^ and, as a consequence, its field effects.

The measurements used by Yu and Zhao (2009) were taken using the whole-cell patch clamp method, i.e. with electrodes implanted into the cytoplasm of the DC and the OHC, but not within their respective bilayers; Thus, weak capacitive interactions, consistent with the action of an FET gate, escaped measurement.

Cupular phenomena represent a more sensitive issue. The {DC Cup /OHC base} junction is very unusual (see below S24 - S25): at the level of the cupule it includes hemi-channels, whose activity is confined in the cupular slot; It is an extremely small dedicated space, without obvious communication with what is generally regarded as the extracellular medium or the Nuel space.

| 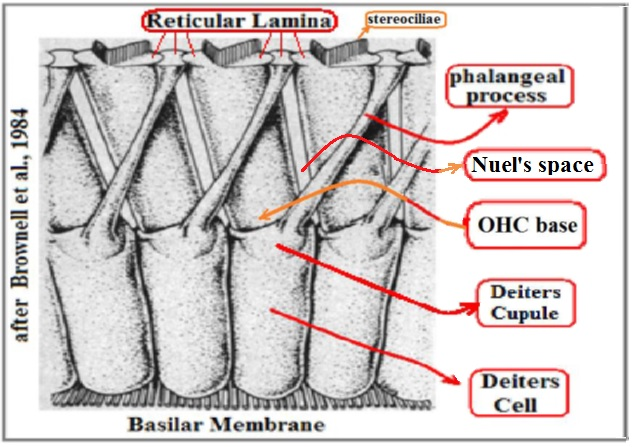 | 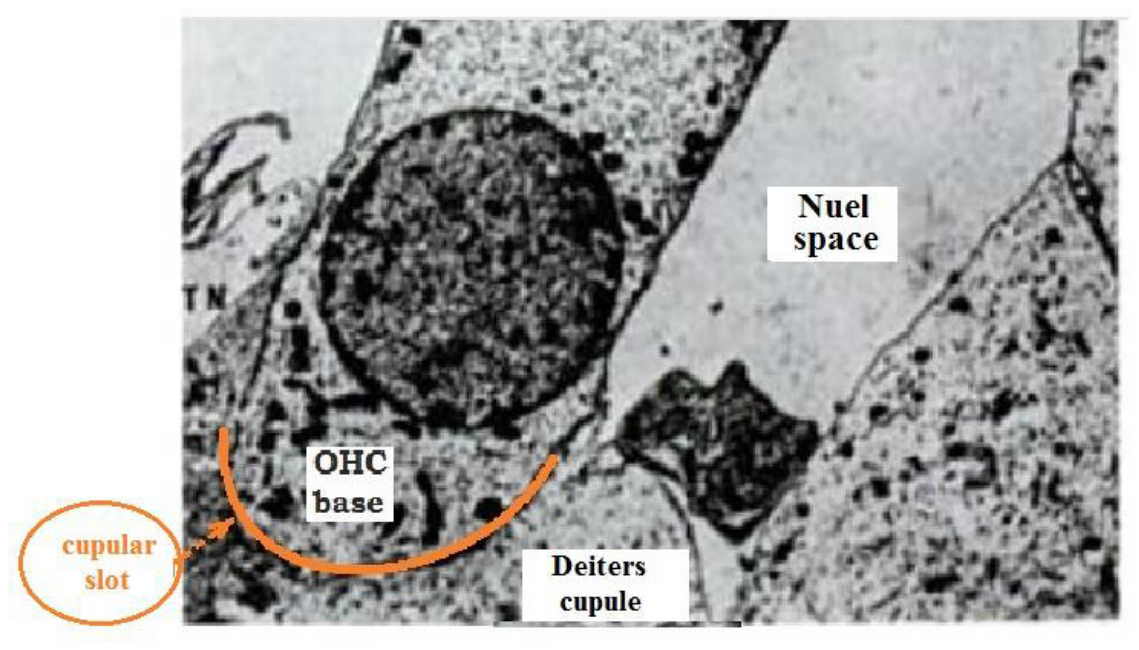 |
| --- | --- |
| Fig. S29 After Brownell (1984) | Fig. S30 Adapted from Coujard R., Poirier J., Précis d'Histologie humaine p728 (1980) |

Yu and Zhao (23) have put the extracellular milieu into communication with the earth, probably allowing effective control of the extra cellular environment after the breakdown of the cupule-OHC junction. This control is irrelevant, however, as regards variations affecting the slot between the cupule and the OHC as a specific space. The same is true for the variations within the intra-membranous TAJ cuticle-phalanx space^S^^[96]^.

In another experiment, these authors blocked the ionic channels of the OHC basal membrane in such a way that the ionic changes in the cupular slot could not elicit charge-carrier exchanges between OHC and cupule. But, of course, when the physical joining between the DC cup and the OHC base is preserved, this precaution does not prevent the intervention of a capacitive electrical signal between cupular and OHC membranes. So, the blocking of ion channels is not sufficient to eliminate all of the effect of the Deiters cupule in the OHC membrane.

The OHC and the cupule are joined at the membrane level. The two membranes act as capacitors, and mechanically severing the coupling between them changes the capacitance of the set radically. In the case of the separation between cupule and OHC, the cupular slot enclosure is totally open and allows a fast diffusion of ions from the cupular hemi-channels: it follows that ionic changes from the cupule get "lost" in the extracellular medium and cannot reach a level sufficient to act upon the OHC.

In each phalanx, the micro-tubules set, might manage an electrical resonance which would optimize the tuning.

When a potential (eg pT amplified and filtered by phalanxes) is applied to the bio-organic semi-conductors of the cuticular bilayer, a current can flow through this cuticular bilayer, between the suitably biased source and drain "electrodes".

#### Si 10H The Drain: Prestin

There is good evidence that prestin has undergone adaptive evolution in mammals ^[^***^6^***^]^. This is associated with acquisition of high frequency hearing in mammals.^[^***^7^***^]^ The prestin protein shows several parallel amino acid replacements in bats and dolphins that have independently evolved ultrasonic hearing and [echolocation](https://en.wikipedia.org/wiki/Animal_echolocation), and it represents a rare case of [convergent evolution](https://en.wikipedia.org/wiki/Convergent_evolution) at the sequence level. Unlike the classical, enzymatically driven motors, this new type of motor is based on direct voltage-to-displacement conversion and acts several orders of magnitude faster than other cellular motor proteins. A targeted [gene disruption](https://en.wikipedia.org/wiki/Gene_knockout) strategy of prestin showed a >100-fold (or 40 dB) loss of auditory sensitivity***^S^***^[97]^.

The piezo tympanic signal reaches a DC_5_ cup, which supports the OHC in question. DC_5_ is distinct from the four other DCi (1..4), the phalanxes of which are in contact with the cuticle of the OHC. At the same time, the pT signal is carried by the microtubules of the phalanxes of the four DC_i(1..4)_ toward the TJs. There is no electrical conduction between the phalanxes and OHCs, but there is a capacitive effect (Scut) between phalanxes and the intra-cuticular membrane space. The cuticular bilayer comprises a great many homo-conjugated PUFAs which are semiconductors. This property makes the phalangeo-cuticular structure similar to a FET (*Trickystor)*. The characteristic frequency of the DOHC complex is inversely related to its distance from the oval window (Greenwood relation), the length of the phalanxes of the DCis, the length of the OHC and the length of stereociliae.

For the classical pathway, the duration of the mechanical signal path increases with the distance between the eardrum and the Greenwood area corresponding to a given frequency. For the hidden pathway, the duration of the electrical signal path increases with the length of the phalanx, and depends on characteristics of the cytoskeleton***^S^***^[98]^. If the Scil (ciliary signal) and the S_cut_ are approximately synchronous (d ≈w), this results in an amplified mechanical signal (S_amp_). If there is a precession either of the S_cut_ signal on the S_cil_ signal, or of the S_cil_ signal on the S_cut_ signal, the *trickystor* does not immediately acquire an maximal effectiveness since this effectiveness requires synchronous interaction of both signals.

But, the acoustic signal (TW) is transduced into an electrical signal by the stereocilia; this electrical signal is amplified by the *trickystor*; then it is once more transduced into a new overamplified mechanical signal by prestinic movements. These movements act on the basilar membrane and the supra-reticular region in such a way that the local amplitude and accuracy of the TW are enhanced.

The wavelength depends on the velocity of propagation of the wave in the medium it crosses. When the wave passes from one medium to another, in which its velocity is different, its frequency remains unchanged, but its wavelength varies (Dic. Phys.). As a result, this feedback process could set up a phase lag***^S^***^[99]^ (τfb), increasing with each cycle. The increasing phase lag would tend, by progressive desynchronization, to self-limitation, damping and extinction of this type of amplification (fig. S31).

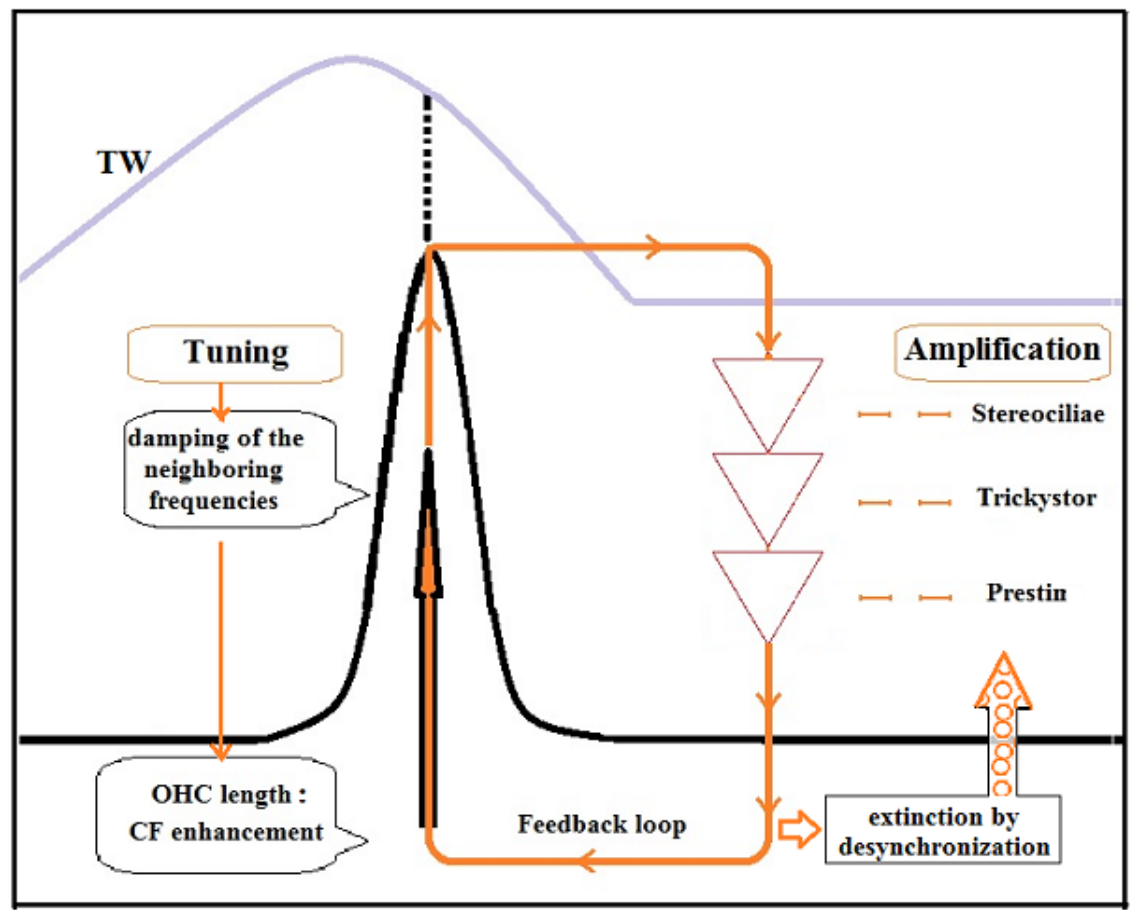

Figure S31
Schematic view of some putative processes regarding
amplification, tuning, damping by DOHC and also creation of OAE.

#### Si 10I About prestin and low frequencies

The conductance of the Mechano Electrical Transducer channels changes are depending upon the tonotopical position within the cochlea, and that suggests differential requirements at different frequencies^S[100]^.

Oddly enough, some profoundly deaf people are able to hear some so-called infra-sound frequencies***^S^***^[101]^ ***^- S^***^[102]^ through the labyrinth (sacculus). The cochlea of normally hearing people has mechanisms that weaken or suppress these sound elements before they are transmitted to the brain. Ashmore, points out that “prestin is expressed […] surprisingly, in the cells of the vestibular system***^S^***^[103]^”.

Thus, into its vestibular locus, prestin (otherwise the motor protein of OHCs), might increase or decrease the sound amplitude according to whether the frequency is above or below 50Hz***^S^***^[104]^.

It is possible that this "suppressive" effect would be produced by a phase inversion during the amplification process of these infra-sound frequencies, thus producing an attenuation of these stimuli either when they are not useful, or would become disruptive.

### **Si 11** Phase and Times

#### Si 11A Phase shift of the TW and pT for f > 3 kHz (Quix)

#### Discussion about the phase lag of TWm relatively to the without delay signal of the pT:

The princeps (originator) TW of Von Bekesy, derived from ossicles, retains the frequency of the acoustic stimulus but with a delay resulting from the speed of the sound waves in water (1.500 m/s)***^S^***^[105]^, whereas the shorter delay of the pT results from the speed of the electric signal through the GJs (ie, approximately, the speed of an electric signal in salt water, i.e. a speed of 226 000 m/s) ***^S^***^[106]^. The pressure variation on the eardrum, sine-type (TW), is transduced into electrical potential by the stereocils ("source" of the TkS).

Another sine of the same origin and frequency produces a pT potential and reaches the "gate" of the TkS. Since the movement of TW is slower than the movement of the pT electrical signal, there is a phase shift between these two waves that may tend to lessen the effect of the TkS.

To evaluate the effect of this phase shift, we can take two examples. Let us use for this evaluation 4 khz and 5 khz (Table S09 below).

| **Effects of phase shift on 4 kHz**.  Calculation of the time delay for f = 4 kHz | **Effects of phase shift on 5 kHz.**  Calculation of the time delay for f = 5 kHz |
| --- | --- |
| Be the delay of TW: D1_4_.  If a 4 kHz acoustic stimulation is used, the TW must travel 8.7 mm, ie 0.0087 m.  To traverse this distance at the speed of a sound in water, i.e. 1,500 m/s, the delay will be 🡪 D1_4_ = 0.0087/1500 = 0.0000058 = 5.8 * 10^-6^ sec.  D_pT_ being the D2_4_ time of the pT potential:  Delay of pT to reach the same zone 0.0044 m, at the speed of the electric wave in salt water (roughly identical to the speed via the Gap junctions) = > 0.0087/226 000 = 3.85 * 10^-8^ sec.  For 4 kHz, the difference dD_4_ = D1_4_-D2_4_ = 5.8 * 10^-6^ sec-3.85 * 10^-8^ sec;  D2_4_ represents 6 thousandths of D1_4_ and can be considered negligible because the value of this delay is included in measurement errors or calculation rounds and can be accepted.  dD = D1-D2 ± ε = 5.8 * 10^-6^ ± ε sec.  The phase shift at 4000 Hz : The period of a wave of 4000 Hz is T = 1/4000 = 0.00025 sec.  Δ (ϕ) = 5.8 * 10^-6^ * 360/0.00025 = 8° 35 = 0.15 rad  In percent, 0.15 radian/6.28 Radian = 0.02 (i.e. 2% of a cycle).  **Generalization** : if f2>f1 the travel of f2 will be shorter than that of f1(Greenwood relation). The shift will be thus all the more small as F will be large. In fact the calculation shows that the phase shift of TW and pT for f > 3 kHz is < 3% and therefore should not interfere with the operation of the TkS. | Be the delay of TW: D1_5_  If a 5 kHz acoustic stimulation is used, the TW must travel 6.3 mm, ie 0.0063 m.  To traverse this distance at the speed of a sound in water, i.e. 1,500 m/s, the delay will be 🡪 D1_5_=0.0063/1500 = 0.0000042 = 4.2* 10^-6^ sec.  D_pT_ being the D2_5_ time of the pT potential:  Delay of PT to reach the same zone 0.0063 m, at the speed of the electric wave in salt water (roughly identical to the speed via the Gap junctions) = > 0.0063/226 000 = 2.79 * 10^-8^ sec.  For 5 kHz, the difference dD_5_ = D1_5_-D2_5_ = 4.2 * 10^-6^sec - 2.79 * 10^-8^ sec;  D2_5_ is smaller than 7 thousandths of D1_5_ and can be considered negligible because the value of this delay is included in measurement errors or calculation rounds and can be accepted.  dD = D1 ± 0.007 = 4.2 * 10^-6^ ± 0.007 sec.  The phase shift at 5000 Hz : The period of a wave of 5000 Hz is T = 1/5000 = 0.0002 sec.  Δ (ϕ) = dD * 360/T = 4.2 * 10^-6^ * 360/0.0002 = 7°56 = 0.13 rad  As a percentage, 0.13 radian/6.28 radian = 0.02 (ie 2% of a cycle). |
| Table S16 Effects of phase shift according frequency | |

#### **Si 11B** Precession of the electric pathway on acoustical pathway ?

A more specific work on the phase shift between the vibratory movements of the Tectoria Lamina (TL) and the basilar membrane (BM) has determined that the vibration of the BM is leading that of the TL^S^^[107]^. Yet, the movements between the top and the bottom are in phase opposition.

Note that for the most part the movements have reverse phase between the top and bottom except for the Best Frequency. There is evolution so that it synchronizes with the best frequency.

It is difficult to draw clear lessons from this.

We believe that the electrical signal is faster than the mechanical signal and that (despite a delay in the phalanx) this phase shift suggests a causal series of the type:

pT → GJs →Deiters → Phalanx (+delay) → Gate of TkS → signal between Reticular Lamina and Prestin
→ Prestin contractions → BM and LR in phase opposition
→ evolution (by resonance) towards synchronization at the best frequency.

The difference in delay between TW and pT implies a precession of the onset of the pT wave in relation to the TW and the stereociliar potential it produces. Fidberger et al.***^S^***^[108]^ had discovered that: "Electric potentials inside Corti's organ preceded BM velocity" [...] And they went on "If such an obvious phase lead were present also at the start of the stimulus, it would mean that electric potentials were produced before BM motion". But unsatisfied by that result incompatible with the accepted design, they oddly enough modified their experimental protocol and, "using a 3 kHz stimulus at 100 dB SPL", they saw that the TW was moving before that "*detectable"* potentials could be generated. Indeed, the 3 kHz frequency they chose is included within the limits under the Quix frequency and does not conflict in any way against our own view.

#### Si 11C About the Quix frontier between low and high frequencies

Is there any possibility of defining a boundary between these two sets ?

The frequency of Quix (3 kHz), concerns the minimum of the pT curve, the minimum of the iso-sonic curves, and many other phenomena of the physiology of human hearing.

With regard to humans, it is logical to use the frequency of Quix which, for greater rigor should be provided with a ± Δf (1 Khz?).

**Quix = 3 khz ± Δf = 3 ± 1 khz**

The study of other mammals invites one to consider that Quix should be applied to the animal species under
consideration (cetaceans; bat, etc.) whose "Quix area" is much higher.

The greatest sensitivity of cetaceans lies between 40 and 80 kHz. Their sensitivity decreases between 40 kHz and 1 kHz, and also between 80 kHz and 150 kHz :

***cetaceans* f_QUIX_ = f_QUIX_ ± Δf = (60 ± 20) kHz**

#### Si 12 OAE versus BEW

The existence of OAEs (Oto Acoustic Emissions) has led to the postulation of a "Backward Travelling Wave" (BTW). Until now, however, no clear evidence has been found for it ***^S^***^[109]^.

The process of the Electrically Evoked Otoacoustic Emissions (EEOAE), includes electrical stimulation of the OHCs. The effect of this cochlear stimulation is extremely rapid and its upper cutoff frequency is > 80 kHz^S[110], S[111]^. This permits the recording of an acoustic response of the same frequency at the level of the external auditory conduit***^S^***^[112]^.

Under the postmortem condition, the electrically evoked basilar membrane (BM) vibration almost disappears while **the EEOAE shows no significant change**. These results indicate that the BM vibration is not involved in the backward propagation of the EEOAE. To explain this, it has been proposed to accept the existence of fast "backward compressional sound waves" instead of the elusive “slow backward traveling waves” ^S^^[113]^. Yet, that hypothesis is puzzling because it is difficult to figure out how this acoustical compressive phenomenon might be attributable to the motility of the OHCs given that, after death, the OHCs would cease to function actively.

We have shown that electrical potentials that we measured near either the eardrum or the mastoid cannot be explained solely by the diffusion of the cochlear microphonic. But if a partial diffusion of this electrical signal does exist, this ‘backward electric wave' (BEW) may cause slight acoustic vibrations of the eardrum by inverse piezoelectric effect. Thus, this BEW could be responsible for the EEOAEs and, what is more, of all OAEs !

#### Si 13 The global system using together the overt and covert paths

The whole system (overt path and covert path together) constitute a complex system which is a biologic Micro-Electro-Mechanical-System, similar to an industrial MEMS. This acronym MEMS is interesting since these few letters include: the mechanical point of view (or mechanical aspect), very present in our model, the piezo-electric aspect of collagen and the electronic aspect of the trickystor (with its biological semi-conductors); the term "micro" is perfectly justified since the size of several of its components are of the micrometric order.

**Si 14** **Notes Concerning Statistical Analysis**

The table of raw data is available as an excel file: raw_data_covert_path.xlsx (<https://figshare.com/s/3338464a25212da726cf>)

###### **Si 14A Description of the data**

We did 221 experimental series of microvoltage measurements in function of the sound frequencies (Hz) and the sound level (dB). These experimental series were measured on 62 individuals (A01-A10, B11-B20, C21-C30, D31-D35, E43-E44, E48-E50, F1-F12). Of these recordings, 110 correspond to a male and 111 to a female. The ages of individuals were < 93 years^S^^[114]^. From these 221 series of measurements, 79 were obtained at the level of an eardrum. To check the properties of collagen fibers different from those of the eardrum, we also made these measurements on the forearm, the Achilles tendon, the nasal cartilage and especially, in a very systematic way, the ligaments patellar right and left: 123 series of measurements were obtained at the level of a knee. For eardrums 40 were the left one, and 39 the right one. For the knees, 60 were a left knee and 63 a right one.

16 specific sound frequencies were globally tested (125, 200, 400, 500, 800, 1000, 2000, 4000, 6000, 8000, 10000, 15000, 16000, 20000, 25000, 30000 Hz), but 9 main frequencies were more particularly considered (500, 1000, 2000, 4000, 8000, 10000, 16000, 20000 Hz). We also tested 5 sound pressure levels from 55 to 80 dB (55, 60, 65, 70 and 80 dB). The data table is not complete, and there are missing data for some frequencies.

###### **Si 14B Body Sites of measurements regarding piezoelectricity**

###### **Si 14Ba Forearm**

###### Piezoelectric response of collagen at forearm level (μV / kHz)

For exploratory purposes, we measured synchronous potentials responding to sounds of different frequencies on four subjects, between two points of the left forearm, according to the axis of the limb, (Table S10; Fig. S27).

###### **Si 14Bb Achilles tendon**

| Vsp=2.5V | Subject D39 |
| --- | --- |
| f (kHz) | V (µV) |
| .2 | 3.2 |
| .5 | 2.6 |
| 1 | 4.0 |
| 2 | 4.7 |
| 4 | 7.1 |
| 8 | 11.9 |
| 10 | 14.8 |
| 12 | 15.9 |
| 15 | 19.7 |
| 20 | 29.8 |

Table S11

**Achilles' tendon**

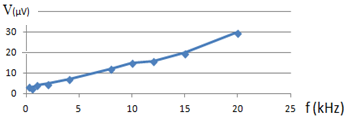

Figure S28

We used the lock-in amplifier to measure synchronous potentials V (μV) of the Achilles tendon in response to the acoustic stimulation of frequency f (kHz), emitted by a Harman/Kardon Loudspeaker: DP/N 0865DV. It was stimulated by a potential Vsp of 2.5 V, according to an installation modelled on that which we have described for the fibres of the eardrum (Table S11).

Si 14Bc Knees

As was the case for the Achilles tendon, we used a lock-in amplifier to measure synchronous potentials V (μV) of the patellar tendon in response to an acoustic stimulation of frequency f (kHz), emitted by a Harman/Kardon LoudSpeaker: DP/N 0865DV stimulated by the VLS (Volts) potential. The magnitudes of the synchronous potentials (μV) recorded on the patellar tendon, in function of the acoustic frequency, are reported below for different subjects (Table S12 and Fig. S29).

| **f (Hz)** | **D36** | **D37** | **D38R** | **D38L** | **D39** | **Total** |
| --- | --- | --- | --- | --- | --- | --- |
| **100** | 14 | 5 | 8 | 8 | 8,5 | 43.5 |
| **500** | 12,5 | 4,2 | 6 | 6 | 6,3 | 35.0 |
| **1000** | 15,5 | 5,8 | 7 | 6,1 | 8,5 | 42,9 |
| **2000** | 21 | 9,1 | 8,4 | 7,6 | 10,4 | 56,5 |
| **4000** | 27 | 13,4 | 10,9 | 9,8 | 16,5 | 77,6 |
| **8000** | 42 | 23,7 | 17,1 | 14 | 26,7 | 123,5 |
| **10000** | 54 | 28,4 | 20,5 | 17 | 32,5 | 152,4 |
| **15000** | 73 | 42 | 27,2 | 22 | 47 | 211,2 |
| **20000** | 92 | 61,4 | 35 | 29 | 62 | 279,4 |
| **Total** | 351 | 193 | 140,1 | 119,5 | 218,4 | 1022 |
| **age** | **30** | **51** | **73** | **73** | **17** |  |
| Table S12  Speaker LVL 2.5 V ; F (Hz) ; responses (µV) | | | | | | |

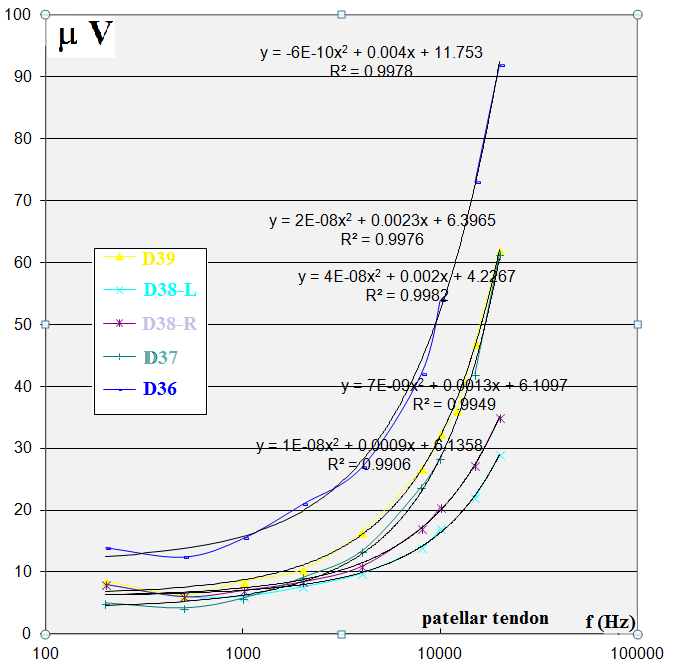

Figure S29

VLS = 2.5 V. Synchronous voltage vs acoustic frequencies for 5 patellar tendons. Every R² > 0.99

Every musculo-tendinous formation that we tested (mastoid, arm, Achilles tendon, patellar ligament) produced electric potentials synchronous to the acoustic stimuli. The piezoelectric effect sheds some light on this phenomenon.

**Si 15 Effect of pathological history on measures**

Regarding eardrums, we noted some phenomena related to pathological histories;

For example, weakness of piezoelectric responses in the case of allergy inside the ear canal (B16, B17). Anomaly of measurements carried out on a scar zone (D31 on the left eardrum).

D33 uses hearing aids "PS60 Resound" on both ears. Measures were made without any of the two prostheses present. However, the electrical response is particularly high on the eardrum G of this subject. In this case, would the histological structure of the eardrum had evolved by eliciting a more important piezoelectric response, so that the deafness would be more or less compensated ?

**Si 16 Pain felt during the measurement taken on the eardrum**

Regarding  the 79 eardrum series, an index of "*felt pain*" (0 to 5) was recorded. 51 recordings describe no pain (0) and 12 a gene or major  pain (between 3 and 5). Sometimes the placement of the probe was too painful to result in a measure concerning the ear {A07 / D34}.

**Si 17 Explaining fixed effects by characteristic variables of the individuals**

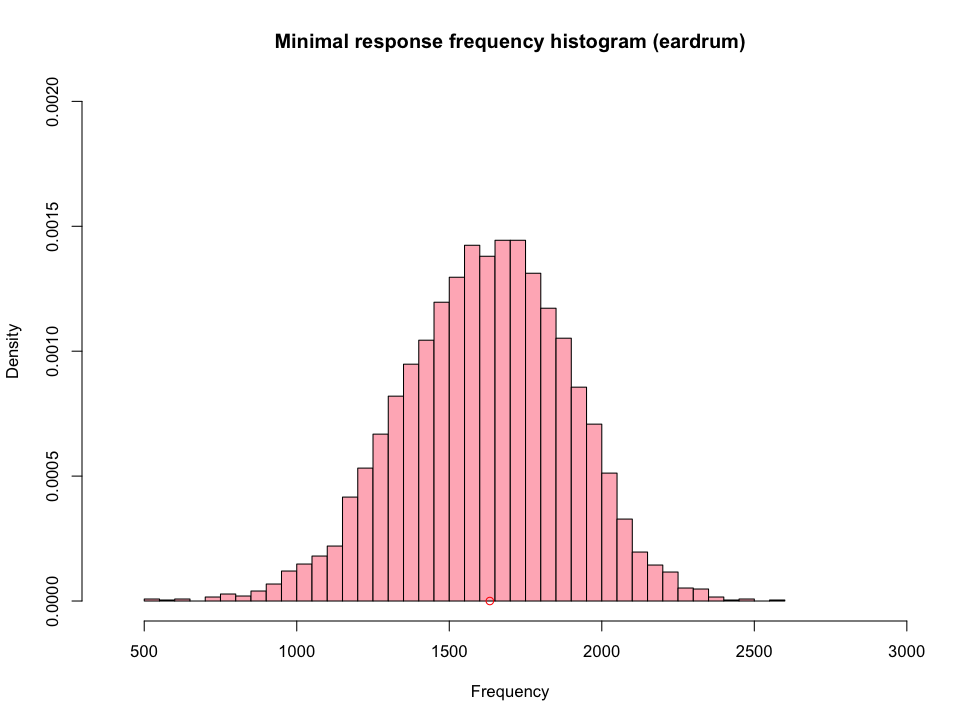

Figure S30
Distribution of the minimal frequency value for eardrum data.

For eardrums the minimal frequency (Quix frequency small red circle in Fig.S30) is lower than the knees one : It appears that separate estimates of the log-quadratic model in the frequencies F for the two sound levels 70 and 80 dB lead to two distinct minimal values 2791Hz and 3375Hz (Fig. S31). Contrary to what we observe for eardrums, it seems that there is for knees a significant interaction effect between the sound frequency and the sound level.

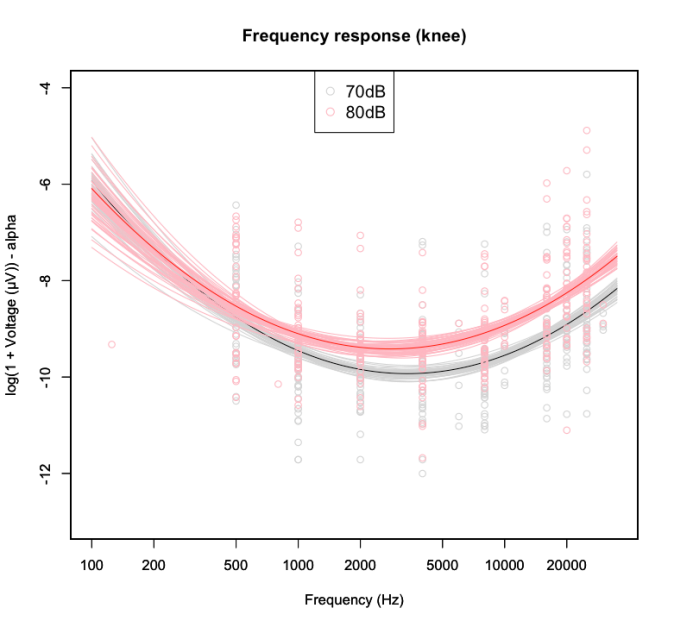

Fig. S31 Effect of the amplitude of the sound stimulus for knee measurements

#### **Si 18** Comparison of the Wegel curve of the acoustic pressures, versus the curve of the piezoelectric tensions (pT)

For all the individuals, and for the two sides, the minimum value of the curve is obtained for a frequency equal to about f_pTmin_ =1632 hz. The frequency f_QUIX_ ≈ 3000 hz (fig. S32)would be the frequency which marks the region of maximum audibility, i.e. for which there is the lowest threshold threshold***^S^***^[115]^ ***^- S^***^[116]^ (hearing *'yellow spot'* of Quix). These two minima (1632 hz and 3000 hz) are relatively different. Yet we may see that f_QUIX_ (3000 hz) ≈ 2 f_pTmin_ (1632 hz); ie f_QUIX_ is approximately the harmonic of f_pTmin.._

**f**_QUIX_ ≈ 2 **f_pTmin_**

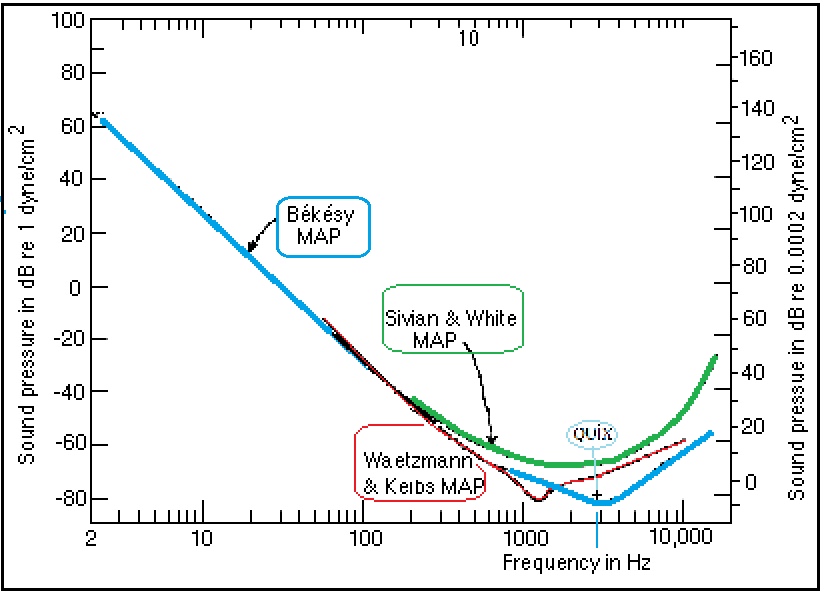

Figure S32
Wegel diagram and other classical schemas excerpted and modified from
Licklider, Handbook of Experimental Psychology, S.S. Stevens, ed., 1951

The frequency of Quix seems to be less fixed than its author thought. The frequency of Quix is that which is audible by the lowest acoustic stimulation compared to the other frequencies. In our experiments, the stimulation of the eardrums used relatively high amplitudes in relation to the threshold of hearing. As a result, the frequency of the minimum of the pT curve is lower than the frequency of Quix, but the two frequencies may be closer, or coincide, if we use liminal acoustic stimuli, in both cases (acoustic and piezo-electric).

The Wegel curve and the many related curves of the physiology of hearing give the minimum pressure required for each frequency to be heard. The Piezo-electric potentials depending on the frequency, have a variation that seems homogeneous to the physical pressures (dB SPL) of the audiometric thresholds that one has in the curve of Wegel. If the minima coincided one could say - depending on the frequency - that the less acoustic energy is necessary to reach the audibility, the less the piezo-electric response is high. The more acoustic energy is needed, the more the piezo-electric response is high!

#### Si 19 individual effect

ICI 2

**inclure ici stats:**

**Age**

**BMI**

**droite/gauche**

## **Si 20**

#### **Si 20A** Judicious redundancy

We think that multiple channels enhance the quality of information received with respect to the information emitted: If the transmission consists of several channels contaminated by radically different noise sources, and if, upon reception, you retain only what is common to these different collected signals (judicious redundancy), the result might be a cleaner signal, partially freed of parasitic noise***^S^***^[117]^.

#### Si 20B Vicariances^[118]^

Moreover, the deficiency of one of the pathways might be partially compensated by the vicariance of the other (or other) pathway(s). Hearing simultaneously uses a mechanical transmission (acoustical TW) and electric transmission (pT sent to the trickystor). The noise polluting the overt path (mechanical transmission by ossicles, TW and mobilization of the stereociliae), is different from the noise polluting the covert path (piezoelectric activity of the tympanum, electrical signal transmitted through GJs, functioning of the trickystor). If we select what is common to the outputs of each of these two pathways, we can reduce the specific noise of each of the two.

The multiplicity of channels used to transmit the acoustic information, even if some of these channels are impeded ***^S^***^[119]^, is beneficial for an optimal hearing. This multiplicity allows some parts of the signal to reach the neurological structures with sufficient practical value.

In addition, the electrical signal itself, is transmitted using a very large number of pathways, isolated from each other, so that their final summation reinforces what is common (signal) and weakens their differences (noise).

### **Si 21** Supplementary Information Bibliography

1. Gavilan C., Sanjuàn J., Microphonic Potential picked up from the human tympanic membrane. *Ann Otol Rhinol Laryngol.* **73,** 102-109 (1964).
2. Auriol B., Béard J., Broto J.M., Descouens D., Procédé et dispositif de mesure d'une tension électrique relative à une fibre de collagène pour l'aide au diagnostic par un praticien en vue de l'identification d'une éventuelle altération et de l'évaluation de la qualité fonctionnelle de la fibre de collagène", *French patent application* n° FR11/54672 (2011).
3. Gavilan C., Sanjuàn J., Microphonic Potential picked up from the human tympanic membrane. *Ann Otol Rhinol Laryngol.* **73,** 102-109 (1964).
4. Tasaki I., Davis H., Legouix J., The space time pattern of the cochlear microphonics (guinea pig) as recorded by differential electrodes, JASA **24**, 502-18 (1952).
5. Offutt G., *The electromodel of the auditory system*, Golo Press, 23-24 (1984).
6. Mammano F., oral discussion at the [Colloquium of Sylvanès 2012](http://www.sylvanes.com/centre_culturel_colloques.html)
7. *"Although the piezoelectric potentials in vitro can be above* ***ten mV*** *if the fibres share a homogenous direction and polarity, they are drastically reduced for a set of bundles which have opposite polarities. In vitro research with respect to collagen I piezoelectricity has consistently found the fibrils to be randomly oriented; i.e. one direction mixed with the other. This is somewhat in contradiction with the macroscopic in vivo measurements: if the fibrils are oppositely oriented (50/50), then the piezoelectric effect should be cancelled, but this is not the case. This suggests that globally, the number of fibrils oriented in one direction is greater than the number of fibrils oriented in opposite direction”* (Harnagea, pers. com., 2012).
8. # [Denning](http://pubs.acs.org/author/Denning%2C+Denise) D. et al., The piezoelectric tensor of collagen fibrils determined at the nanoscale, *ACS Biomater Sci Eng*. Publication Web, DOI: 10.1021/acsbiomaterials.7b00183 ( 2017).
9. According to the "Wolff’s Law", structure and shape of bone permanently adapt to the loading conditions. Wolff regarded this transformation law, which had also been influenced by Rudolf Virchow (1821-1902), as a "brick to complete the building of Charles Darwin" (1809-1892).
10. R.D. Sinelnikov, Atlas of the anatomy of man, Study on the bones, joints, ligaments and muscles (updated & augmented, 1963, I: 232). Р. Д. CИHEЛБHИKOB, ATЛАС AHATOMИИ ЦELOBEKA (CCCP 1963) I: 232; [R.D. Sinelnikov, Atlas Anatomie Celoveka, v trekh tomakh]
11. [Denning](http://pubs.acs.org/author/Denning%2C+Denise) D. et al., The piezoelectric tensor of collagen fibrils determined at the nanoscale, *ACS Biomater. Sci. Eng;* DOI:10.1021/acsbiomaterials.7b00183 ( 2017).
12. Denning Denise, et al., Piezoelectric Tensor of collagen fibrils Determined at the Nanoscale, ACS Biomaterials Science & Engineering, 3 : 929-935 (2017).
13. Aron M., Floyd D., and Bance M., Voluntary eardrum movement: a marker for Tensor Tympani contraction ? *Otol Neurotol* **36** 1-9  (2014).
14. Bance M. et al., Effects of Tensor Tympani muscle contraction on the middle ear and markers of a contracted muscle, *Laryngoscope* **123,** 1021–27 (2013).
15. Henson M.M., Madden V.J., RaskAndersen H., Henson Jr O.W.., Smooth muscle in the annulus fibrosus of the tympanic membrane in bats, rodents, insectivores, and humans, *Hear Res* **200,** 2937 (2005).
16. Puria S, Fay RR, ‎Popper AN,‎ Smooth muscle in the annulus fibrosus of the tympanic membrane in bats, rodents, insectivores, and humans (2013).
17. <https://upload.wikimedia.org/wikipedia/commons/thumb/3/32/Ground_loop.svg/640px-Ground_loop.svg.png>
18. Q is the electrical charge (unit = Coulomb), V is the voltage (unit = Volt) and C is the capacitance (unit = Farad)
19. <http://www.thinksrs.com/downloads/PDFs/ApplicationNotes/AboutLIAs.pdf>
20. Gavilan C., Sanjuàn J., Microphonic Potential picked up from the human tympanic membrane, *Ann Otol Rhinol Laryngol* **73,** 102-9 (1964).
21. Davis H., Derbyshire A.J., Lurie M.H. and Saul L.J., The electric Response of the Cochlea, *Am. J. of Ph*. **107**, 311-32 (1934).
22. Reyes S., Ding Da., Sun W., Salvi R., Effect of Inner and Outer Hair cell lesions on electrically evoked otoacoustic emissions, *Hear Res* **158,** 139-50 (2001).
23. Wever, E.G. and C.W. Bray, Auditory nerve impulses, *Science* **71,** 215 (1930).
24. Trautwein P., Hofstetter P., Wang J., Salvi R. and Nostrant A., Selective inner hair cell loss does not alter DPOAEs, *Hear. Res*. **96,** 71-82 (1996).
25. Gavilan C. and Sanjuàn J., Microphonic Potential picked up from the human tympanic membrane, *Ann Otol Rhinol Laryngol* **73,**102-9 (1964).
26. Carricondo F, GilLoyzaga P, SanjuànJuaristi J, PochBroto J., Cochlear microphonic potentials: a new recording technique. *Ann Otol Rhinol Laryngol* **110**, 56573 (2001).
27. Carricondo F, GilLoyzaga P, SanjuànJuaristi J, PochBroto J., Cochlear microphonic potentials: a new recording technique. *Ann Otol Rhinol Laryngol* **110**, 56573 (2001).
28. PochBroto J. et al., Cochlear microphonic audiometry: a new hearing test for objective diagnosis of deafness, *Otol Rhinol Laryngol* **129,** 74954 (2009).
29. Mammano F., personal discussion [[Colloque de Sylvanès 2012](http://www.sylvanes.com/centre_culturel_colloques.html) (18-20 Mai 2012)]
30. Carricondo F et al., (2001).
31. PochBroto J. et al. (2009).
32. Behari J. *Biophysical Bone Behaviour: Principles and Applications*, Wiley - Science (2009).
33. [Wada](http://asa.scitation.org/author/Wada%2C+Hiroshi) H., [Ando](http://asa.scitation.org/author/Ando%2C+Masayoshi) M., [Takeuchi](http://asa.scitation.org/author/Takeuchi%2C+Masataka) M., [Sugawara](http://asa.scitation.org/author/Sugawara%2C+Hironori) H., [Koike](http://asa.scitation.org/author/Koike%2C+Takuji) T., Vibration measurement of the tympanic membrane of guinea pig temporal bones using time-averaged speckle pattern interferometry, JASA **111,** 2189-99 (2002).
34. Okayasua T. Human ultrasonic hearing is induced by a direct ultrasonic stimulation of the cochlea, *Neurosci Lett* **539**, 71–6 (2013).
35. Humbel R.L. Auto-anticorps dans les maladies de l'oreille interne, 4° Colloque GEAI (2006)
36. ISO, Acoustics, Hearing protectors, Part 1: Subjective method for the measurement of sound attenuation (1990).
37. Margolis R.H., The Vanishing Air-Bone Gap - Audiology's Dirty Little Secret, *Audiol Online* (2008).
38. Stenfelt, S.& Reinfeldt, S., A model of the occlusion effect with bone-conducted stimulation. *International J Audiology* **46**, 595-608 (2007).
39. [Denning](http://pubs.acs.org/author/Denning%2C+Denise) D. et al., The piezoelectric tensor of collagen fibrils determined at the nanoscale, *ACS Biomater. Sci. Eng;* DOI:10.1021/acsbiomaterials.7b00183 ( 2017).
40. The longitudinal piezoelectric coefficient for individual fibrils at the nanoscale was found to be roughly an order of magnitude greater than that reported for macroscopic measurements of tendon, the low response of which stems probably from the presence of oppositely oriented fibrils, as confirmed here.
41. Nachtigall PE, Lemonds DW, and Roitblat HL, Psychoacoustic of Dolphin and Whale Hearing, p.336-9 in Au WWL, Popper AN, and Fay RR, Hearing by Whales and Dolphins, Springer NY (ISBN 0-387-94906-2) (2000).
42. Dolphin W.F. pp.322-325.
43. Bullock et al., Electrophysiological studies of central auditory mechanisms in cetaceans, *Zeitschrift Vergleichen Physiol* **59**, 117-56 (1968).
44. Fraser FC and Purves PE, A Discussion on the 'Ear' Under Water, *Proc R Soc Lond B Biol Sci*, **152,** 62-77, (1960).
45. Ridgway SH The Auditory Central Nervous System of Dolphins. In: Au WWL. Fay RR, Popper AN (eds) Hearing by Whales and Dolphins, *Springer Handbook of Auditory Research* **12**. Springer, New York, NY (2000).
    Ridgway SH The Auditory Central Nervous System of Dolphins. In: Au WWL, Fay RR, Popper AN (eds) Hearing by Whales and Dolphins, Springer Handbook of Auditory Research, vol.**12**, Springer, New York (2000).
46. Ridgway SH The Auditory Central Nervous System of Dolphins. In: Au WWL. Fay RR, Popper AN (eds) Hearing by Whales and Dolphins, *Springer Handbook of Auditory Research* **12**. Springer, New York, NY (2000).
47. Alberts B. et al., *Molecular biology of the cell,* 5°ed., Garland Science, NY, pp. 1474-6, ISBN 978-0-8153-4106-2 (2008).
48. Yamato, M., Ketten, D. R., Arruda, J., Cramer, S. and Moore, K. (2012), The Auditory Anatomy of the Minke Whale (Balaenoptera acutorostrata): A Potential Fatty Sound Reception Pathway in a Baleen Whale. Anat Rec, 295: 991–998. doi:10.1002/ar.22459
49. Cranford TW, Krysl P, Amundin M (2010) A New Acoustic Portal into the Odontocete Ear and Vibrational Analysis of the Tympanoperiotic Complex. PLoS ONE5(8): e11927. https://doi.org/10.1371/journal.pone.0011927
50. Cranford TW, Krysl P, Amundin M, A new acoustic portal into the odontocete ear and vibrational analysis of the tympanoperiotic complex, *PLoS ONE* **5**:e11927 (2010).
51. Yamato MK et al., *Anat Rec*, **295,** 991-8 (2012).
52. Alberts B. et al., *Molecular biology of the cell,* 5°ed., Garland Science, NY, pp. 1474-6, ISBN 978-0-8153-4106-2 (2008).
53. Graham, A. , Okabe, M. and Quinlan, R., The role of the endoderm in the development and evolution of the pharyngeal arches. J Anat, 207: 479-487 (2005).
54. Hill, M.A. Embryology *Hearing - Middle Ear Development*. Cf. <https://embryology.med.unsw.edu.au/embryology/index.php/Hearing_-_Middle_Ear_Development> (2018).
55. Gelman SR Wood S., Spellacy WN, Abrams RM, [Fetal movements in response to sound stimulation](https://www.ajog.org/article/0002-9378(82)90097-7/abstract), Am J Obstet Gynecol 143: 484-485 (1982).
56. Dunn, K., Reissland, N., Reid, V.M.,The Functional Fetal Brain: A Systematic Preview of Methodological Factors in Reporting Fetal Visual and Auditory Capacity, Developmental Cognitive Neuroscience, <http://dx.doi.org/10.1016/j.dcn.2015.04.002> (2015)
57. Boistel, R., et al., How minute sooglossid frogs hear without a middle ear. PNAS, 110(38): 15360–15364. <https://doi.org/10.1073/PNAS.1302218110>. (2013).
58. Tillaux P.J., *Traité d’Anatomie topographique avec applications à la chirurgie*, Paris Asselin et Houzeau, 4°ed. p.116 (1903).
59. Testut L. et Jacob O., Traité d'Anatomie Topographique, T.1 Fig. 220, p.302, Doin Paris (1905)
60. Petit C., Cours au Collège de France, 18 et 25 mars, ler et 8 avril 2004, Mécanismes d'analyse des fréquences sonores.
61. Santos-Sacchi J., On the frequency limit and phase of outer hair cell motility: effects of the membrane filter, J of Neurosc, 12, 1906-16 (1992).
62. Gale J. E., Ashmore J. F, An intrinsic frequency limit to the cochlear amplifier, *Nature* **389**, 63–6 (1997).
63. Zha D et al., In Vivo Outer Hair Cell Length Changes Expose the Active Process in the Cochlea. PLoS ONE 7(4): e32757 (2012).
64. Dallos P, Evans BN, High-frequency motility of outer hair cells and the cochlear amplifier, *Science* **267,** 2006-9 (1995).
65. Fridberger A, Organ of Corti Potentials and the Motion of the Basilar Membrane, *J. Neurosc*, **24,** 10057-63 (2004).
66. Spector A.A., Popel A.S., Eatock RA, Brownell WE, Mechanosensitive channels in the lateral wall can enhance the cochlear outer hair cell frequency response. *Ann. Biomed. Eng*. **33**, 991–1002 (2005).
67. Kaneko T. , Harasztosi C. , Mack AF , Gummer AW., Membrane traffic in outer hair cells of the adult mammalian cochlea, Eur. J. Neurosci **23**, 2712–22, (2006).
68. Dallos P, Evans BN, High-frequency motility of outer hair cells and the cochlear amplifier, *Science* **267**, 2006-9 (1995).
69. Ashmore JF., Cochlear outer hair cell motility, *Physiol. Rev* **88**, 173–210 (2008).
70. Mistrik, P., Mullaley, C., Mammano, F., Ashmore, J., Three-dimensional current flow in a large-scale model of the cochlea and the mechanisms of amplification of sound, *J. Roy. Soc*. Interface **6**, 279–91 (2009).
71. Mistrík P., and Ashmore JF., Reduced Electromotility of Outer Hair Cells Associated with Connexin-Related Forms of Deafness: An In silico Study of a Cochlear Network Mechanism, JARO **11**, 559–71 (2010).
72. Ashmore J et al., The remarkable cochlear amplifier, *Hear Res* **266**, 1–17 (2010).
73. Zhao HB and Yu N, Distinct and gradient distributions of connexin26 and connexin30 in the cochlear sensory epithelium of guinea pigs, *J Comp Neurol* **499,** 506-18 (2006).
74. Laird DW, The gap junction proteom and its relationship to disease, [*Trends Cell Biol*](javascript:AL_get(this,%20'jour',%20'Trends%20Cell%20Biol.');)*,* **20**,92-101, (2010).
75. Dallos P et al. (1995).
76. Fridberger A., Loud sound-induced changes in cochlear mechanics*. J Neurophysiol* **88**, 2341- 8 (2002).
77. Cody AR, Russell IJ, The responses of hair cells in the basal turn of the guinea-pig cochlea to tones. *J Physiol* **383**, 551–69 (1987).
78. Lagostena L, Cicuttin A, Inda J, Kachar B, Mammano F, Frequency Dependence of Electrical Coupling in Deiters' Cells of the Guinea Pig Cochlea, *Cell Communication & Adhesion,* **8**, 393 –9 (2001).
79. cf. Cody and Russell p.554 : “ in response to tones of 10 kHz or greater, the c.m. recorded extracellularly in the organ of Corti and intracellularly in the supporting cells is 2.20 times greater than the outer hair cell receptor potentials. […] Furthermore the c.m. recorded from supporting cells is unchanged by current injection while the receptor potentials of outer hair cells are increased by hyperpolarizing currents and decreased by depolarizing current injection”.
80. Harroun TA., Katsaras J, and Wassall SR, Cholesterol is Found to Reside in the Center of a Polyunsaturated Lipid Membrane, Biochemistry, **47**, 7090–6, (2008).
81. Marrink S. J., de Vries A H, Harroun T A, Katsaras J, and Wassall SR Cholesterol Shows Preference for the Interior of Polyunsaturated Lipid Membranes. *J Am Chem Soc,* **130**, 10–11 (2008).
82. Nguyen TV, Brownell WE, Contribution of membrane cholesterol to outer hair cell lateral wall stiffness, *Otolaryngol Head Neck Surg* **119**,14–20 (1998).
83. These domains correspond to segregated membrane proteins, Prestin in the lateral wall (Ashmore 2008) and KCNQ4 channels at the base (Mustapha et al. 2009), suggesting a unique relationship between these cholesterol domains and membrane function.
84. Harroun TA, Katsaras J, and Wassall SR, Cholesterol is Found to Reside in the Center of a Polyunsaturated Lipid Membrane, *Biochemistry*, **47,**  7090–6 (2008)
85. Wanders R.J.A., Peroxisomes, lipid metabolism, and peroxisomal disorders, *Mol Genet Metab* **83,** 16-27 (2004).
86. The capacitance of a cell membrane varies according to the voltage applied to the membrane of a neighbour cell ("ephaptic effect" ). If we apply this model to the DOHC complex, we can suggest that the capacitance of the cuticular bilayer varies in a manner controlled by the voltage of the contiguous phalangeal apexes. In this way, the piezoelectric information, carried from the tympanum to the four phalanxes of the DCs, should cause its capacitance to vary in a synchronous manner.
87. Arvanitaki A., Effects evoked in an axon by the activity of a contiguous one, *J Neurophysiol* **5**, 89-108 (1942).
88. [Goldwyn JH](https://www.ncbi.nlm.nih.gov/pubmed/?term=Goldwyn%20JH%5BAuthor%5D&cauthor=true&cauthor_uid=26823512), [Rinzel J](https://www.ncbi.nlm.nih.gov/pubmed/?term=Rinzel%20J%5BAuthor%5D&cauthor=true&cauthor_uid=26823512), Neuronal coupling by endogenous electric fields: cable theory and applications to coincidence detector neurons in the auditory brain stem, [*J Neurophysiol*,](https://www.ncbi.nlm.nih.gov/pubmed/26823512) **115**, 2033-51 (2016).
89. Zhang M et al.., Propagation of epileptiform activity can be independent of synaptic transmission, gap junctions, or diffusion and is consistent with electrical field transmission. *J Neurosci* **34**, 1409–1419, 2014.
90. LilienfeldJ.E.,https://www.google.com/search?tbo=p&tbm=pts&hl=en&q=ininventor:%22Lilienfeld+Julius+Edgar%22 Method and apparatus for controlling electric currents, US 1745175A patent (1930).
91. Van Itallie Christina M., Fanning Alan S., Bridges Arlene, and Anderson James M., ZO-1 Stabilizes the Tight Junction Solute Barrier through Coupling to the Perijunctional Cytoskeleton, Molecular Biology of the Cell, Vol. 20, 3930–3940, September 1, 2009.
92. Goldwyn JH and Rinzel J, Neuronal coupling by endogenous electric fields, in <http://adsabs.harvard.edu/abs/> arXiv150801741G (2015). [The authors provide a cable theory framework to study how a bundle of model neurons generates V_e_ and how this V_e_ feeds back and influences membrane potential (V_m_)].
93. Buzsáki G, Anastassiou CA, Koch C. The origin of extracellular fields and currents. Nat Rev Neurosci 13: 407–420, **2012**.
94. Szarama Katherine B., Núria Gavara, Petralia Ronald S., Kelley Matthew W. and Chadwick Richard S., Cytoskeletal changes in actin and microtubules underlie the developing surface mechanical properties of sensory and supporting cells in the mouse cochlea, Development 139, 2187-2197 (2012) ; doi:10.1242/dev.073734
95. Sataric, M.V., et al., Role of nonlinear localized Ca2+ pulses along microtubules in tuning the mechano-sensitivity of hair cells, *Prog Biophys Mol Biol* **119**,162–74 (2015).
96. See: Vater M., Lenoir M., and Pujol R., Developpment of the organ of Corti in Horseshoe Bats: Scanning and Transmission Electron Microscopy, *J Comp Neurol*, **377**, 520-534 (1997).
97. Mahendrasingam S, Beurg M, Fettiplace R, Hackney CM, [The ultrastructural distribution of prestin in outer hair cells: A post-embedding immunogold investigation of low and high frequency regions of the rat cochlea](https://www.ncbi.nlm.nih.gov/pmc/articles/PMC2925464). *Eur J Neurosci,* **31,** 1595–605 (2010).
98. Priel A, Tuszynski JA, A nonlinear cable-like model of amplified ionic wave propagation along microtubules,. *Eur Phys Lett,* **83**, 68004 (2008).
99. Zha D et al., In Vivo Outer Hair Cell Length Changes Expose the Active Process in the Cochlea. PLoS ONE 7(4): e32757 (2012).
100. Ricci, AJ, Crawford AC and Fettiplace R, Tonotopic variation in the conductance of the hair cell mechanotransducer channel, *Neuron* **40**, 983–90 (2003).
101. Toupet M., Les sons forts peuvent donner le vertige – peut-on entendre par le vestibule ?, *Inst Méd*, **4,** 21-4, (1981).
102. Sheykholeslami, K, Kaga K, The otolithic organ as a receptor of vestibular hearing revealed by vestibular-evoked myogenic potentials in patients with inner ear anomalies, *Hear Res,* **165**, 62–7 (2002).
103. Ashmore J, Cochlear outer hair cell motility, ‎*Physiol Rev* **88**, 173-210 (2008).
104. It is possible that this "suppressive" effect would be produced by a phase inversion during the amplification process of these infra-sound frequencies, thus producing an attenuation of these stimuli either when they are unuseful, or would become disruptive.
105. Rogers PH, Hawkins AD, Popper AN, Fay RR, Gray MD, Parvulescu Revisited, Small Tank Acoustics for Bioacousticians In: Popper A, Hawkins A (eds) The Effects of Noise on Aquatic Life, *Adv Exp Med Biol,* **875**, Springer, NY (2016).
106. [Steinmetz](https://www.google.fr/search?tbo=p&tbm=bks&q=inauthor:%22Charles+Proteus+Steinmetz%22) CP, [Electric engineering](https://www.google.fr/search?tbo=p&tbm=bks&q=subject:%22Electric+engineering%22&source=gbs_ge_summary_r&cad=0), Theory and Calculation of Transient Electric Phenomena and Oscillations, **8**, McGraw-Hill Book (1920).
107. Wenxuan He, David Kemp and Tianying Ren, Timing of the reticular lamina and basilar membrane vibration in living gerbil cochleae, eLife 2018; 7: e37625.
108. Fridberger A., et al., Organ of Corti Potentials and the Motion of the Basilar Membrane, *J Neurosci* **24**, 10057–63 (2004).
109. Ren T, He W, Scott M and Nuttall AL, Group Delay of Acoustic Emissions in the Ear, *JN Physiol* **96** , 2785-91 (2006).
110. Grosh K, Zheng J, Zou Y, de Boer E, and Nuttall AL., High-frequency electromotile responses in the cochlea, JASA, **115**, 2178-84 (2004).
111. Reyes S, Ding D, Sun W, Salvi R, Effect of Inner and Outer Hair cell lesions on electrically evoked otoacoustic emissions, *Hear Res*, **158,** 139-50 (2001).
112. Reyes S. et al. (2001).
113. [He](http://www.ncbi.nlm.nih.gov/pubmed/?term=He%20W%5Bauth%5D) W.and [Ren](http://www.ncbi.nlm.nih.gov/pubmed/?term=Ren%20T%5Bauth%5D)a T., Basilar membrane vibration is not involved in the reverse propagation of otoacoustic emissions, *Sci Rep,* **3,** 1874 (2013).
114. For 4 Individuals (9 Series), their exact age was not specified and will be recovered
115. Quix F. H.. Mil. Geneesk Tydschr (1903).
116. Zwaardemaker H., Sur la sensibilité de l'oreille aux différentes hauteurs des sons, [*L'année psychologique*](http://www.persee.fr/collection/psy) **10,** 161-78 (1903).
117. Vasilescu G., Electronic Noise and interfering signals, principles and applications, *Signals and communication technology*, Springer Verlag, ISBN 3-540-40741-3 (2005).
118. http://www.cnrtl.fr/definition/vicariance
119. Seldran, F., A model-based analysis of the “combined-stimulation advantage”, *Hear. Res.* **282**, 252-64 (2011).
