## Supplementary Information for "Overt and covert paths for sound in the auditory system of mammals"

| | |
|---|---|
| (a) | (b) |

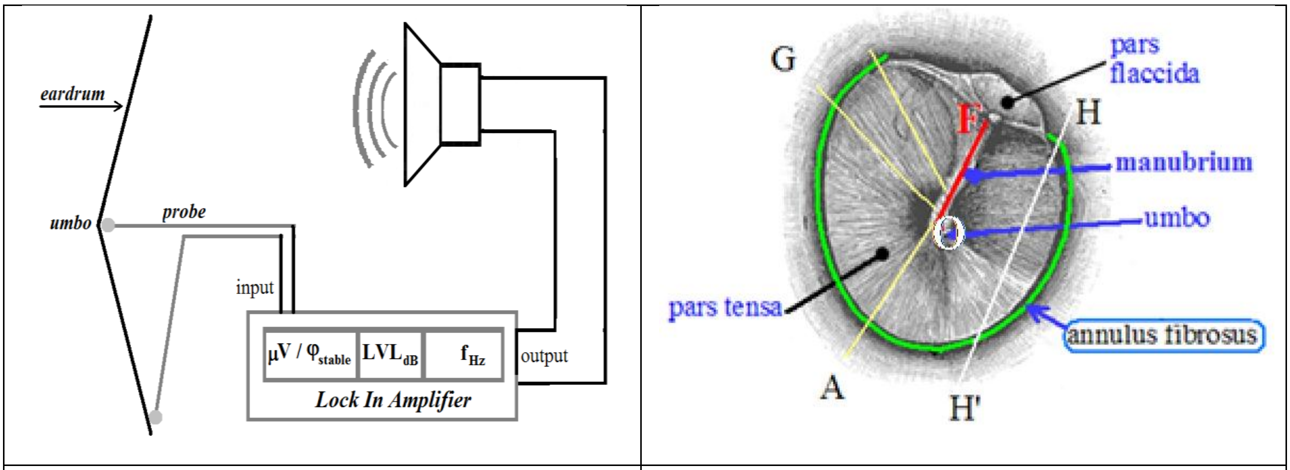

**a** **b**

**Figure S01**

---

[1] Gavilan C., Sanjuàn J., Microphonic Potential picked up from the human tympanic membrane. *Ann Otol Rhinol Laryngol.* **73,** 102-109 (1964).

The closer together the electrodes are placed, the smaller the area recorded will be (i.e. local recordings). This is one advantage of the differential recording technique[111]. This "differential technique" is necessary for determining the source of a potentials[112].

#### Si 01F  Measurements of piezoelectric response on knees (or other tendons)

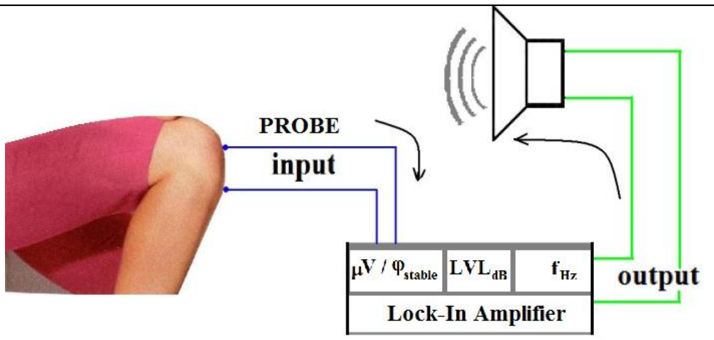

Figure S02
Diagram of the design used to measure the piezoelectric potential from the knee tendon.
□V indicates the isochronous voltage obtained when the phase j is stabilized, in response to the sound wave of frequency f (Hz) with a LVL (dBSP) amplitude.

#### Measurement  Electrodes for tendons

For measurements on the **mastoid** area, the patellar ligament or other tendons, we used « single use » *pregelled ECG-electrodes (Ref. “Comepa” 3.02.0200); they were connected to the* LockIn Amplifier by two *medical leadwires* AIP018 with *Touch proof connectors* DIN42802. (Table S09)

##### Effect of the length of the collagen fibres

To evaluate the effect of the length of the collagen fibre bundles on their piezoelectric response, we performed some experiments on the most accessible bundles (patellar ligament). This allowed us to understand in particular why the response of these bundles was much larger than those of the eardrum.

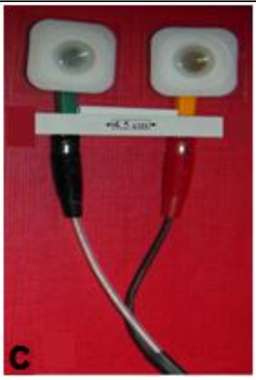

**(c)** Probe used for measurement on the **knees**
As the amplitude of the voltage generated (in µV) increases with the distance (in mm) between electrodes; we, thus, ensured a fixed distance (45 mm) between the centre of the two COMEPA, 3020200 CA electrodes.
**Figure S03**

| d (mm) | V (µV) |
|---|---|
| 40 | 12.1 |
| 45 | 15.1 |
| 50 | 18.0 |
| 55 | 22.8 |
| 60 | 28.2 |
| 65 | 30.0 |

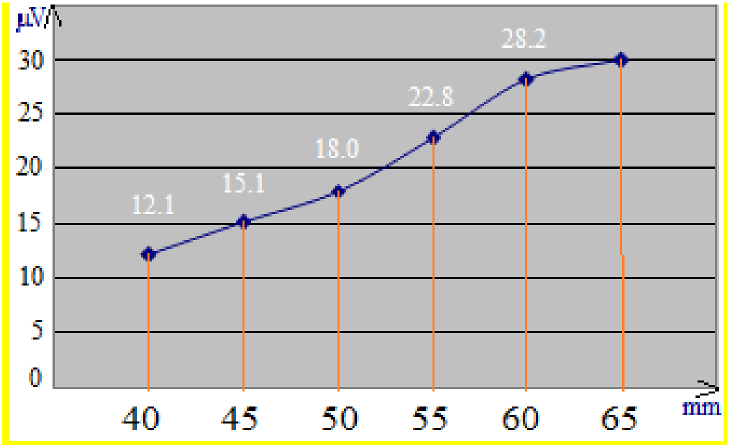

**Figure S04**

**Measurements on knee** : Acoustical frequency of the stimulus was 4000 Hz
x axis (d): distance (mm); y-axis : voltage (µV).
Result : **V (µV) = 0.76 d (mm) -19 mm ; $R^2$ = 0.98**

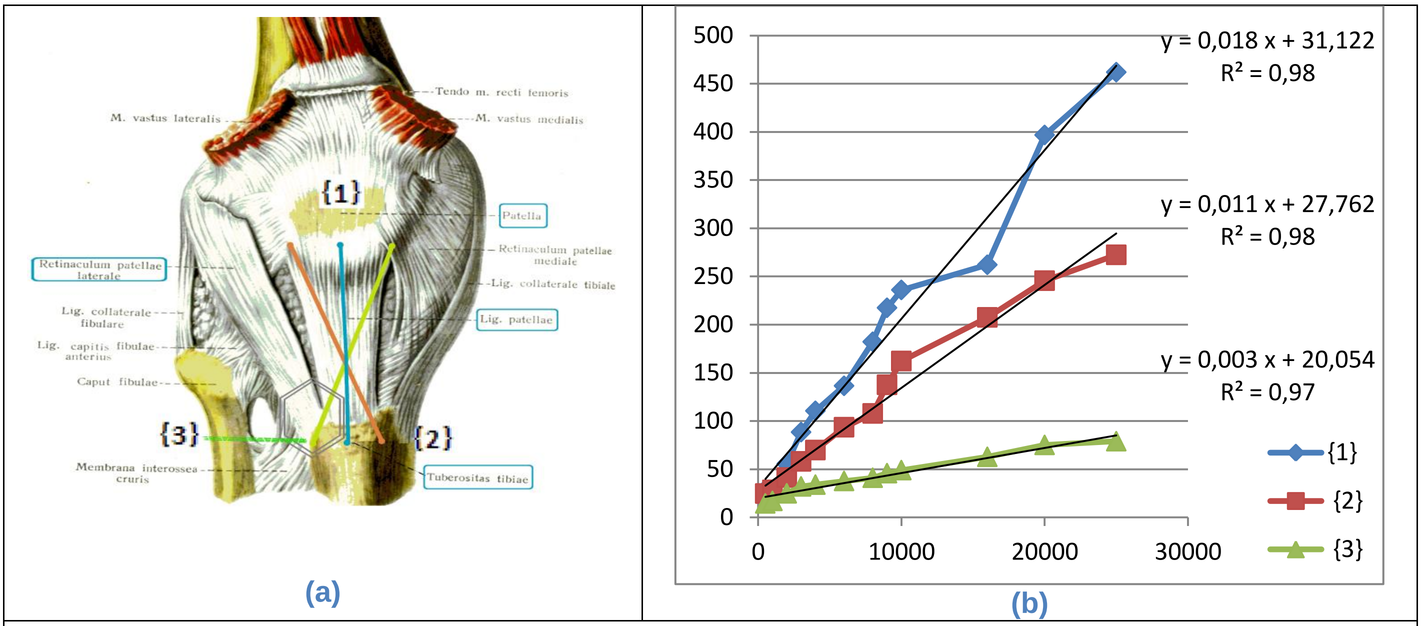

Figure S05a and S05b

Experimental mimicry of offset on fibres of a patellar ligament: We used voluntarily three configurations for comparisons between a good precision {1} versus bad ones {2 & 3} (The knee anatomy figure is excerpted from Sinelnikov[S115]).

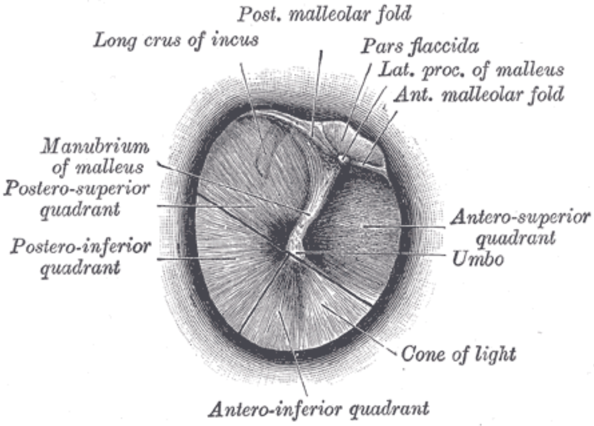

Figure S06
internal aspect of tympanic membrane

The radial fibres of collagen grow from the umbo to the periphery .
The ENT experimenter tries to put both of the probe electrodes at the ends of one single fibre. Yet such an utmost precision is unreachable because every fibres are extremely thin and not separately identifiable. So it happens an unwished offset on fibres of the tympanum.

Furthermore we have connected the different outlets of the system (lock-in, speaker) on the same power strip and the same ground socket.

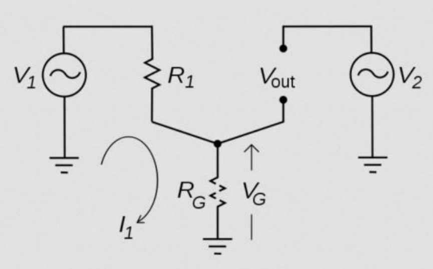

Figure S07
"ground artefact" excerpted from en-WP[S121] (R resistance, G ground, V Voltage, I intensity; in this schema there are three links to the ground).

##### Si 04B Artefact resulting from a "hand effect"

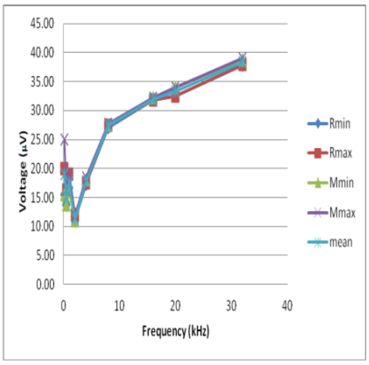

Figure S08
Mean Voltages (µV) measured for several frequencies
(R means Results without Hand Effect; M means results with Hand effect). In all configurations, when there is a change, the "hand effect" causes an increase of the voltage <1.5 µV for voltages around 25 µV , ie 6% (Fig. S08 above).

Additional connection of a conducive bracelet around the wrist of the experimenter engenders a higher "hand effect" unless the metallic case is also connected to the ground. The most practical solution, in order to avoid any hand effect, is to remove the hand of more than 20 cm from the casing.

##### Si 04C Artefact depending on the contact between electrodes and the skin-target

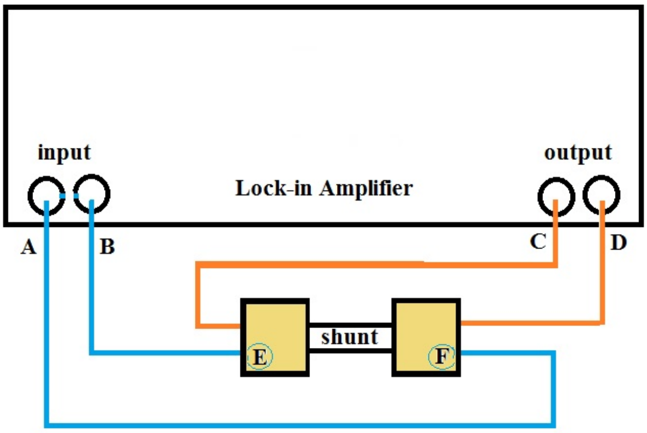

Figure S09

In fig. S04 (above) we replaced the skin of the subject by a simple copper shunt, with a resistance Rs = 0.2 Ω (AOIP 02.2351); This shunt is purely resistive up to a few tens of kHz. And the current from the output of the Lock-in (C and D) is introduced (by red circuit: coaxial cable and alligator clips) into the shunt.
The electrodes (E and F), normally applied to the skin, here are connected by the blue wires to the Lock-in Amplifier; they have different forms and surfaces (round or square COMEPA, or Alligator Clips).
Since our measures on tendons present values in the range from 10 to 200 µV, we adjust the settings so that the potential difference between E and F is close to 200 µV. As the resistance of the shunt is 0.2 Ω, we must send a current of 200 µV / 0.2 Ω, ie 1mA flowing through this shunt. To arrive at this value of 1 mA, if we take the device as a whole, including the two resistors in series, the output of the Lock-in (Ro = 50 Ω), and the resistance of the shunt (Rs = 0.2 Ω), we have to adjust the output voltage of the Lock-in to about 50 mV since {1 mA * (Ro + Rs)} = 1 * 50.2 = 50.2 mV. The resulting voltage between E and F (in µV) to the terminals of the shunt is measured at the input of the Lock-in (be A and B; fig. S04 above) with the blue cables. By varying the frequency (kHz), we obtain the results presented in Table S01 and Fig. S05.

| | **two square electrodes** | **one round and one square electrode** | **two round electrodes** | **two alligator clips** |
|---|---|---|---|---|
| **f (kHz)** | (µV) | (µV) | (µV) | (µV) |
| **0.2** | 0.135 | 0.142 | 0.164 | 0.196 |
| **0.8** | 0.136 | 0.138 | 0.166 | 0.198 |
| **2** | 0.138 | 0.145 | 0.165 | 0.196 |
| **4** | 0.142 | 0.146 | 0.160 | 0.196 |
| **8** | 0.134 | 0.135 | 0.143 | 0.202 |
| **20** | 0.185 | 0.188 | 0.198 | 0.219 |

Ro=50 Ω et Rs=0.2 Ω
**Table S08**

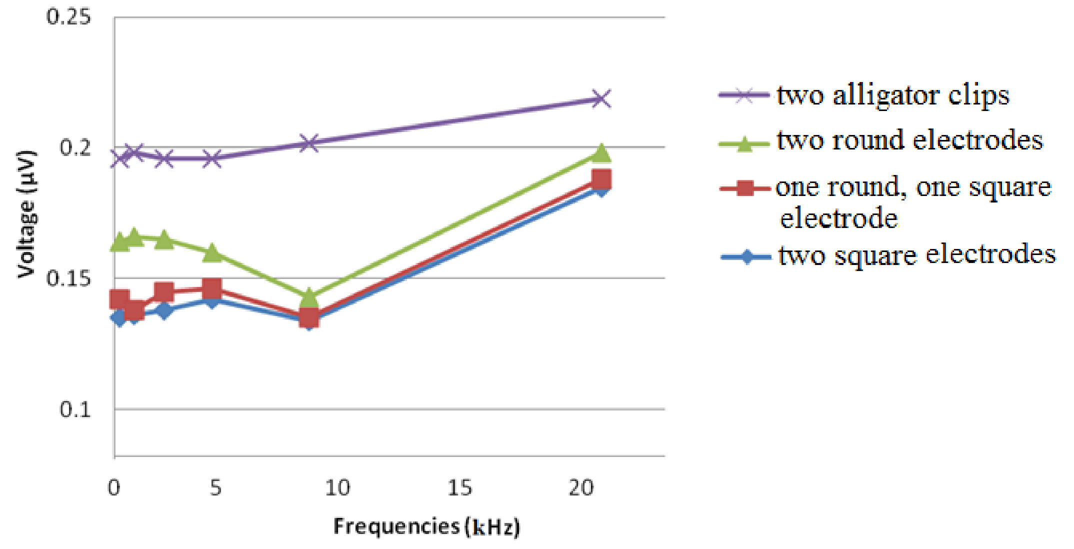

Figure S10

##### Si 04D Is there artefact depending on microphonic of measurement cables ?

Mechanical noise can be translated into electrical noise by microphonic effects. Physical changes in the cables (due to vibrations for example) can result in electrical noise over the entire frequency range of the lock-in. For example, consider a coaxial cable connecting a detector to a lock-in. The capacitance of a coaxial cable is a function of its geometry. Mechanical vibrations in the cable translate into a capacitance that varies in time typically at the vibration frequency. Since the cable is governed by CV= Q[5] taking the derivative yields: C(dV/dt) + V(dC/dt) = dQ/dt = i. Mechanical vibrations in the cable which cause a dC/dt will give rise to a current in the cable. This current may affect the measured signal [S122]. We conducted an evaluation of this effect in the context of our measurements: We set the power of the loudspeaker by the lock-in with an increased voltage (5V against 2.5V for previous experiments) and with the speaker very close to the coaxial cable that we use. Variations in capacitance of the coaxial cables that we used, are very low given the broadcast sound power. We observe no voltage cable, short circuit, or open on the range 0.1 - 40 kHz.

- Bone conduction allows to hear ultrasounds.

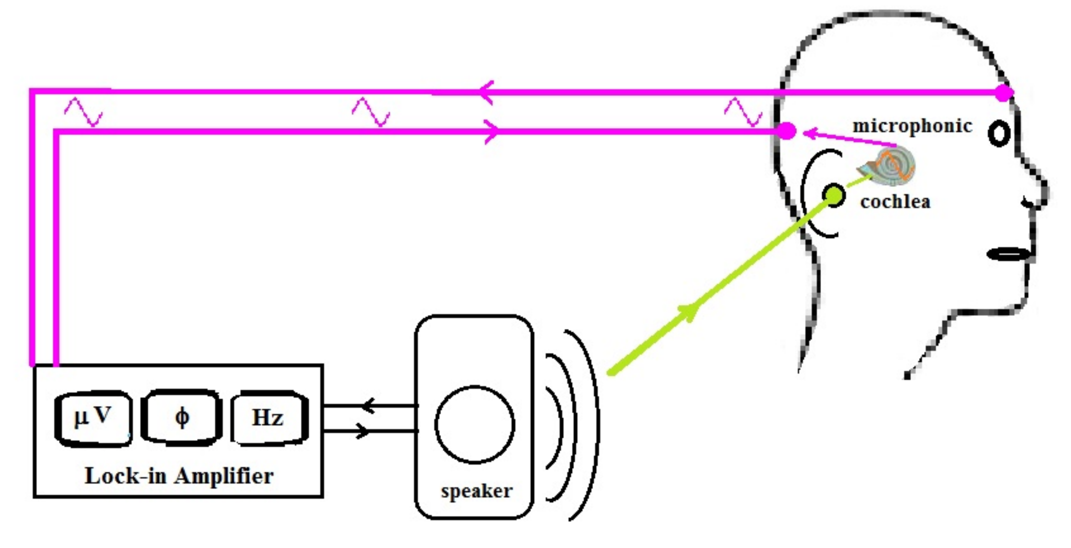

Figure S11
Sometimes accepted theory of the generation of the microphonic near the mastoid

The acoustic vibrations of the eardrum are transduced, by the cochlear OHCs, into a synchronous electric potential, known under the name of microphonics[S133]. Furthermore, such a synchronous electric potential can be recorded at the level of the skin behind the ear (with the other electrode being placed, for example, on the forehead). It is postulated that the cochlear generator alone would be responsible for that recorded potential. In other words, the recorded potential would result only from the scattering of the "cochlear microphonic" and for no other reason.

#### Si 05C The synchronous potentials, tympanic or mastoidian, are not uniquely dependent on the cochlea

In order to test whether pure-frequency sound stimulation produced a piezoelectric response, we measured the tympanum of a pig a few hours after its death, and after removal of the cochlea performed without damaging the anatomy of the eardrum.

**Piezoelectricity measurements on the pig tympanum at the Puylaurens abattoirs (1 07 2018).**
We carried out measurements on pig eardrums shortly after their slaughter, in the Puylaurens Archeology Club premises (room temperature 24 ° C).

**Acoustic measurements**
Sound source: Yamaha DBR12 loudspeaker, powered by the LockIn Stanford SR 830 (frequency and amplitude adjustment). The sound intensity measurement was carried out using a Bruel & Kjaer sound level meter, Type 2231 (measurements were made by Bernard Bibé, INRA). For the tests, we opted for a relatively moderate acoustic amplitude of about 75 dB SPL archeology of Puylaurens (room temperature 24 ° C).

**Measurements of electrical responses:**
Lock in Standford SR 830 (measuring frequencies 150 Hz-30kHz, 300ms time constant).
The probes consist of a coaxial cable that is connected to the input of the LockIn by a BNC plug.
This cable has two components: a fiber core consisting of a single fiber and a peripheral shield made of very fine braided metal fibers. The end of these two components is completed by a small metal ball.

**Measured samples:**

1. **Tympan 1** taken from a head of the day, not scalded.

2. **Tympan 2** taken from a scalded head

3. **Tympan 3** taken from a head of the day, not scalded but damaged during preparation.

4. **Bone conduction**: Bone taken from the external auditory canal, this bone is hemicylindrical (semicircular section). It consists of a wall of X mm thick, and a "channel" of Y mm width; this wall and this channel are oriented like generatrices of the bone cylinder.

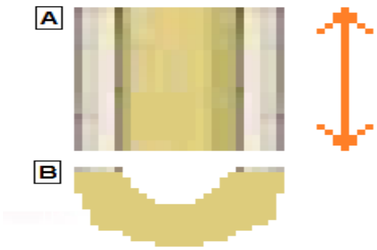

A ) Superior view of the osseous canal.
On the right: double arrow symbolizing the potential difference between the ends of the bone channel, when it is stimulated by a stable frequency acoustic wave.
B ) Sectional view of the 1/2 bony channel

6 measurements were made:

1) Tympanum 1: loudness 70dB
2) Tympanum 2: loudness 74dB
3) Tympanum 2 after removal of the cochlea: loudness 74dB
4) Tympanum 3: loudness 74dB

Hemisection of the bone canal:
5) loudness 70dB
6) loudness 90dB

We have calculated (Excel) the polynomial-type trend curve for each of the measurements made

**Measurement 1**: Tympanum 1 taken from a head of the day, not scalded: stimulus 70dB SPL

| Frequency (Hz) | Stimulus (dB SPL) | Electrical response (μV) |
|---|---|---|
| 200 | 70 | 34 |
| 500 | 70 | 9 |
| 1000 | 70 | 10 |
| 2000 | 70 | 4 |
| 4000 | 70 | 0 |
| 8000 | 70 | 3 |
| 16000 | 70 | 1,5 |

Tympanum 1; y(μV)= 2E-07 $x^2$ - 0,005 x + 18,59; $R^2$ = 0,47
ordinates: voltages (μV) ; abscissa: frequencies (Hz)

**Measurements 2**: Tympan 1 (Stimulus amplitude 70dB SPL)

| Frequency (Hz) | Piezoelectric signal (μV) |
|---|---|
| 500 | 0,2 |
| 1000 | 0,5 |
| 2000 | 0,2 |
| 4000 | 0,06 |
| 8000 | 0,08 |
| 16000 | 0,3 |
| 20000 | 0,27 |
| Although the signal was very weak, the measurements were stable during the measurement and the repetition of the measurement gave an identical result. | |

y = 2E-09 $x^2$ - 5E-05 x + 0,3214; $R^2$ = 0,30
ordinates: voltages (μV) ; abscissa: frequencies (Hz)

**Measurements 3**: Tympan 2 taken from a head of the day, not scalded, after ablation of the cochlea, sound intensity 74dB SPL.

| Frequency (Hz) | Piezoelectric signal (□V) |
|---|---|
| 250 | 4 |
| 500 | 0,25 |
| 1000 | 0,15 |
| 2000 | 0,18 |
| 4000 | 0,07 |
| 8000 | 0,11 |
| 16000 | 0,2 |

$$y = 2E\text{-}08\ x^2 + 1{,}62\ ;\ R^2 = 0{,}26$$

ordinates: voltages (μV) ; abscissa: frequencies (Hz)

**Measurements 4**: Tympanum 3 (stimulus amplitude 74dB SPL)

| Frequency (Hz) | Signal piezo (□V) |
|---|---|
| 250 | 20 |
| 500 | 6 |
| 1000 | 3 |
| 2000 | 2 |
| 4000 | 0,4 |
| 8000 | 0,8 |
| 16000 | 1,4 |

$$y = 1E\text{-}07x^2 - 0{,}003\ x + 10{,}51;\ R^2 = 0{,}44$$

ordinates: voltages (μV) ; abscissa: frequencies (Hz)

**Measurements 5**: Piece of External Bone Canal (70dB)

| Frequency (Hz) | piezoelectric signal (□V) |
|---|---|
| 150 | 0,87 |
| 500 | 0,5 |
| 1000 | 0,75 |
| 2000 | 1 |
| 4000 | 0,3 |
| 8000 | 1,1 |
| 16000 | 2,7 |

$$y = 1E\text{-}08x^2 - 8E\text{-}05x + 0{,}7737;\ R^2 = 0{,}91$$

ordinates: voltages (μV) ; abscissa: frequencies (Hz)

**Measurements 6**: Piece of External Bone Canal (90dB)

| Frequency (Hz) | piezoelectric signal (µV) |
|---|---|
| 150 | 25 |
| 500 | 25.74 |
| 1000 | 4 |
| 2000 | 6,5 |
| 4000 | 1,7 |
| 8000 | 6 |
| 16000 | 15,6 |

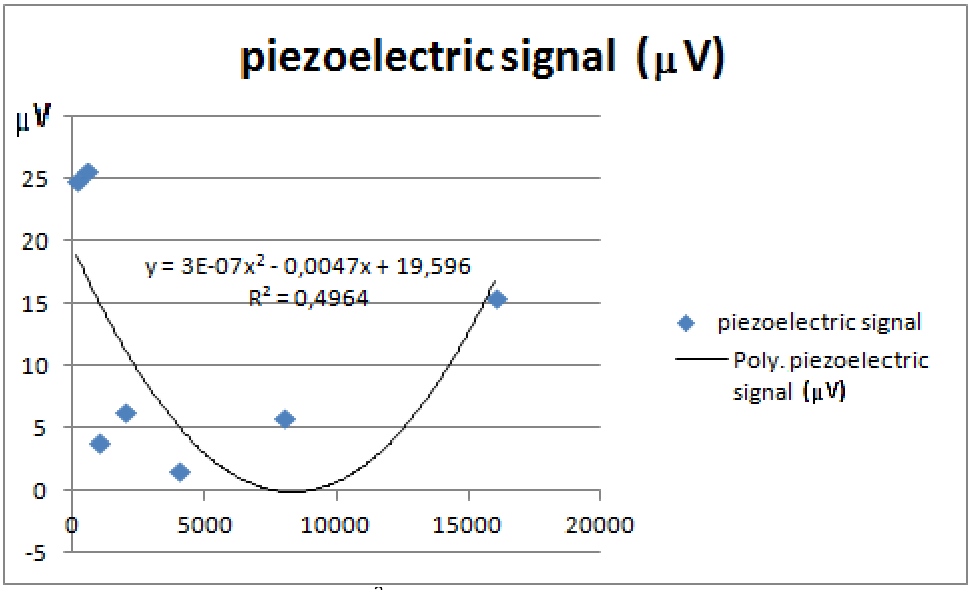

$y = 1E\text{-}06x^2 - 0{,}025x + 96{,}433$; $R^2 = 0{,}20$
ordinates: voltages (µV) ; abscissa: frequencies (Hz)

#### Si 05D Bone Conduction

##### Si 05Da Attempts to define a mastoid audiogram have been problematic

These findings and their restrictive interpretation led Poch-Broto and al. (2009) to try to create an audiometric test based on the recording of the electric potential of the mastoid in response to auditory stimulations. In this theory, the schema of transmission without earplug complies with fig. S07.

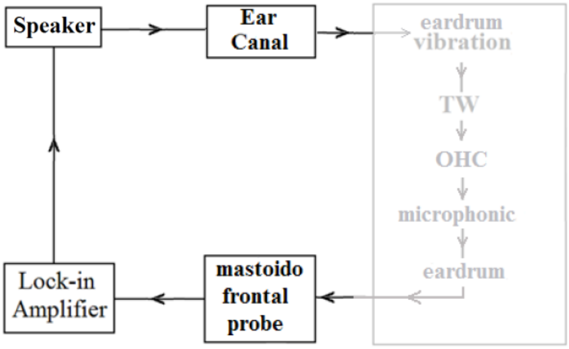

Figure S12
Capture of the mastoid "microphonic". Dimmed: Interpretation of Poch-Broto, attributing all the microphonic amplification to the TWs alone, via the OHC[S53].
But, if we compare the classic audiogram and this " mastoid " audiogram, we can see that very often (>10%) they differ by more than 10 dB HL mainly in the area of high frequencies, which is significant (fig. S08).

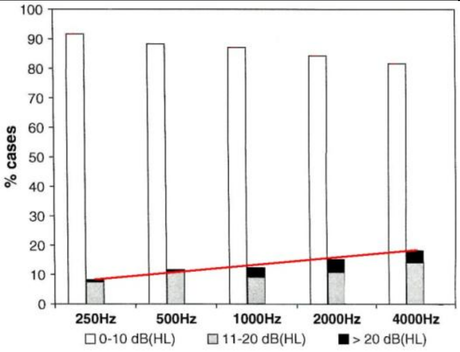

Figure S13
Modified chart based upon the data of Poch-Broto et al., 2009[S134]:
We grouped the black rectangles and the gray rectangles to represent all errors (> 10dB) combined. This clearly shows (red lines) that the error rate increases with frequency. The X axis displays the tested frequencies; the Y axis displays the % of cases according to whether the divergence between subjective audiometry and "microphonic" audiometry is less than (clear bars), or superior to (gray and black bars) 10 dB HL.

Since for all frequencies, in more than 10% of cases, the difference between subjective audiometry and measurement of the microphonic is greater than 10 dB, it can be concluded that the cochlear generator is not the sole origin of collected potentials (cf the line in red); this is particularly true for the high frequencies. We contacted Pablo Gil Loyzaga (corresponding author of the Group of Poch Broto and al.) and asked for their raw data. Yet, he answered that the raw data was no more available (Pablo Gil Loyzaga, com. pers., 07/18/2012).

##### Si 05Db Bone conduction permits the hearing of ultrasounds

Bone collagen is particularly adapted for the transduction of ultra-sounds[S135]. Ultrasounds between 20 kHz and 33 kHz are heard when they are being administered by bone conduction[S136 - S137]. They act directly on the hair cells (ie their hearing is induced by the ultrasounds themselves). And furthermore, in the case of extremely high pitched sounds, toxic damage of the auditory system damages the air-conduction hearing and improves bone-conduction hearing.
It is well demonstrated that the deterioration of collagen II is causative of auto-immune deafness[S138].

#### Si 05E Experiments involving closing off of the external auditory canal (EAC)

The schemas, tables and curves below show the synchronous potentials, depending on whether a plug is not or is inserted into the auditory external conduit (earplug interposed between the sound source and the eardrum, but not between the sound source and the mastoid) : fig. S09, S10, S11, S12.

##### Si 05Ea Mastoid 'microphonic' with EAC closed off :

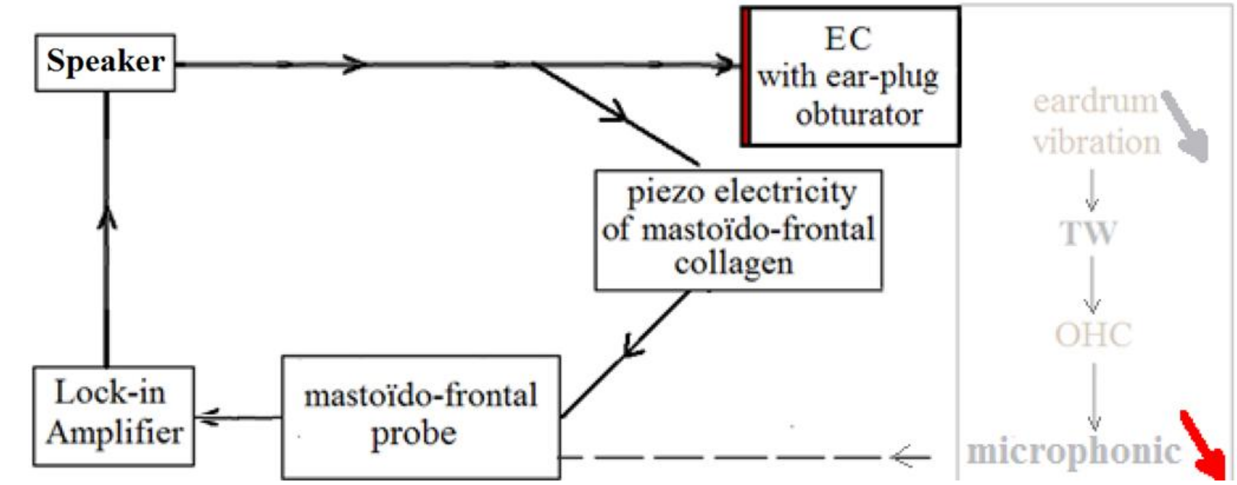

Figure S14
Capture of the mastoid 'microphonic' with the external auditory conduit closed off. (Dimmed and red arrows on the right side of the figure indicate the decrease of the mechanical vibration and of the electric potential as well).

Table S09 (**a**, **b**, **c**)
Synchronous mastoid voltage (µV), broadcasted by a loudspeaker Harman/Kardon, stimulated by a voltage of 2 Volts issuing from the lock-in amplifier;
Δ = Vs *with (closed)* minus Vs *without (open)*

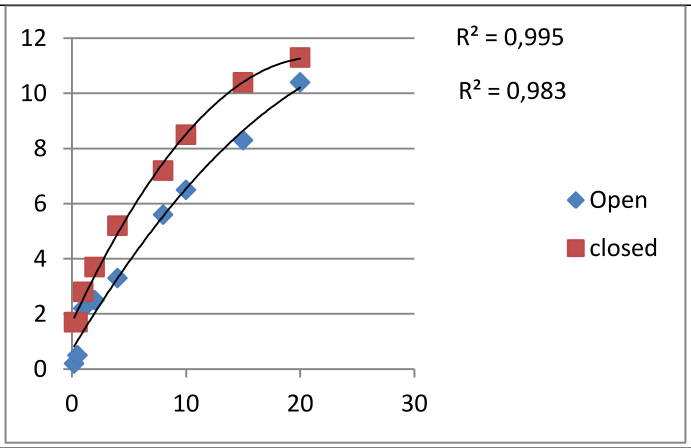

Figure S15
**a** (D37, 51 years old)..

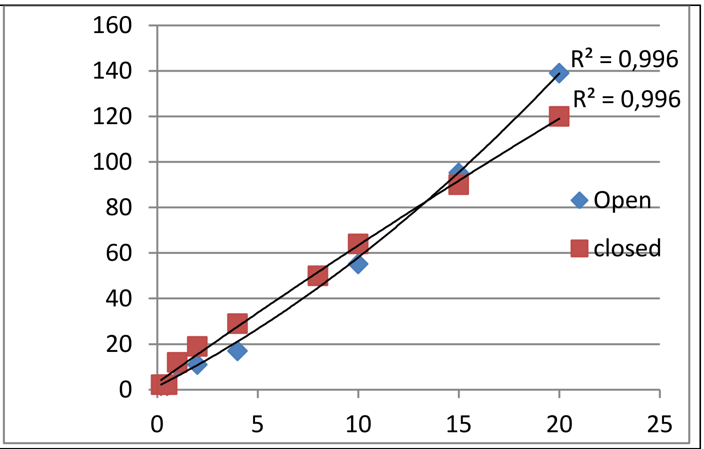

Figure S16 (Ord. Volts)
**b** (D39, 17 years old)

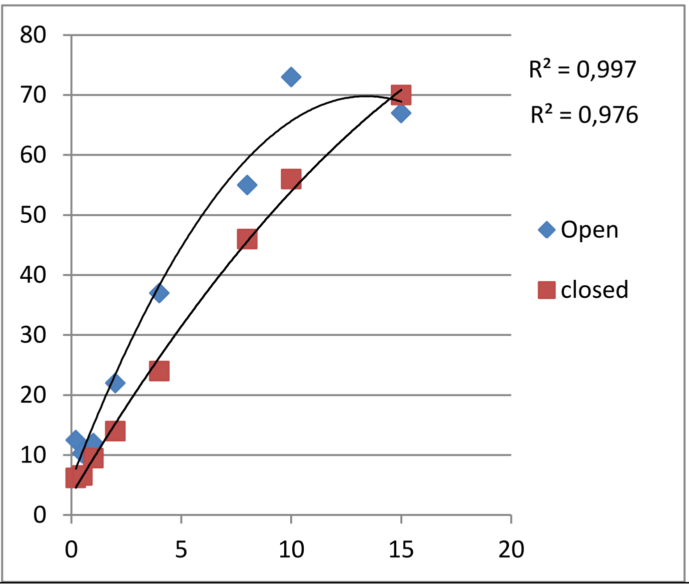

Figure S17
**c** (D44, 25 years old)

The three figures (S10, S11 and S12) show that the mastoid potential with the ear plugged was very close to the mastoid potentials of the same ear when not plugged. Furthermore, regarding the relation between the with earplug and without earplug representative lines, there are three configurations.
In the case of D37 (fig. S10), the magnitude of the "mastoid potential" with airplug (earplug “quies”, pure wax 27 dB) is above that of the without airplug situation. In the case of D39 (fig. S11), the magnitude of the "mastoid potential" "with" airplug is intertwined with that of the "without" situation. In the case of D44 (fig. S12), the magnitude of the mastoid potential "with airplug" is below that of the "without" situation.

However, when the ear is clogged, the signal that reaches the cochlea is necessarily reduced (standard theory) and therefore the cochlear potential should be weakened as well, in all cases! Thus, as logical consequence, the mastoid potential we recorded cannot depend solely upon the cochlear microphonic. Moreover, our measures are consistent with the hypothesis that the mastoid potential depends not only on the impact of sound on the eardrum, but also on the mastoid reaction. So we think that the 'mastoid potential' results, at least in part, from the electrical signal generated by the piezoelectricity of collagenous fibres, located between the points of application of the electrodes (mastoid and forehead). According to our theory linked to the standard interpretation, this piezoelectric response is added to the electric signal from the cochlea.

**Si 05Ec Experiment using 'active noise cancelling headphones' plus ‘custom moulded earplugs' (formed in place)’:**

However, on the three cases above, the occlusion of the external auditory canal, by means of a standard over-the-counter earplug (“quies”, pure wax 27 dB) was, as such, limited to a maximum of 27 dB HL. So, the effects obtained with "earplugs" should be even more marked with “custom-made earplugs (fig. S13) + active noise cancelling headphones (fig. S14 & fig. S15)”. We, therefore, repeated the previous experiments, using active noise cancelling headphones in the context of the situation 'occluded ear'. We filled the external auditory canals of one subject with custom-made earplugs (Audial, Toulouse, Fig. S13), completed by active noise cancelling headphones, (Panasonic RP - HC 500; Fig. S14); Together : fig. S15.

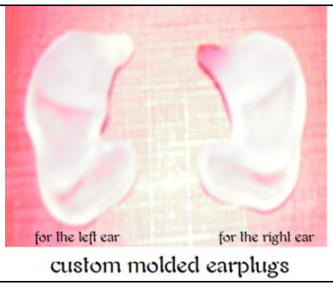

Figure S18
custom-made earplugs

We measured the audiometric thresholds using only custom-made earplugs, then using the active noise cancelling headphones alone (fig. S14), then using both (fig. S15). We used the same placement of the electrodes as in the previous experiments. The previously positioned electrodes, held in place by surgical tape, are pressed by the subject himself, the potential measured depending on the pressure on the skin. The Yamaha HS5 speaker is installed at 1 m from the ear of the subject, its centre facing the subject at the height of the ear to be tested. We performed a calibration of acoustic amplitude with a *Bruel and Kjaer 2231* sound level meter, the rod bearing the microphone being perpendicular to the face of the speaker. The voltage of the output of the Lock-in, plugged into the speaker is set so that the sound level meter displays about 80 dB.

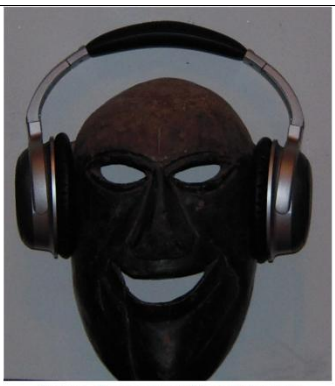

Noise Cancelling Headphones
Panasonic RP-HC 500

Figure S19

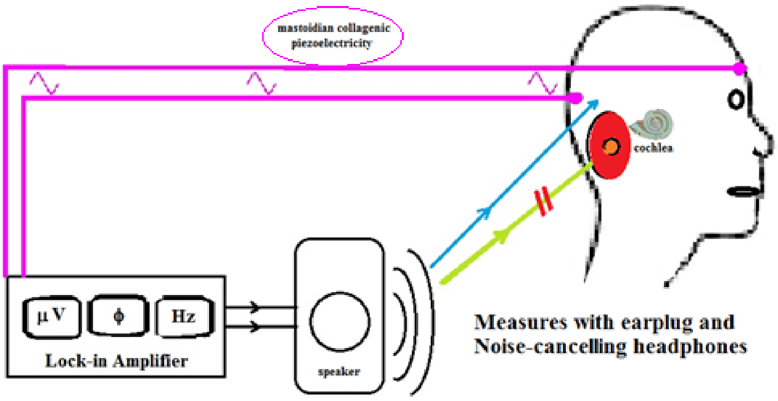

Figure S20

We present below (subject D38, Table S03, Fig. S16), the voltage output of the Lock-in amplifier and measurements of the synchronous electrical potentials, with (Plugged: P) and without (Unplugged: U) occlusive equipment. The data are displayed according to the sound frequency (f kHz) emitted by the speaker, either toward the right auditory canal (REAC) or toward the left one (LEAC).

| f (kHz) | Output voltage (to the speaker) | Vs (µV) U-REAC | Vs (µV) P-REAC | Vs (µV) U-LEAC | Vs (µV) P-LEAC |
|---|---|---|---|---|---|
| 0.5 | 0.4 V | 5 | 5 | 6 | 6 |
| 1 | 0.3 V | 6 | 7 | 5 | 7 |
| 2 | 0.5 V | 6 | 5 | 5 | 6 |
| 4 | 0.2 V | 9 | 7 | 8 | 7 |
| 8 | 0.2 V | 18 | 15 | 13 | 14 |
| 16 | 0.4 V | 44 | 42 | 45 | 43 |
| 20 | 1 V | 123 | 96 | 110 | 105 |
| 32 | 1 V | 144 | 143 | 170 | 145 |

Table S10
External Auditory Canal (EAC); subject D38,
Right (R), Left (L), Unplugged (U), Plugged (P)

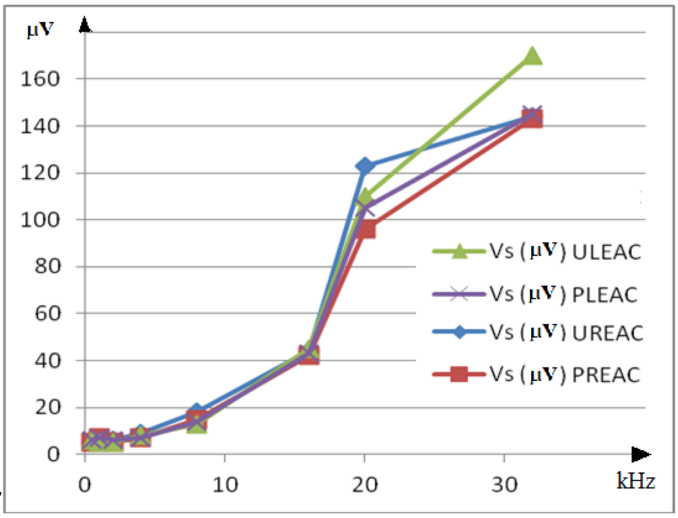

Figure S21

We found that the mastoid potentials with the ear plugged was very close to the mastoid potentials of the same ears when not plugged. As the mastoid potentials are not greatly weakened by plugging the EAC, the classical theory seems to be insufficient and we think that another phenomenon coexists. We hypothesize that this phenomenon is the piezoelectricity of the mastoid collagen. We, therefore, propose the two diagrams below (Fig. S17) in which we have added the effect of the piezoelectricity of the mastoid collagen (blue line) to the classical schema (green line) for purposes of comparison.

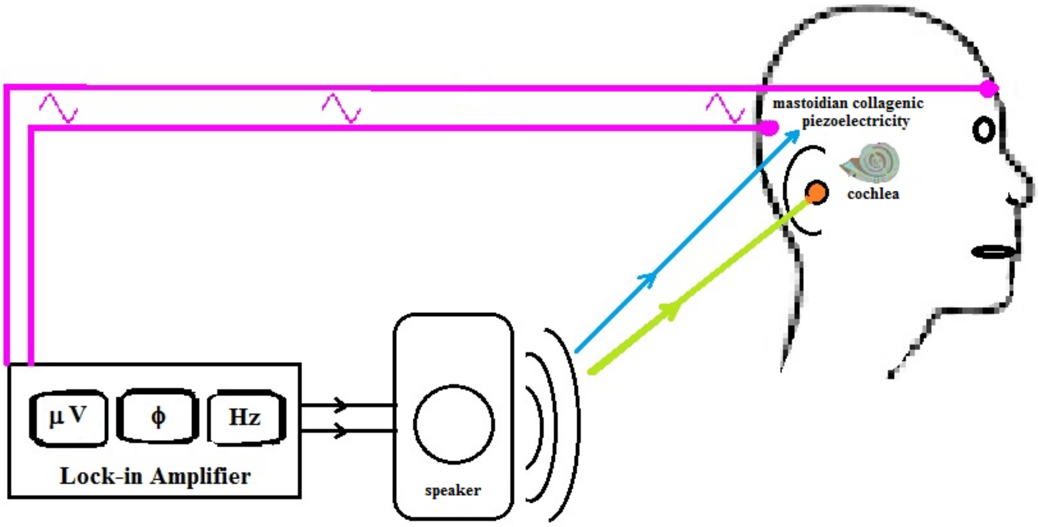

Figure S22
Stimulation restricted to the external auditory conduit (stimulation by headphone only) compared with stimulation targeting the mastoid (stimulation by loudspeaker only)

#### Si 05F Conclusion on Si 05 :

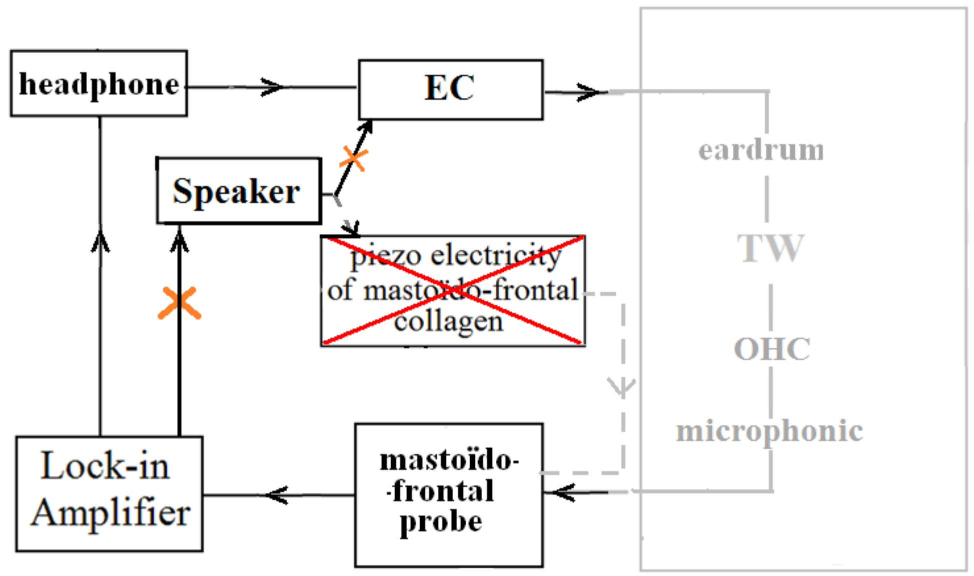

**Figure S23**

Stimulation restricted to the external auditory conduit (stimulation by headphone only) compared with stimulation targeting the mastoid (stimulation by loudspeaker only)

If the mastoid synchronous potential originated from the cochlea alone, then it would be about the same no matter whether the sound was broadcast through a loudspeaker or through the headphone.
If, on the other hand, the mastoid synchronous potential originates locally (at the mastoid), then the sound emitted by the headphone would produce a very low synchronous potential compared to the same sound broadcast through a loudspeaker.

---

[9] Graham, A. , Okabe, M. and Quinlan, R., The role of the endoderm in the development and evolution of the pharyngeal arches. J Anat, 207: 479-487 (2005).
[10] Hill, M.A. Embryology *Hearing - Middle Ear Development*. Cf. https://embryology.med.unsw.edu.au/embryology/index.php/Hearing_-_Middle_Ear_Development (2018).
[11] Gelman SR Wood S., Spellacy WN, Abrams RM, Fetal movements in response to sound stimulation, Am J Obstet Gynecol 143: 484-485 (1982).
[12] Dunn, K., Reissland, N., Reid, V.M.,The Functional Fetal Brain: A Systematic Preview of Methodological Factors in Reporting Fetal Visual and Auditory Capacity, Developmental Cognitive Neuroscience, http://dx.doi.org/10.1016/j.dcn.2015.04.002 (2015)
[13] Boistel, R., et al., How minute sooglossid frogs hear without a middle ear. PNAS, 110(38): 15360–15364. https://doi.org/10.1073/PNAS.1302218110. (2013).

The superior ligament of malleus (K), made of collagen I, attaches the head to the tegmen tympani. The SLM may electrically connect the central structure of the eardrum to bone conductive fibres and to the ground (the electric common return). Peripheral ends of the radial collagen fibres of the eardrum are connected to the osteocytic GJs network.

The eardrum consists of four layers (fig. S19): the epidermis (B), two fibrous collagenic layers (AA'), and an internal mucous membrane (C). The three ossicles are the stapes (F) with its muscle (J), the uncus (E), and the malleus which consists of three parts: the head (D), the lateral process (H) and the manubrium (I). The malleus can be mobilized by its muscle, the tensor tympani (G) which, when enabled, pulls the malleus medially, tensing the eardrum and damping its vibration.

<table>
<tr><td>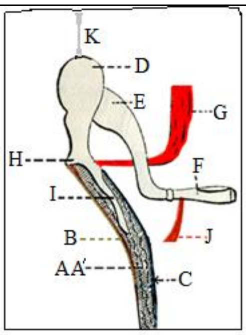
</td><td>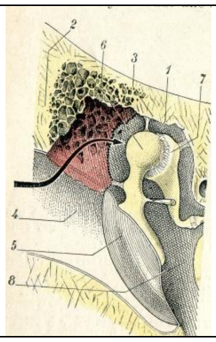
</td></tr>
<tr><td>Figure S24<br>Section of the Eardrum through the handle of the malleus[S153]: Schema of the Superior Ligament of the Malleus (SLM) ensuring the connection of the umbo to the electric common return (K).</td><td>Figure S25<br>Ligaments of the Malleus are inserted into the attic; The Superior Ligament of Malleus (SL M(3)), is inserted in the tegmen tympani[S154].</td></tr>
</table>

The superior ligament of malleus (K), made of collagen I, attaches the head to the tegmen tympani. The SLM may electrically connect the central structure of the eardrum to bone conductive fibres and to the ground (the electric common return). Peripheral ends of the radial collagenic fibres of the eardrum are connected to the osteocytic GJs network (fig. S19 and S20).

##### Si 09B The DOHC complex

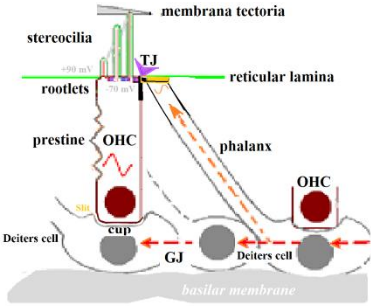

Figure S26
Schema of the DOHC complex
Stereocilia are implanted on the apex membrane of OHCs (cuticular plate / reticular lamina).
They are bathed in endolymph and mobilized by its movements (TW).

#### Si 10 Trickystor

##### Si 10A Source : Ionic channels and time constant

The question of the reality of the contractions at high frequencies of the walls of the OHC in physiological context is poorly understood;the cycle polarization - depolarization created by the movement of the ciliary tuft, induced by the sound, in fact involves the electric plasma membrane time constant; the value of this time constant is not compatible with frequencies of contraction up to 20 kHz in humans and 150 kHz in some bats[S155]. A persistent difficulty in understanding the role of electromotility in amplification is the membrane time constant; the resistance and capacitance of the membrane of an OHC cuticle would suggest that a voltage-driven process should diminish in sensitivity above a corner frequency of only 1 kHz[S156 - S157].
But, the stereociliar signal can trigger prestin electromotility at higher frequencies, provided that it is combined with a second stimulation, whose effects would not be subject to the membrane time constant[S158 - S159].
In reality, such a potential is recordable in the gap (slit) separating the DC cup from the base of the OHC[S160-S161 - S162 - S163 -S164 - S165 - S166 - S167] (Cf. Fig. S21, DOHC5: « cup »). For high frequencies (> 8 kHz), variations of potential of the cup are rapidly transmitted to the gap by means of hemi-channels made of Cx26 and Cx30[S168 - S169] ; This being the case, the prestin could contract at the same frequencies[S170 - S171]. It is also possible that variations in voltage of the Deiters cup would produce a capacitive effect, through the Deiters membrane, the cupular gap and the membrane of the base of the OHC.
This extracellular cupular signal has been attributed to electrical stereociliar phenomena, and this attribution has

caused it to be confused with the "cochlear microphonic potential ". But if we refer to the prevailing opinion, according to which the microphonic potential is due, for the most part, to the mechano-electrical activity of the prestin, we are faced with a vicious circle: The contractions/extensions of the prestin would produce the microphonic and this microphonic, then, is invoked to explain the phenomena of contraction / extension of the prestin!

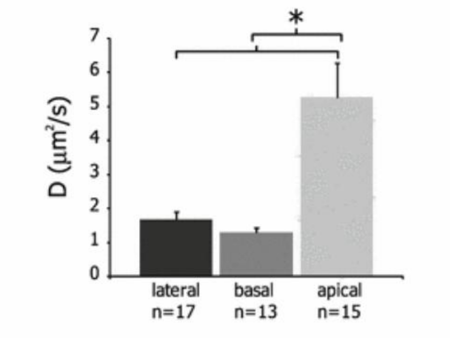

Effective diffusion coefficients, D (µm²/s), across the three OHC regions : D values in the apical region were significantly larger ($p<0.0005$) than in lateral or basal regions. The "membrane fluidity" in the membrane plane is all the more important that the fatty acids of membrane lipids are unsaturated. The fluidity of the cuticular membrane of the OHC is an indication that it contains many more PUFAs than the other parts of the OHC's cellular membrane

Figure S27 (Excerpted from Haroun [S177])

.

#### Si 10D Semi-Conductors of the TkS

The hearing is impaired by genetic disorders of the conjugated and homoconjugated (fig. S24 below) PUFAs semiconductors (Role of peroxisome)[S178]. Peroxisomes are organelles found in all eukaryotic cells. There are at least 32 known peroxisomal proteins that participate in the process of peroxisome assembly. Their deficiency gives impairment hearing of high frequencies and they are involved in catabolism of very long chain fatty acids, branched chain fatty acids, D-amino acids, and polyamines, reduction of reactive oxygen species – specifically hydrogen peroxide – and biosynthesis of plasmalogens, i.e. ether phospholipids.
Peroxin (or peroxisomal/peroxisome biogenesis factor) is a protein found in peroxisomes.

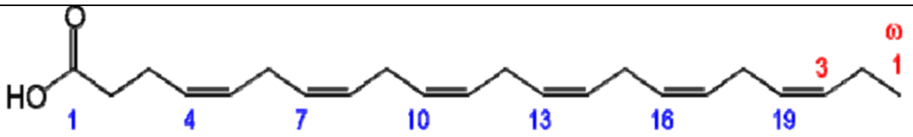

Figure S28
DHA is an example of long chain homoconjugated fatty acid.

#### Si 10E The electrical as well as mechanical state of DCs drives the electromotility of the OHCs

The separation of a DC phalanx from a related OHC cuticle, or of a cupule from the OHC that it supports (or the elimination of GJs between the bodies of Deiters cells) results in deterioration of the electrical behavior of the concerned OHCs (Yu & Zhao, 2009). There being no electrical synapse (GJ) between DCs and OHCs, it has been proposed that this electric effect on the OHC is due not to an electrical interruption between DCs and OHCs, but rather to an interruption of a purely mechanical nature.
Stimulation of the DCs, either electric or mechanical, can modulate the electromotility of the OHCs (Yu&Zhao, 2009) and in vivo active cochlear amplification depends on GJs between the DCs. Cx26 expression in the cochlear supporting cells plays a critical role in active cochlear amplification and its targeted deletion can eliminate active cochlear amplification.
It is coincident with a large reduction in distortion product otoacoustic emission (DPOAE) and severe hearing loss at high frequencies (changes are greater in the shortest OHCs).

#### Si 10F Gate ; Field effect of Deiters Cells on the cuticles of OHCs

The **tunnel field-effect** transistor (TFET) is an experimental type of transistor. Even though its structure is very similar to a metal-oxide-semiconductor field-effect (MOSFET), the fundamental switching mechanism differs, making this device a promising candidate for low power electronics. TFETs switch by modulating quantum tunneling through a barrier instead of modulating thermionic emission over a barrier as in traditional MOSFETs. Because of this, TFETs are not limited by the thermal Maxwell–Boltzmann tail of carriers, which limits MOSFET drain current subthreshold swing to about 60 mV/decade of current at room temperature (exactly 63 mV/decade at 300 K[1]). The concept was proposed by Chang et al while working at IBM [2]. Joerg Appenzeller and his colleagues at IBM were the first to demonstrate that current swings below the MOSFET's 60-mV-per-decade limit were possible.

According to Yu and Zhao (2009), the electric effect of DCs on an OHC must be assigned to an intermediate phenomenon of a mechanical nature. Yet there is an electrostatic link (a field effect) between the cytoskeleton apex of the DCs phalanxes and the OHC cuticle. This link exists without a significant current, and so without electrical

conduction by gap-junctions. That *ephaptic effect*[16] is the field effect of a biological electric conductor, for example the field effect of an axon on its neighbourhood. It is distinguished from the phenomena of simple diffusion in the extra-cellular environment; It is also distinguished from communications between neurons by means of synapses either chemical or electrical [S179, S180, S181].
It is clear that this field effect must be detected mainly by the effects it produces within the cuticular bilayer, where it likely acts as a gate on the communication between the set source {stereocilia / endolymph} and the set drain {OHC cytoplasm, prestin, baso-cellular membrane, cupular slot}.
Indeed, the field effects in the DOHC suggest that it could operate in a way similar to a triode[S182]. A Field Effect Transistor (FET) uses an electric field applied by a gate, to control the conductivity of a "channel" in semiconductor materials. Applied voltage on the gate electrode controls the amount of charge carriers flowing through the system.
In physiological conditions, lipids do not cross TJs: There is a strict insulation, chemical as well as electrical, between the two bound cells, and no current will flow from one to the other of their apical membranes[S183]. For this reason, no electric current can cross the barrier between the phalangeal apexes of the DCs and the cuticle of the embedded OHC. However, this border cannot prevent a hydrophobic intercellular electrostatic coupling unrelated to any GJ[S184 - S185].

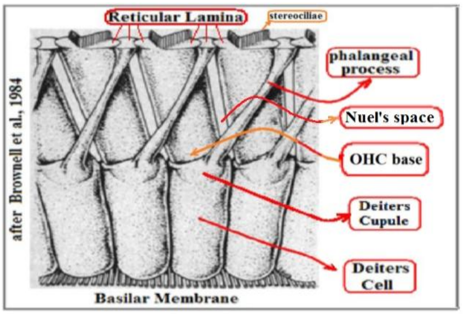

Fig. S29 After Brownell (1984)

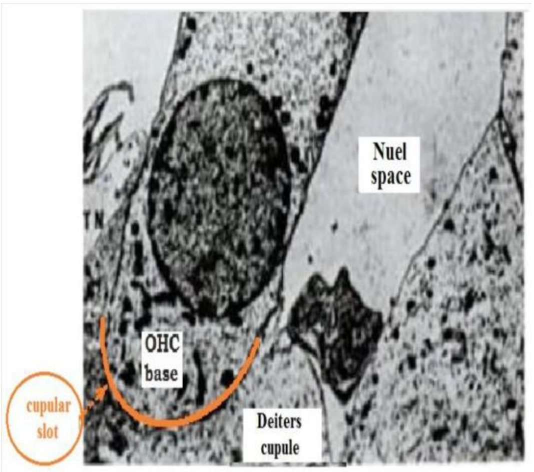

Fig. S30 Adapted from Coujard R., Poirier J., Précis d'Histologie humaine p728 (1980)

[16] The capacitance of a cell membrane varies according to the voltage applied to the membrane of a neighbour cell ("ephaptic effect" ). If we apply this model to the DOHC complex, we can suggest that the capacitance of the cuticular bilayer varies in a manner controlled by the voltage of the contiguous phalangeal apexes. In this way, the piezoelectric information, carried from the tympanum to the four phalanxes of the DCs, should cause its capacitance to vary in a synchronous manner.

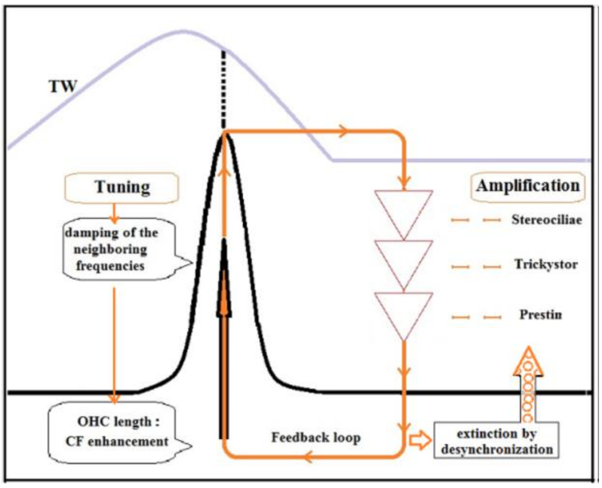

Figure S31
Schematic view of some putative processes regarding
amplification, tuning, damping by DOHC and also creation of OAE.

##### Si 10I About prestin and low frequencies

The conductance of the Mechano Electrical Transducer channels changes are depending upon the tonotopical position within the cochlea, and that suggests differential requirements at different frequencies[S192].
Oddly enough, some profoundly deaf people are able to hear some so-called infra-sound frequencies[S193 - S194] through the labyrinth (sacculus). The cochlea of normally hearing people has mechanisms that weaken or suppress these sound elements before they are transmitted to the brain. Ashmore, points out that “prestin is expressed […] surprisingly, in the cells of the vestibular system[S195]”.
Thus, into its vestibular locus, prestin (otherwise the motor protein of OHCs), might increase or decrease the sound amplitude according to whether the frequency is above or below 50Hz[S17].
It is possible that this "suppressive" effect would be produced by a phase inversion during the amplification process of these infra-sound frequencies, thus producing an attenuation of these stimuli either when they are not useful, or would become disruptive.

***cetaceans*** $\mathbf{f_{QUIX} = f_{QUIX} \pm \Delta f = (60 \pm 20)}$ **kHz**

#### Si 12 OAE versus BEW

The existence of OAEs (Oto Acoustic Emissions) has led to the postulation of a "Backward Travelling Wave" (BTW). Until now, however, no clear evidence has been found for it [S199].
The process of the Electrically Evoked Otoacoustic Emissions (EEOAE), includes electrical stimulation of the OHCs. The effect of this cochlear stimulation is extremely rapid and its upper cutoff frequency is > 80 kHz[S200, S201]. This permits the recording of an acoustic response of the same frequency at the level of the external auditory conduit[S202].

##### Si 14Bb Achilles tendon

| Vsp=2.5V | Subject D39 |
|---|---|
| f (kHz) | V (µV) |
| .2 | 3.2 |
| .5 | 2.6 |
| 1 | 4.0 |
| 2 | 4.7 |
| 4 | 7.1 |
| 8 | 11.9 |
| 10 | 14.8 |
| 12 | 15.9 |
| 15 | 19.7 |
| 20 | 29.8 |

Table S11

**Achilles' tendon**

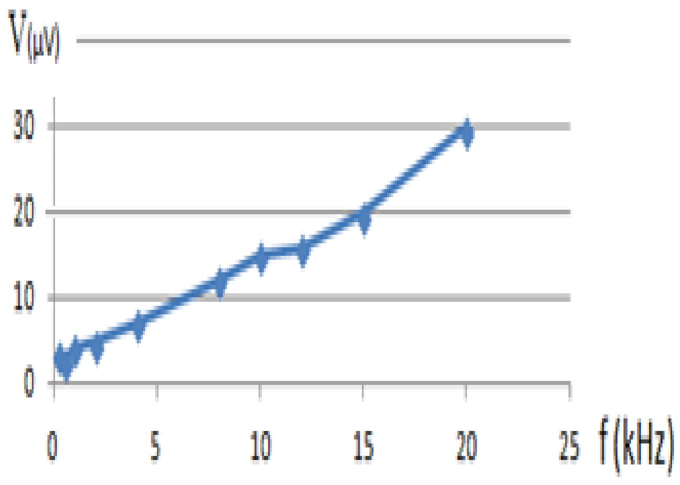

Figure S28

We used the lock-in amplifier to measure synchronous potentials V (µV) of the Achilles tendon in response to the acoustic stimulation of frequency f (kHz), emitted by a Harman/Kardon Loudspeaker: DP/N 0865DV. It was stimulated by a potential Vsp of 2.5 V, according to an installation modelled on that which we have described for the fibres of the eardrum (Table S11).

##### Si 14Bc Knees

As was the case for the Achilles tendon, we used a lock-in amplifier to measure synchronous potentials V (µV) of the patellar tendon in response to an acoustic stimulation of frequency f (kHz), emitted by a Harman/Kardon LoudSpeaker: DP/N 0865DV stimulated by the VLS (Volts) potential. The magnitudes of the synchronous potentials (µV) recorded on the patellar tendon, in function of the acoustic frequency, are reported below for different subjects (Table S12 and Fig. S29).

| f (Hz) | D36 | D37 | D38R | D38L | D39 | Total |
|---|---|---|---|---|---|---|
| 100 | 14 | 5 | 8 | 8 | 8,5 | 43.5 |
| 500 | 12,5 | 4,2 | 6 | 6 | 6,3 | 35.0 |
| 1000 | 15,5 | 5,8 | 7 | 6,1 | 8,5 | 42,9 |
| 2000 | 21 | 9,1 | 8,4 | 7,6 | 10,4 | 56,5 |
| 4000 | 27 | 13,4 | 10,9 | 9,8 | 16,5 | 77,6 |
| 8000 | 42 | 23,7 | 17,1 | 14 | 26,7 | 123,5 |
| 10000 | 54 | 28,4 | 20,5 | 17 | 32,5 | 152,4 |
| 15000 | 73 | 42 | 27,2 | 22 | 47 | 211,2 |
| 20000 | 92 | 61,4 | 35 | 29 | 62 | 279,4 |
| Total | 351 | 193 | 140,1 | 119,5 | 218,4 | 1022 |
| age | 30 | 51 | 73 | 73 | 17 | |

Table S12

Speaker LVL 2.5 V ; F (Hz) ; responses (µV)

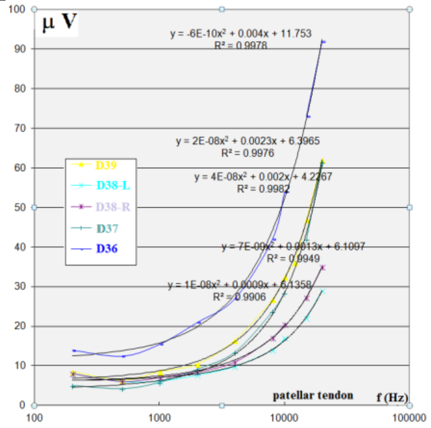

Figure S29

VLS = 2.5 V. Synchronous voltage vs acoustic frequencies for 5 patellar tendons. Every $R^2 > 0.99$

Every musculo-tendinous formation that we tested (mastoid, arm, Achilles tendon, patellar ligament) produced electric potentials synchronous to the acoustic stimuli. The piezoelectric effect sheds some light on this phenomenon.

#### Si 15 Effect of pathological history on measures

Regarding eardrums, we noted some phenomena related to pathological histories;
For example, weakness of piezoelectric responses in the case of allergy inside the ear canal (B16, B17). Anomaly of measurements carried out on a scar zone (D31 on the left eardrum).

D33 uses hearing aids "PS60 Resound" on both ears. Measures were made without any of the two prostheses present. However, the electrical response is particularly high on the eardrum G of this subject. In this case, would the histological structure of the eardrum had evolved by eliciting a more important piezoelectric response, so that the deafness would be more or less compensated ?

#### Si 16 Pain felt during the measurement taken on the eardrum

Regarding the 79 eardrum series, an index of "*felt pain*" (0 to 5) was recorded. 51 recordings describe no pain (0) and 12 a gene or major pain (between 3 and 5). Sometimes the placement of the probe was too painful to result in a measure concerning the ear {A07 / D34}.

### Si 17 Explaining fixed effects by characteristic variables of the individuals

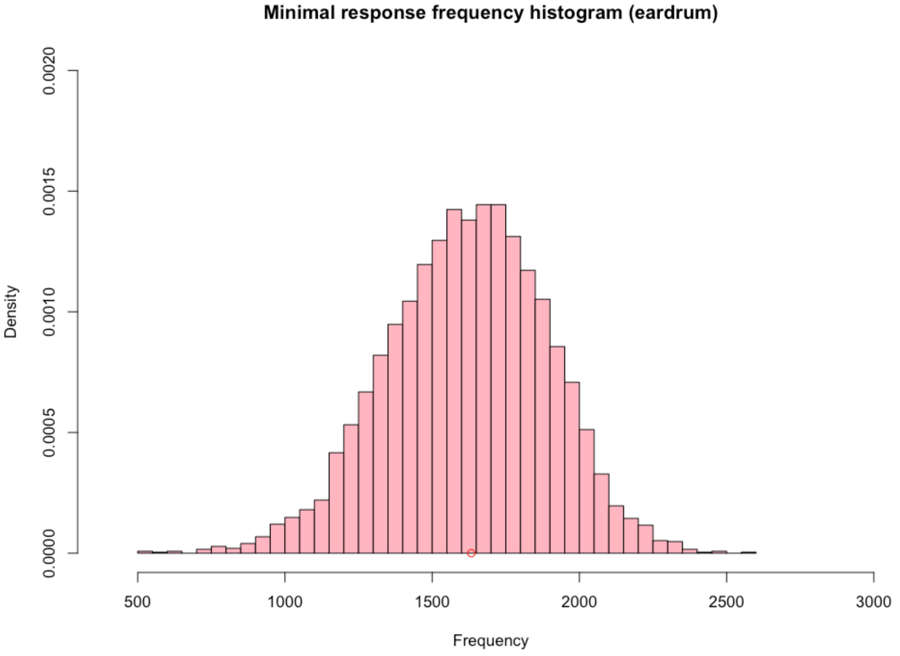

Figure S30
Distribution of the minimal frequency value for eardrum data.

For eardrums the minimal frequency (Quix frequency small red circle in Fig.S30) is lower than the knees one : It appears that separate estimates of the log-quadratic model in the frequencies F for the two sound levels 70 and 80 dB lead to two distinct minimal values 2791Hz and 3375Hz (Fig. S31). Contrary to what we observe for eardrums, it seems that there is for knees a significant interaction effect between the sound frequency and the sound level.

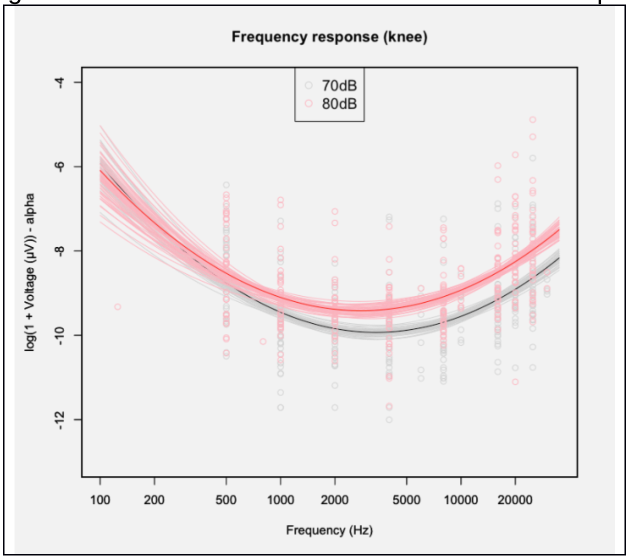

Fig. S31 Effect of the amplitude of the sound stimulus for knee measurements

### Si 18 Comparison of the Wegel curve of the acoustic pressures, versus the curve of the piezoelectric tensions (pT)

For all the individuals, and for the two sides, the minimum value of the curve is obtained for a frequency equal to about $f_{pTmin}$ =1632 hz. The frequency $f_{QUIX} \approx$ 3000 hz (fig. S32)would be the frequency which marks the region of maximum audibility, i.e. for which there is the lowest threshold threshold[s204 - s205] (hearing *'yellow spot'* of Quix). These two minima (1632 hz and 3000 hz) are relatively different. Yet we may see that $f_{QUIX}$ (3000 hz) ≈ 2 $f_{pTmin}$ (1632 hz); ie $f_{QUIX}$ is approximately the harmonic of $f_{pTmin}$..

$$\mathbf{f}_{\mathrm{QUIX}} \approx 2\ \mathbf{f}_{\mathbf{pTmin}}$$

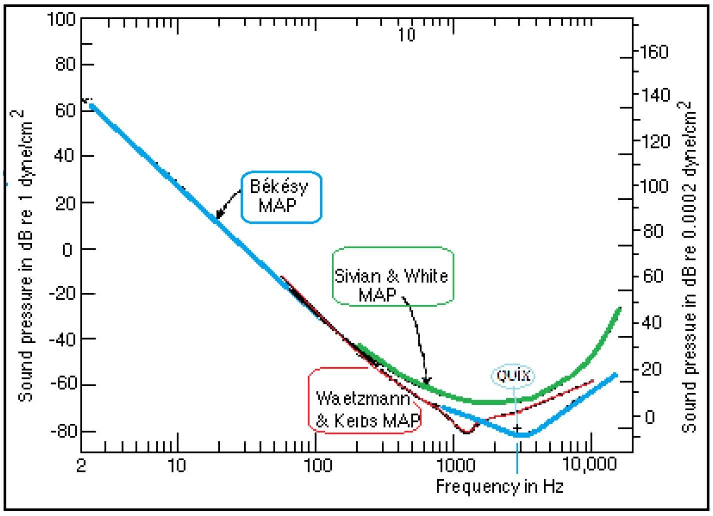

Figure S32
Wegel diagram and other classical schemas excerpted and modified from Licklider, Handbook of Experimental Psychology, S.S. Stevens, ed., 1951

The frequency of Quix seems to be less fixed than its author thought. The frequency of Quix is that which is audible by the lowest acoustic stimulation compared to the other frequencies. In our experiments, the stimulation of the eardrums used relatively high amplitudes in relation to the threshold of hearing. As a result, the frequency of the minimum of the pT curve is lower than the frequency of Quix, but the two frequencies may be closer, or coincide, if we use liminal acoustic stimuli, in both cases (acoustic and piezo-electric).

### Si 21 Supplementary Information Bibliography

[109]Gavilan C., Sanjuàn J., Microphonic Potential picked up from the human tympanic membrane. *Ann Otol Rhinol Laryngol*. **73,** 102-109 (1964).
[110]Auriol B., Béard J., Broto J.M., Descouens D., Procédé et dispositif de mesure d'une tension électrique relative à une fibre de collagène pour l'aide au diagnostic par un praticien en vue de l'identification d'une éventuelle altération et de l'évaluation de la qualité fonctionnelle de la fibre de collagène", *French patent application* n° FR11/54672 (2011).
[111]Tasaki I., Davis H., Legouix J., The space time pattern of the cochlear microphonics (guinea pig) as recorded by differential electrodes, JASA **24**, 502-18 (1952).
[112]Offutt G., *The electromodel of the auditory system*, Golo Press, 23-24 (1984).
[113]Mammano F., oral discussion at the Colloquium of Sylvanès 2012
[114]Denning D. et al., The piezoelectric tensor of collagen fibrils determined at the nanoscale, *ACS Biomater Sci Eng*. Publication Web, DOI: 10.1021/acsbiomaterials.7b00183 ( 2017).
[115]R.D. Sinelnikov, Atlas of the anatomy of man, Study on the bones, joints, ligaments and muscles (updated & augmented, 1963, I: 232). Р. Д. СИНЕЛЬНИКОВ, АТЛАС АНАТОМИИ ЦЕLOВЕКА (СССР 1963) I: 232; [R.D. Sinelnikov, Atlas Anatomie Celoveka, v trekh tomakh]
[116]Denning D. et al., The piezoelectric tensor of collagen fibrils determined at the nanoscale, *ACS Biomater. Sci. Eng;* DOI:10.1021/acsbiomaterials.7b00183 ( 2017).
[117]Aron M., Floyd D., and Bance M., Voluntary eardrum movement: a marker for Tensor Tympani contraction ? *Otol Neurotol* **36** 1-9 (2014).
[118]Bance M. et al., Effects of Tensor Tympani muscle contraction on the middle ear and markers of a contracted muscle, *Laryngoscope* **123,** 1021–27 (2013).
[119]Henson M.M., Madden V.J., RaskAndersen H., Henson Jr O.W.., Smooth muscle in the annulus fibrosus of the tympanic membrane in bats, rodents, insectivores, and humans, *Hear Res* **200,** 2937 (2005).
[120]Puria S, Fay RR, Popper AN, Smooth muscle in the annulus fibrosus of the tympanic membrane in bats, rodents, insectivores, and humans (2013).
[121]https://upload.wikimedia.org/wikipedia/commons/thumb/3/32/Ground_loop.svg/640px-Ground_loop.svg.png
[122]http://www.thinksrs.com/downloads/PDFs/ApplicationNotes/AboutLIAs.pdf
[123]Gavilan C., Sanjuàn J., Microphonic Potential picked up from the human tympanic membrane, *Ann Otol Rhinol Laryngol* **73,** 102-9 (1964).
[124]Davis H., Derbyshire A.J., Lurie M.H. and Saul L.J., The electric Response of the Cochlea, *Am. J. of Ph*. **107**, 311-32 (1934).
[125]Reyes S., Ding Da., Sun W., Salvi R., Effect of Inner and Outer Hair cell lesions on electrically evoked otoacoustic emissions, *Hear Res* **158,** 139-50 (2001).
[126]Wever, E.G. and C.W. Bray, Auditory nerve impulses, *Science* **71,** 215 (1930).
[127]Trautwein P., Hofstetter P., Wang J., Salvi R. and Nostrant A., Selective inner hair cell loss does not alter DPOAEs, *Hear. Res*. **96,** 71-82 (1996).
[128]Gavilan C. and Sanjuàn J., Microphonic Potential picked up from the human tympanic membrane, *Ann Otol Rhinol Laryngol* **73,**102-9 (1964).
[129]Carricondo F, GilLoyzaga P, SanjuànJuaristi J, PochBroto J., Cochlear microphonic potentials: a new recording technique. *Ann Otol Rhinol Laryngol* **110**, 56573 (2001).
[130]Carricondo F, GilLoyzaga P, SanjuànJuaristi J, PochBroto J., Cochlear microphonic potentials: a new recording technique. *Ann Otol Rhinol Laryngol* **110**, 56573 (2001).
[131]PochBroto J. et al., Cochlear microphonic audiometry: a new hearing test for objective diagnosis of deafness, *Otol Rhinol Laryngol* **129,** 74954 (2009).
[132]Mammano F., personal discussion [Colloque de Sylvanès 2012 (18-20 Mai 2012)]
[133]Carricondo F et al., (2001).
[134]PochBroto J. et al. (2009).
[135]Behari J. *Biophysical Bone Behaviour: Principles and Applications*, Wiley - Science (2009).
[136]Wada H., Ando M., Takeuchi M., Sugawara H., Koike T., Vibration measurement of the tympanic membrane of guinea pig temporal bones using time-averaged speckle pattern interferometry, JASA **111,** 2189-99 (2002).
[137]Okayasua T. Human ultrasonic hearing is induced by a direct ultrasonic stimulation of the cochlea, *Neurosci Lett* **539**, 71–6 (2013).
[138]Humbel R.L. Auto-anticorps dans les maladies de l'oreille interne, 4° Colloque GEAI (2006)
[139]ISO, Acoustics, Hearing protectors, Part 1: Subjective method for the measurement of sound attenuation (1990).
[140]Margolis R.H., The Vanishing Air-Bone Gap - Audiology's Dirty Little Secret, *Audiol Online* (2008).
[141]Stenfelt, S.& Reinfeldt, S., A model of the occlusion effect with bone-conducted stimulation. *International J Audiology* **46**, 595-608 (2007).
[142] Denning D. et al., The piezoelectric tensor of collagen fibrils determined at the nanoscale, *ACS Biomater. Sci. Eng;* DOI:10.1021/acsbiomaterials.7b00183 ( 2017).
[143]Nachtigall PE, Lemonds DW, and Roitblat HL, Psychoacoustic of Dolphin and Whale Hearing, p.336-9 in Au WWL, Popper AN, and Fay RR, Hearing by Whales and Dolphins, Springer NY (ISBN 0-387-94906-2) (2000).
[144]Dolphin W.F. pp.322-325.
[145]Bullock et al., Electrophysiological studies of central auditory mechanisms in cetaceans, *Zeitschrift Vergleichen Physiol* **59**, 117-56 (1968).
[146]Fraser FC and Purves PE, A Discussion on the 'Ear' Under Water, *Proc R Soc Lond B Biol Sci*, **152,** 62-77, (1960).
[147]Ridgway SH The Auditory Central Nervous System of Dolphins. In: Au WWL. Fay RR, Popper AN (eds) Hearing by Whales and Dolphins, *Springer Handbook of Auditory Research* **12**. Springer, New York, NY (2000).
Ridgway SH The Auditory Central Nervous System of Dolphins. In: Au WWL, Fay RR, Popper AN (eds) Hearing by Whales and Dolphins, Springer Handbook of Auditory Research, vol.**12**, Springer, New York (2000).
[148] Yamato, M., Ketten, D. R., Arruda, J., Cramer, S. and Moore, K. (2012), The Auditory Anatomy of the Minke Whale (*Balaenoptera acutoros-trata*): A Potential Fatty Sound Reception Pathway in a Baleen Whale. Anat Rec, 295: 991–998. doi:10.1002/ar.22459

[149]Cranford TW, Krysl P, Amundin M (2010) A New Acoustic Portal into the Odontocete Ear and Vibrational Analysis of the Tympanoperiotic Complex. PLoS ONE5(8): e11927. https://doi.org/10.1371/journal.pone.0011927
[150]Cranford TW, Krysl P, Amundin M, A new acoustic portal into the odontocete ear and vibrational analysis of the tympanoperiotic complex, *PLoS ONE* **5**:e11927 (2010).
[151]Yamato MK et al., *Anat Rec*, **295,** 991-8 (2012).
[152]Alberts B. et al., *Molecular biology of the cell,* 5°ed., Garland Science, NY, pp. 1474-6, ISBN 978-0-8153-4106-2 (2008).
[153] Tillaux P.J., *Traité d'Anatomie topographique avec applications à la chirurgie*, Paris Asselin et Houzeau, 4°ed. p.116 (1903).
[154] Testut L. et Jacob O., Traité d'Anatomie Topographique, T.1 Fig. 220, p.302, Doin Paris (1905)
[155] Petit C., Cours au Collège de France, 18 et 25 mars, ler et 8 avril 2004, Mécanismes d'analyse des fréquences sonores.
[156] Santos-Sacchi J., On the frequency limit and phase of outer hair cell motility: effects of the membrane filter, J of Neurosc, 12, 1906-16 (1992).
[157] Gale J. E., Ashmore J. F, An intrinsic frequency limit to the cochlear amplifier, *Nature* **389**, 63–6 (1997).
[158] Zha D et al., In Vivo Outer Hair Cell Length Changes Expose the Active Process in the Cochlea. PLoS ONE 7(4): e32757 (2012).
[159]Dallos P, Evans BN, High-frequency motility of outer hair cells and the cochlear amplifier, *Science* **267,** 2006-9 (1995).
[160]Fridberger A, Organ of Corti Potentials and the Motion of the Basilar Membrane, *J. Neurosc*, **24,** 10057-63 (2004).
[161]Spector A.A., Popel A.S., Eatock RA, Brownell WE, Mechanosensitive channels in the lateral wall can enhance the cochlear outer hair cell frequency response. *Ann. Biomed. Eng*. **33**, 991–1002 (2005).
[162]Kaneko T. , Harasztosi C. , Mack AF , Gummer AW., Membrane traffic in outer hair cells of the adult mammalian cochlea, Eur. J. Neurosci **23**, 2712–22, (2006).
[163]Dallos P, Evans BN, High-frequency motility of outer hair cells and the cochlear amplifier, *Science* **267**, 2006-9 (1995).
[164]Ashmore JF., Cochlear outer hair cell motility, *Physiol. Rev* **88**, 173–210 (2008).
[165] Mistrik, P., Mullaley, C., Mammano, F., Ashmore, J., Three-dimensional current flow in a large-scale model of the cochlea and the mechanisms of amplification of sound, *J. Roy. Soc*. Interface **6**, 279–91 (2009).
[166] Mistrík P., and Ashmore JF., Reduced Electromotility of Outer Hair Cells Associated with Connexin-Related Forms of Deafness: An In silico Study of a Cochlear Network Mechanism, JARO **11**, 559–71 (2010).
[167] Ashmore J et al., The remarkable cochlear amplifier, *Hear Res* **266**, 1–17 (2010).
[168]Zhao HB and Yu N, Distinct and gradient distributions of connexin26 and connexin30 in the cochlear sensory epithelium of guinea pigs, *J Comp Neurol* **499,** 506-18 (2006).
[169] Laird DW, The gap junction proteom and its relationship to disease, *Trends Cell Biol,* **20**,92-101, (2010).
[170] Dallos P et al. (1995).
[171]Fridberger A., Loud sound-induced changes in cochlear mechanics. *J Neurophysiol* **88**, 2341- 8 (2002).
[172] Cody AR, Russell IJ, The responses of hair cells in the basal turn of the guinea-pig cochlea to tones. *J Physiol* **383**, 551–69 (1987).
[173] Lagostena L, Cicuttin A, Inda J, Kachar B, Mammano F, Frequency Dependence of Electrical Coupling in Deiters' Cells of the Guinea Pig Cochlea, *Cell Communication & Adhesion,* **8**, 393 –9 (2001).
[174]Harroun TA., Katsaras J, and Wassall SR, Cholesterol is Found to Reside in the Center of a Polyunsaturated Lipid Membrane, Biochemistry, **47**, 7090–6, (2008).
[175]Marrink S. J., de Vries A H, Harroun T A, Katsaras J, and Wassall SR Cholesterol Shows Preference for the Interior of Polyunsaturated Lipid Membranes. *J Am Chem Soc,* **130**, 10–11 (2008).
[176] Nguyen TV, Brownell WE, Contribution of membrane cholesterol to outer hair cell lateral wall stiffness, *Otolaryngol Head Neck Surg* **119**,14–20 (1998).
[177]Harroun TA, Katsaras J, and Wassall SR, Cholesterol is Found to Reside in the Center of a Polyunsaturated Lipid Membrane, *Biochemistry*, **47,** 7090–6 (2008)
[178]Wanders R.J.A., Peroxisomes, lipid metabolism, and peroxisomal disorders, *Mol Genet Metab* **83,** 16-27 (2004).
[179] Arvanitaki A., Effects evoked in an axon by the activity of a contiguous one, *J Neurophysiol* **5**, 89-108 (1942).
[180]Goldwyn JH, Rinzel J, Neuronal coupling by endogenous electric fields: cable theory and applications to coincidence detector neurons in the auditory brain stem, *J Neurophysiol*, **115**, 2033-51 (2016).
[181]Zhang M et al.., Propagation of epileptiform activity can be independent of synaptic transmission, gap junctions, or diffusion and is consistent with electrical field transmission. *J Neurosci* **34**, 1409–1419, 2014.
[182]LilienfeldJ.E.,https://www.google.com/search?tbo=p&tbm=pts&hl=en&q=ininventor:%22Lilienfeld+Julius+Edgar%22 Method and apparatus for controlling electric currents, US 1745175A patent (1930).
[183]Van Itallie Christina M., Fanning Alan S., Bridges Arlene, and Anderson James M., ZO-1 Stabilizes the Tight Junction Solute Barrier through Coupling to the Perijunctional Cytoskeleton, Molecular Biology of the Cell, Vol. 20, 3930–3940, September 1, 2009.
[184] Goldwyn JH and Rinzel J, Neuronal coupling by endogenous electric fields, in http://adsabs.harvard.edu/abs/ arXiv150801741G (2015). [The authors provide a cable theory framework to study how a bundle of model neurons generates $V_e$ and how this $V_e$ feeds back and influences membrane potential ($V_m$)].
[185]Buzsáki G, Anastassiou CA, Koch C. The origin of extracellular fields and currents. Nat Rev Neurosci 13: 407–420, **2012**.
[186] Szarama Katherine B., Núria Gavara, Petralia Ronald S., Kelley Matthew W. and Chadwick Richard S., Cytoskeletal changes in actin and microtubules underlie the developing surface mechanical properties of sensory and supporting cells in the mouse cochlea, Development 139, 2187-2197 (2012) ; doi:10.1242/dev.073734
[187] Sataric, M.V., et al., Role of nonlinear localized Ca2+ pulses along microtubules in tuning the mechano-sensitivity of hair cells, *Prog Biophys Mol Biol* **119**,162–74 (2015).
[188] See: Vater M., Lenoir M., and Pujol R., Developpment of the organ of Corti in Horseshoe Bats: Scanning and Transmission Electron Microscopy, *J Comp Neurol*, **377**, 520-534 (1997).
[189] Mahendrasingam S, Beurg M, Fettiplace R, Hackney CM, The ultrastructural distribution of prestin in outer hair cells: A post-embedding immunogold investigation of low and high frequency regions of the rat cochlea. *Eur J Neurosci,* **31,** 1595–605 (2010).
[190]Priel A, Tuszynski JA, A nonlinear cable-like model of amplified ionic wave propagation along microtubules,. *Eur Phys Lett,* **83**, 68004 (2008).
[191]Zha D et al., In Vivo Outer Hair Cell Length Changes Expose the Active Process in the Cochlea. PLoS ONE 7(4): e32757 (2012).
[192]Ricci, AJ, Crawford AC and Fettiplace R, Tonotopic variation in the conductance of the hair cell mechanotransducer channel, *Neuron* **40**, 983–90 (2003).

[193] Toupet M., Les sons forts peuvent donner le vertige – peut-on entendre par le vestibule ?, *Inst Méd*, **4,** 21-4, (1981).

[194]Sheykholeslami, K, Kaga K, The otolithic organ as a receptor of vestibular hearing revealed by vestibular-evoked myogenic potentials in patients with inner ear anomalies, *Hear Res,* **165**, 62–7 (2002).

[195]Ashmore J, Cochlear outer hair cell motility, *Physiol Rev* **88**, 173-210 (2008).

[196] Rogers PH, Hawkins AD, Popper AN, Fay RR, Gray MD, Parvulescu Revisited, Small Tank Acoustics for Bioacousticians In: Popper A, Hawkins A (eds) The Effects of Noise on Aquatic Life, *Adv Exp Med Biol,* **875**, Springer, NY (2016).

[197]Steinmetz CP, Electric engineering, Theory and Calculation of Transient Electric Phenomena and Oscillations, **8**, McGraw-Hill Book (1920).

[198]Fridberger A., et al., Organ of Corti Potentials and the Motion of the Basilar Membrane, *J Neurosci* **24**, 10057–63 (2004).

[199]Ren T, He W, Scott M and Nuttall AL, Group Delay of Acoustic Emissions in the Ear, *JN Physiol* **96** , 2785-91 (2006).

[200]Grosh K, Zheng J, Zou Y, de Boer E, and Nuttall AL., High-frequency electromotile responses in the cochlea, JASA, **115**, 2178-84 (2004).

[201]Reyes S, Ding D, Sun W, Salvi R, Effect of Inner and Outer Hair cell lesions on electrically evoked otoacoustic emissions, *Hear Res*, **158,** 139-50 (2001).

[202] Reyes S. et al. (2001).

[203] He W.and Rena T., Basilar membrane vibration is not involved in the reverse propagation of otoacoustic emissions, *Sci Rep,* **3,** 1874 (2013).

[204]Quix F. H.. Mil. Geneesk Tydschr (1903).

[205] Zwaardemaker H., Sur la sensibilité de l'oreille aux différentes hauteurs des sons, *L'année psychologique* **10,** 161-78 (1903).

[206] Vasilescu G., Electronic Noise and interfering signals, principles and applications, *Signals and communication technology*, Springer Verlag, ISBN 3-540-40741-3 (2005).

[207] http://www.cnrtl.fr/definition/vicariance

[208] Seldran, F., A model-based analysis of the “combined-stimulation advantage”, *Hear. Res.* **282**, 252-64 (2011).
